# GIRAFFOIDS FROM THE SIWALIKS OF PAKISTAN

**DOI:** 10.1101/2023.10.06.561267

**Authors:** Nikos Solounias, Maria Rios Ibáñez

## Abstract

Although today they occur in Africa, during the Miocene Giraffoids were widespread in Eurasia and Africa. We describe the giraffoid faunas found in the Siwaliks of Pakistan. These faunas are extremely rich in fossil mammals among which there are several new giraffoid taxa. In total, 28 species. They are an addition to taxa known before and give new insights in evolutionary relationships of all the taxa. Non-Siwalik localities that contribute key information are: Fort Ternan, Kalodirr, Tung Gur, Gebel Zelten, Gebel Hamrin and Bou Hanifia. (1) The Palaeomerycidae are distinct with ossicones and an occipital horn. The list of taxa includes five new Palaeomerycidae: *Tauromeryx canteroi* nov. sp., *Nuchalia gratia* nov. gen. sp. nov., *Fovea fossata* nov. gen. sp. nov., *Goniomeryx flynni* gen. nov. sp. nov., and *Lateralia morgani* nov. gen. nov. sp. nov.; (2) new Climacoceridae: *Vittoria soriae* gen nov. sp nov. *Orangemeryx badgleyi* nov. sp. *Pachya moruoroti* gen. nov. sp. nov. *Prolibytherium fusus* has already been described. We also define two new ranks within the Climacoceridae family: the Climacocerinae and the Prolibytheriinae. (3) Preliminary systematics suggest that Giraffidae can be subdivided into two clades: the long and the short neck groups. The Siwaliks sample a total of fourteen different giraffids: *Progiraffa exigua*, *Giraffokeryx punjabiensis*, *Ua pilbeami*, *Orea leptia* new gen and new sp, *Injanatherium hazimi*, *Giraffa punjabiensis, Giraffa sivalensis*, *Decennatherium asiaticum*, *Bramatherium megacephalum*, *Bramatherium perminese*, *Bamiscus micros* new gen and new sp., and *Sivatherium giganteum*. *Palaeotragus germanii* from Bou Hanifia and *Bohlinia tungurensis* from Tung Gur are new determinations. Progiraffinae and Bramatheriinae are new subfamilies of Giraffidae. The subfamilies of Giraffidae seem to hold as isolated evolutionary silos. Thus, they are hard to connect systematically. They are: Progiraffinae, Giraffokerycinae, Okapiinae, Bohlininae, Giraffinae, Palaeotraginae, Samotheriinae, Bramatheriinae and Sivatheriinae. It is hard to connect these subfamilies as they appear distinct around 16 Ma.

## INTRODUCTION

The base of all zoological investigations is new data and the resulting new names. Establishing that identification base is critical for future research in animal diversity and in paleoecology. This study provides such a base which is sometimes based on a single specimen. Ignoring these “singletons” would change our understanding of animal diversity. As long as they contain diagnostic morphology these “singletons” are necessary. The Siwalik formations found in the Indian Subcontinent consist of Miocene to Pleistocene fluvial sediments that were deposited in a sequence of basins located along the southern edge of the collision zone between Asia and India. These sediments are characterized by their thickness and abundance of fossils, including a varied collection of terrestrial and freshwater vertebrates. As a result, they have captured the attention of scientists for more than 170 years (Barry et al., 2013). They are the Zinda Pir and the Kamlial (oldest), Chinji, Nagri, Dhok Pathan and the Plio-Pleistocene. Additional localities are necessary and key to the Siwalik study. The major ones are: Fort Ternan, Moruorot, Kalodirr, Tung Gur, Gebel Zelten and Bou Hanifia. Solounias and Danowitz 2019) provided a preliminary survey of the Giraffoidea a Siwalik volume to be published by The Johns Hopkins University Press. In that survey multiple authors did not name or figure anything new. The current study accomplishes the naming and the figures of the Giraffoidea and supersedes other simpler interpretations.

The Siwalik Giraffoidea are 28 species. They are divided into three different parts: The Lower Siwaliks are the older, and include the Zinda Pir and the Kamlial (oldest), Chinji, formations, which are mainly Lower and Middle Miocene in age. The Kamlial Formation which extends into the Early Miocene; the Middle Siwaliks are mostly Late Miocene in age and are comprised by the Dhok Pathan Formation and the underlying Nagri Formation and is where we found a high abundance of giraffoids. Finally, the Upper Siwaliks correspond to the layers that contain the Plio-Pleistocene levels (Flynn et al., 2013). In the upper layers the giraffoid diversity is low.

The Siwalik giraffids first came to knowledge thanks to the expeditions of Falconer and Cautley during the 1830’s and Pilgrim during the 1900s. Most of the giraffids found were described on fragmentary or/and undiagnostic remains such as *Giraffokeryx punjabiensis* (a few teeth), *Progiraffa exigua* (one tooth), *Giraffa priscilla* (one tooth), *Giraffa punjabiensis* (teeth in Calcutta Indian Museum never figured). *Giraffa sivalensis* and *Sivatherium giganteum* were described on good material (Falconer and Cautley, 1836; Pilgrim, 1908, 1911).

We describe the material collected by the Yale/Harvard Geological Survey of Pakistan during the decades of the 1970s-90s, led by David Pilbeam, John Barry, Michèle Morgan, Larry Flynn, Mahmood Raza, Nikos Solounias and others. These materials have accurate chronological and stratigraphic data that help with the interpretations and show us a highly rich continental faunas in which giraffoids where both abundant and morphologically diverse.

## MATERIAL AND METHODS

Material is temporarily deposited at the Peabody Museum of Harvard – It is a Yale/ Harvard and Geological Survey of Pakistan collection from 1976 to 2011. The figures show the new taxa and a review most key older taxa. The study is a pictorial data base that will be used in the future. Ma is mega-anum – a million years.

## SYSTEMATIC PALEONTOLOGY

Order Artiodactyla Owen 1848

Suborder Ruminantia Scopoli 1777

Infraorder Pecora Linnaeus 1758

Superfamily Giraffoidea Simpson 1931

Pecoran ruminants with ossicone type horns. It includes three families:

Palaeomerycidae, Climacoceridae and Giraffidae. Their ossicones are separate ossifications above the frontal bone. They appear as distinct little cups above the frontal in fetal life. They eventually fuse in older age to the frontal. Ossicones are commonly covered by skin which is sometimes keratinized by the apex (*Giraffa*). Ossicones can be bare at the apex and dead bone there in *Okapia*, *Palaeotragus,* and *Samotherium*. The integument retreats and exposes the apical bone (*Okapia - Ua*). Rarely an ossicone is made from keratin instead of bone which is found in *Okapia*.

### Family Palaeomerycidae Lydekker, 1883

Palaeomerycids are Old World giraffoids that appeared during the early Miocene. They are characterized for the presence of frontal ossicones as in Giraffidae and an additional occipital horn, convergent but not related to the one of the north American Dromomerycidae. The confusion of dromomerycids with palaeomerycids is widespread. The dromomerycids have horns of frontal origin not ossicones. The palaeomerycids have true ossicones. They both share the convergence of an occipital horn. For reference see Prothero and Litter (2007; Astibia and Morales 2006; Sánchez et al. 2015; Costeur et al. 2015). Palaeomerycids commonly have a dental enamel spur termed the *Palaeomeryx* fold.

*Palaeomeryx sp.* A new skull under study from Gansu (Fig 1a).

**Fig. 1a.**
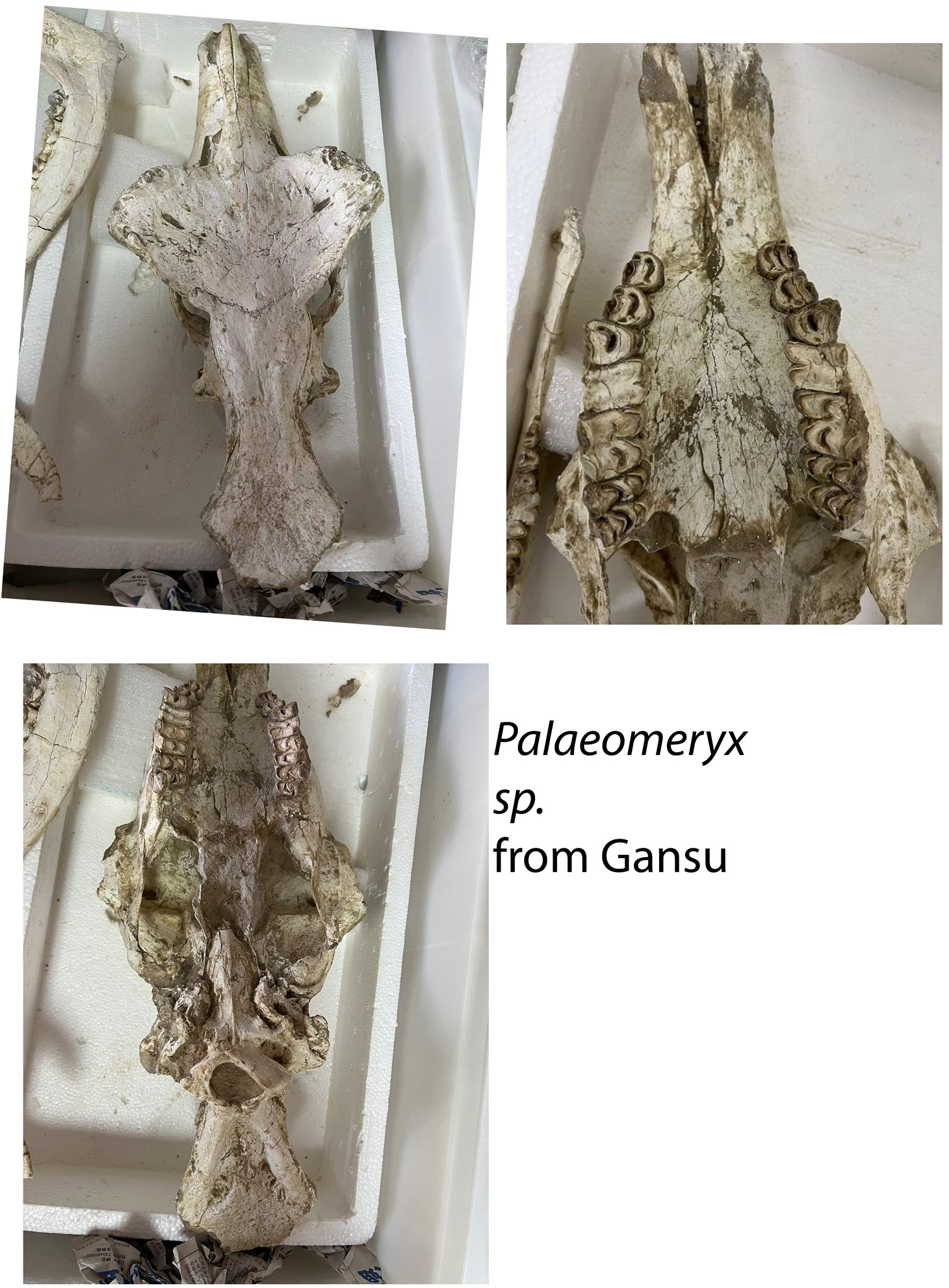
*Palaeomeryx sp.* from Gansu. Specimen under study.

**Fig. 1b.**
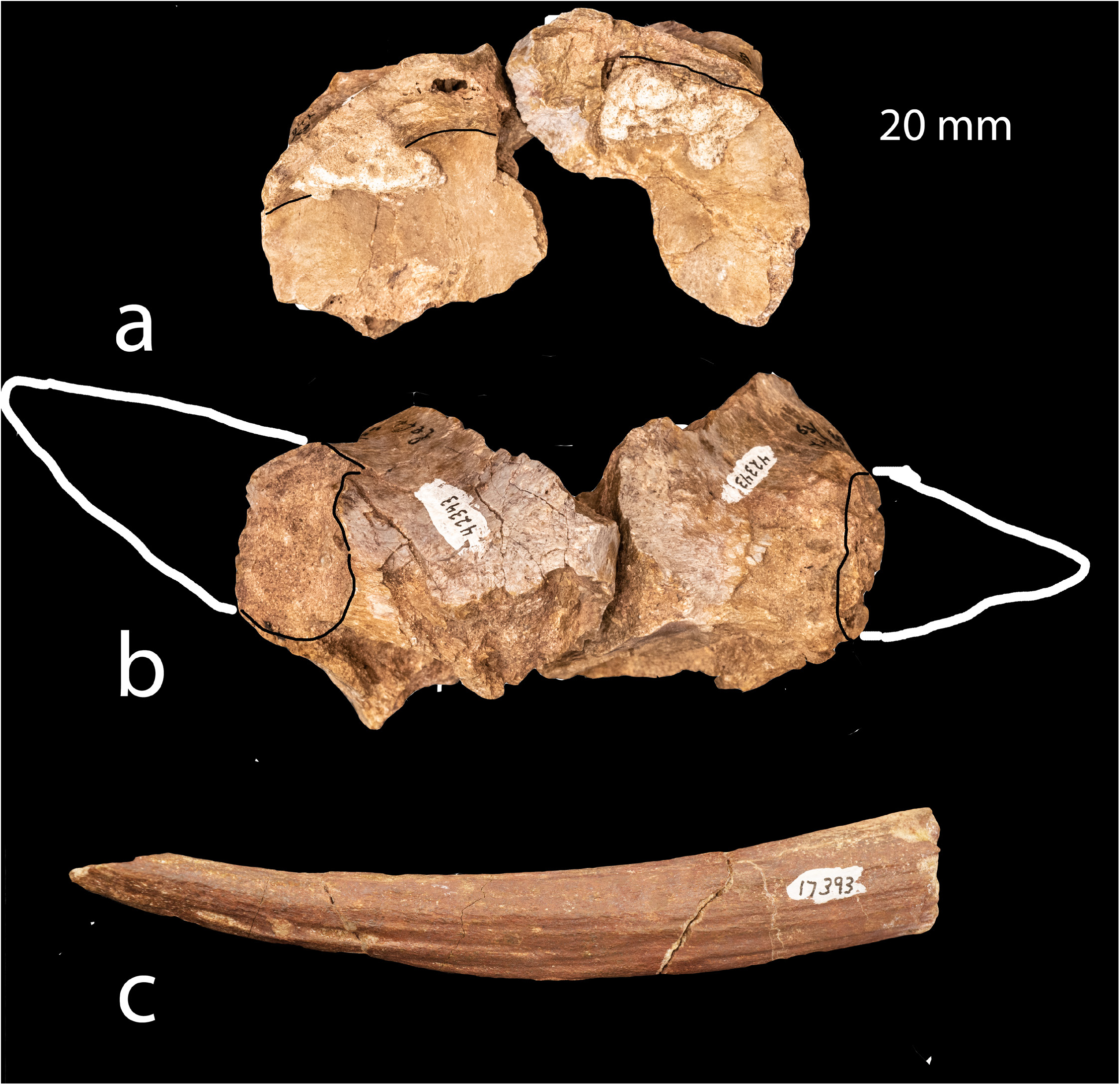
*Tauromeryx canteroi* A) occipital bone with base of horns Y 42343. White line shows possible direction of horns; B) occipital horns Y 42343 in dorsal view; C) Y 17393 frontal horns not associated specimen.

*Tauromeryx canteroi* nov. sp.

Holotype- YGSP 42343, two occipital horn fragments belonging to the same individual. Type locality - Y0502.

Age- 11.951-11.99 Ma.

Etymology- *canteroi*-Named after Enrique Cantero, preparator at the MNCN.

Diagnosis- The ossicones above the orbits are thin and long and slender curving gently back. The grooving is minimal. The horns are at the occipital edge. They are broken off, but the base is wide, and they seem to thin abruptly. They are deeply grooved. The occipital surface is remarkably flat with no details of nuchal ligament. There is distortion and the left horn is oriented more lateral than the other. The radius (Y GSP 26792) is much shorter than the metacarpal. This is the only ruminant with such a specialization (If the two specimens belong to the same individual – (J. C. Barry personal communication claims they are). The astragalus has a well-developed head with distinct ridges. Astibia et al. (1998) described a *Tauromeryx* specimen. It is similar to the current species but differs in the horn morphology (Fig. 1b).

Dimensions: Y 42343: width and length 3.1? x 3.65; 17393 horn above orbit Y 17393 width at base the base it is 2.92 x 1.60 cm and the preserved length is 13.45 cm. That is not along the curve, which would be longer.

*Nuchalia gratia* nov. gen. nov. sp.

Holotype- Y GSP 30933, partial skull, detached ossicone and two maxillary fragments.

Type locality – Y0691.

Age- 13,032-13,137 Ma.

Etymology- Named *Nuchalia* due to the deep nuchal pits on the occipital. Gratia-means thankful as for the finding of a skull in the Siwaliks is rare.

Diagnosis- The braincase is crushed lateromedially. The calvaria is flat and there are no parietal or frontal sinuses. The ossicone has smooth surface and small longitudinal ridges. The apex of the ossicone is blunt and the cross-section is oval-triangular. The ossicone is a typical shape similar to the *Okapia*. It gently curves back it is triangular in cross section with no canals. The apex is a simple round. The occipital has two very deep pits for the funicular nuchal ligament. The bullae are oval and large. The occipital condyles are merging in the median plane. The posterior basioccipital tuberosities are strongly lateral thin ridges. The anterior basioccipital tuberosities are large ovals. The glenoid fossae are large. There is a *Palaeomeryx*-fold in the molars. The buccal cingulum is very small. The lingual cingulum is also very small and there are no basal pillars. There was an occipital horn which is broken off. The edge of the occipital shows this by the presence of two strong supporting ridges. This species is similar to *Palaeomeryx* but it differs in the occipital horns and the nuchal pits. Similar ossicones were found at Kalodirr (undescribed) (Fig. 2 and 3).

**Fig. 2.**
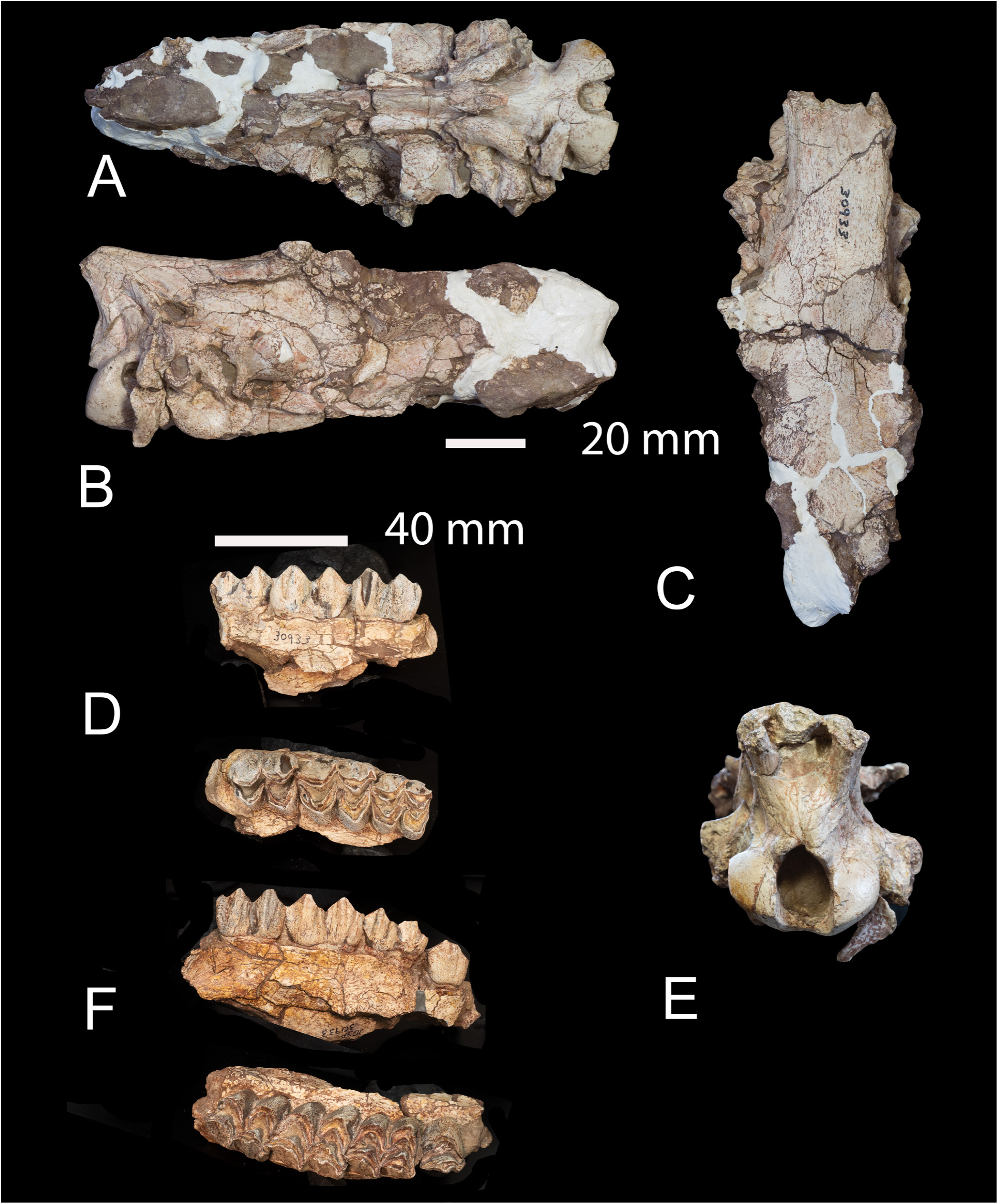
*Nuchalia gratia* holotype YGSP 30933 A) ventral view; B) right lateral view; C) dorsal view; D) occipital view; E) right maxilla; F) left maxilla.

**Fig. 3.**
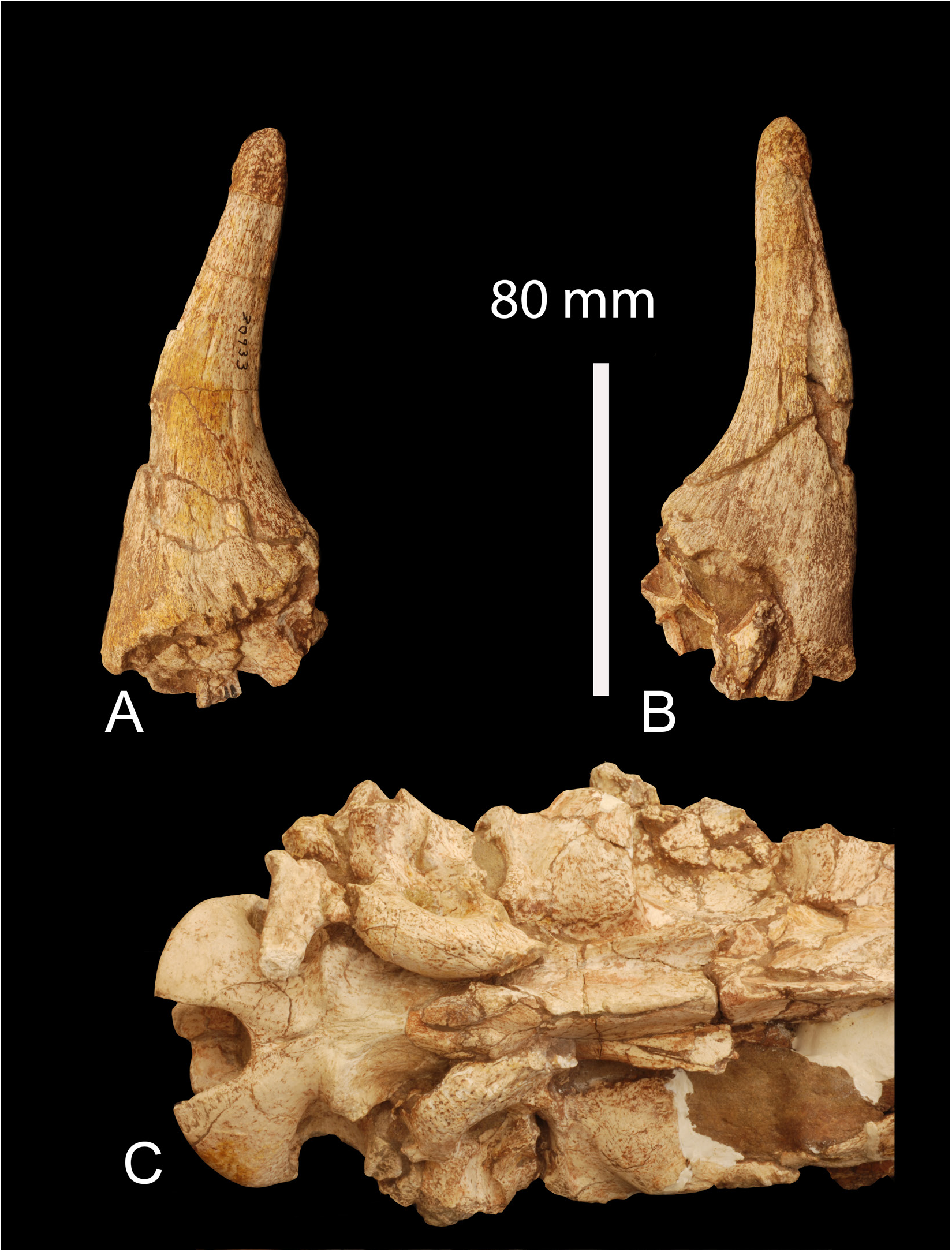
*Nuchalia gratia* holotype YGSP 30933 A) ossicone dorsal aspect; B) ossicone ventral aspect; C) basicranium. The ossicones were oriented lateral and dorsal.

Dimensions: Y 21897: ∼3.45 x ∼ 2.55 (cm); Y 23715 is from Locality Y 311.

*Fovea fossata* nov. gen. nov. sp.

Holotype- Y GSP 21897, detached ossicone.

Other specimens- Y GSP 23715, detached ossicone from locality Y0311.

Type locality- Y0504.

Localities- Y0504 (11.6-11.6 Ma.), Y0311 (10,026-10,100 Ma.)

Age- 10.0-11.6 Ma.

Etymology- *Fovea*- is pit as in the eye, *fossata* is fossa.

Diagnosis-species with small ossicones of triangular short shape. They curve inward as most other palaeomerycid ossicones. The apex is blunt and the cross-section is oval. There is a remarkable ornamentation in the form of little fossae that cover the whole surface of the ossicone. The ossicones are slightly similar to *Triceromeryx pachecoi* in having a messy surface. This species resembles *Triceromeryx* in having a disorganized horn surface (Crusafont 1952).

Dimensions: YGSP 21897 basal length 4.4 mm height 6 mm. YGSP 23715 basal width 3 mm available height 6 mm (Fig. 4).

**Fig. 4.**
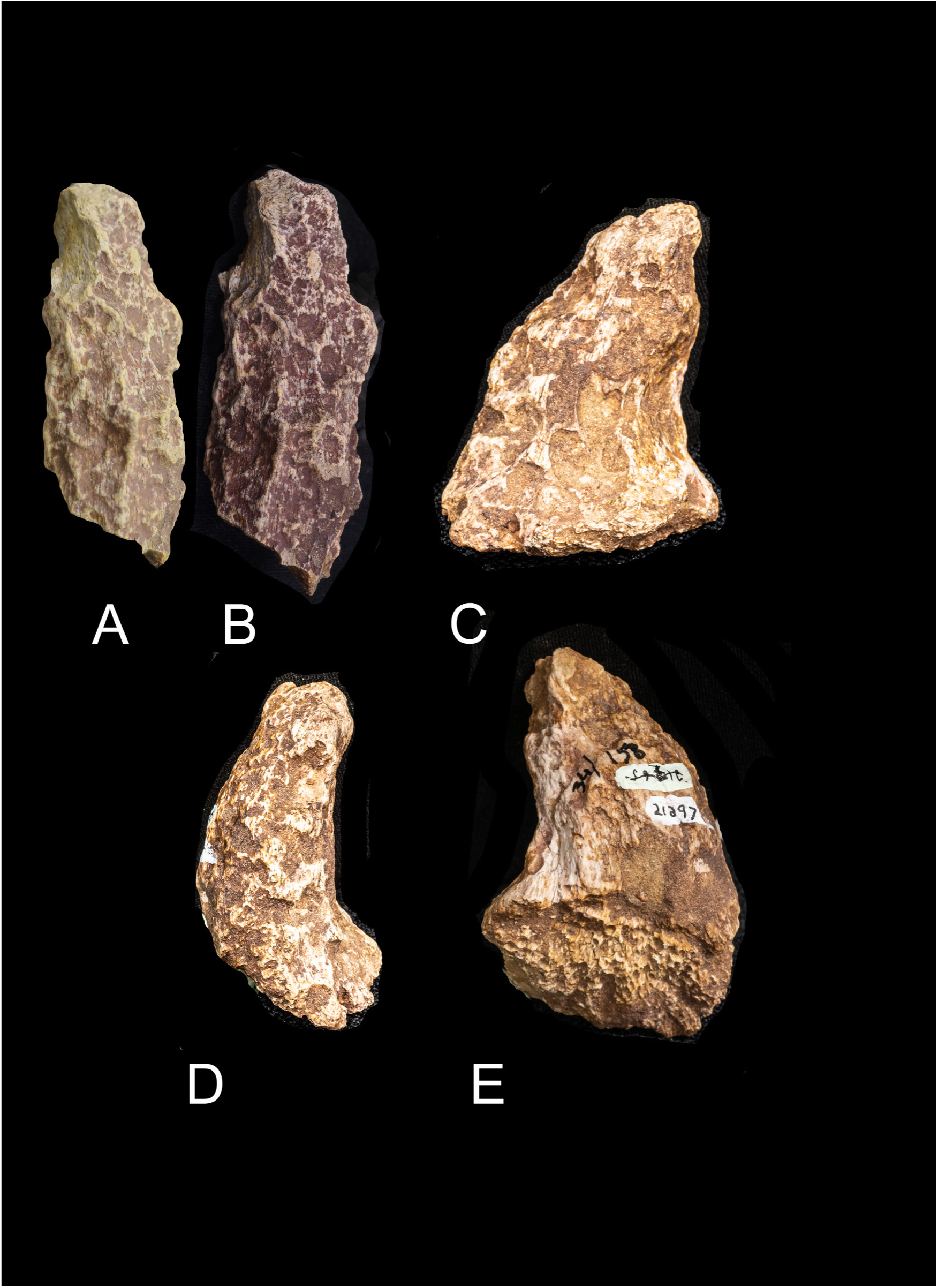
*Fovea fossata*. A and B) Y23715 and C) medial view of holotype Y21897; D) anterior view of holotype; E) lateral view of holotype.

*Goniomeryx flynni* nov. gen. nov. sp.

Holotype- Y GSP 30608, detached ossicone.

Other specimens- Y GSP 31810, detached ossicone from locality Y0750.

Type locality- Y0758.

Localities- Y0758 (13,940-14,161 Ma.), Y0750 (12,740-12,758 Ma.)

Age- 12.7-14.1 Ma.

Etymology- Gonio-is angle as the ossicone is angled in the cross sections, -meryx ruminant - flynni - the trivial name after Larry Flynn who has worked in the Siwaliks.

Diagnosis- Ossicones with thin longitudinal keels in the surface and a thin oval cross section. Remarkable lateral facet at the ossicone base with a smooth surface. The holotype has a surface under the ossicone showing that it was detached in early life. The second specimen is fused to the frontal. The specimens have a strongly angular outline with two keels a mid-anterior keel the abruptly turns laterally at the base and a posterior keel which terminates 20 mm above the base. The pedicle is absent but in YGSP 31810; a little excavation with a smooth surface is visible posteriorly (like a pedicle). The ossicone is above the orbit. The orbit is typical with no specializations. There is no internal canal. The shaft of the ossicone thins abruptly (Figure 5).

**Fig. 5.**
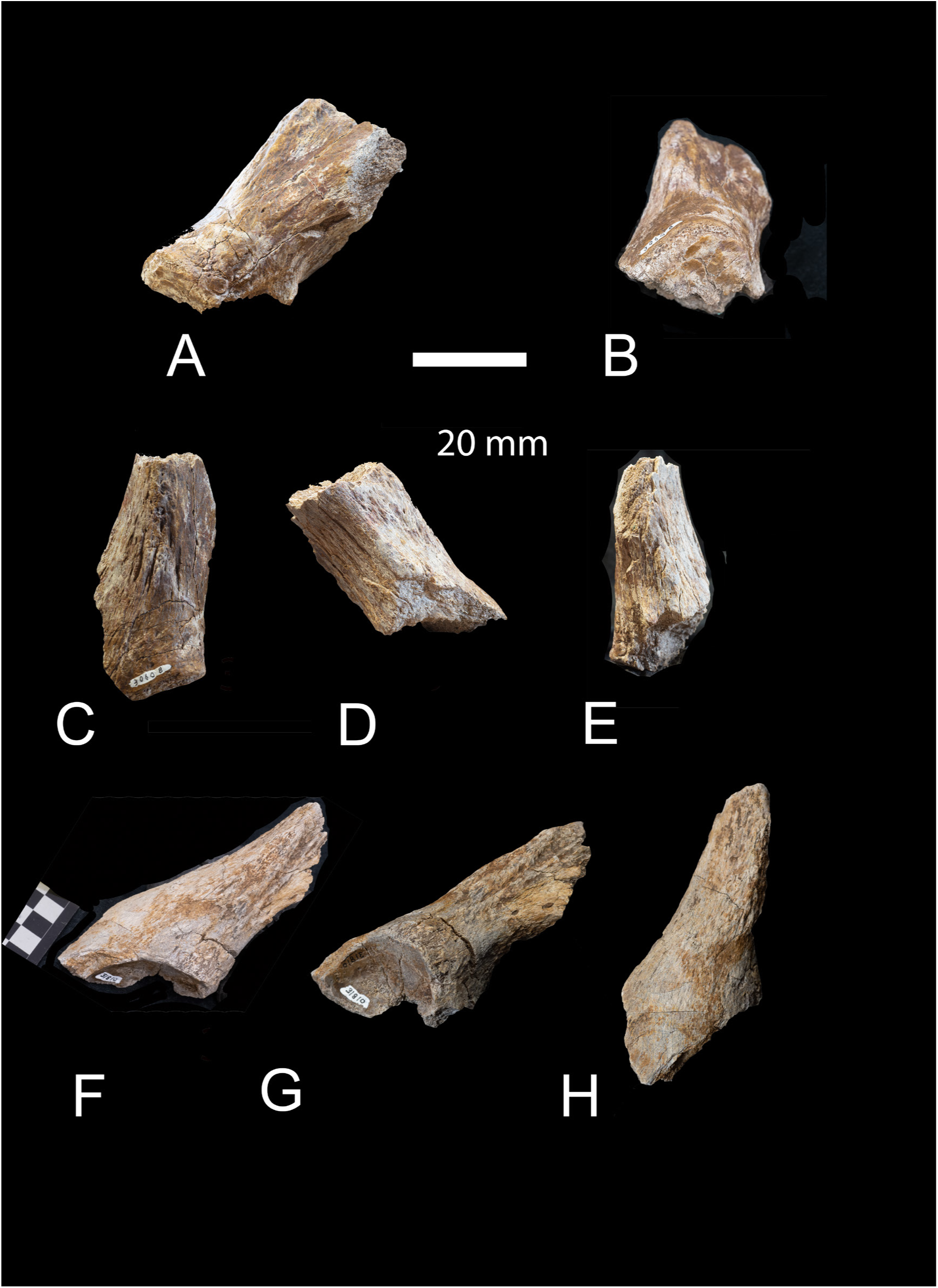
*Goniomeryx flynni* holotype Y 30608 A) lateral view; B) ventral view showing the detached ossiocne; C) anterior view; D) lateral view; E) posterior view; F) 31810 loc 758 14 Ma dorso-lateral view; G) lateral view; H) anterior view.

Dimensions: YGSP 30608 is 35.7 × 34.4 mm; YGSP 31810 is 39.3 × 34.0 mm.

*Lateralia morgani* nov. gen. nov. sp.

Holotype- Y GSP 30593, ossicone and skull fragment.

Type locality- Y0500.

Age- 12- 12.1 Ma.

Etymology- *Lateralia*- refers to the lateral position of the ossicones, -*morgani* named after Michèle Morgan who has worked in the Siwaliks.

Diagnosis- The smallest palaeomerycid. The ossicone is situated laterally on the frontal behind the orbit. There is grooving on the ventral side the shaft which is straight. At the base there is a restricted sinus. There are no keels and no canal internally. The lateral margin of the ossicone extends further inferiorly than the medial margin. It has a maximum length of 17.8 mm in antero-post direction (Figure 6).

**Fig. 6.**
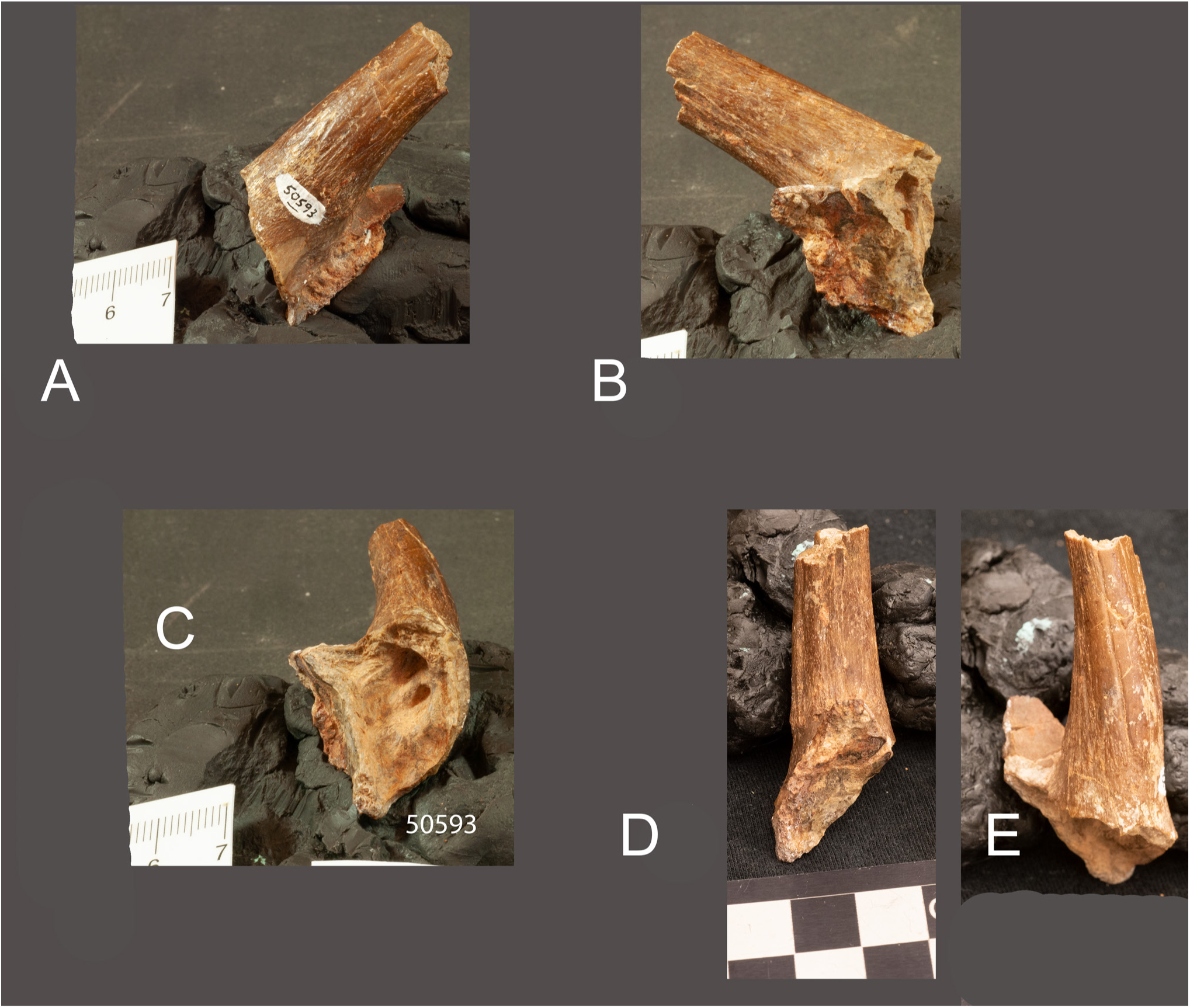
*Lateralia morgani* hototype Y 50593. A) ventral view; B) medial; C) the axis of the ossicone view; D) posterior view; E) dorsal view.

Dimensions: Maximum length of 17.8 mm in antero-post direction. The width perp to the maximum length (not a cross section to max length but measuring maximum extent of ossicone in this orientation) is 15.9 mm. The length is 13.8 mm. It is possible that this taxon maybe related to *Orygotherium* as it is also very small. But *Orygotherium* is known only from dentitions (Vislobokova 2004). Alan Gentry has identified this specimen as a bovid (Personal communication), (Fig. 6).

### Family Climacoceridae Hamilton 1978

#### Climacocerinae new rank

Diagnosis- Taxa with ossicones that branch on the anterior part of the shaft superficially resembling those of deer. Most taxa form horn shafts rounded cross sections and do not taper going up from the base. One taxon had no branches (*Vittoria*). Another taxon had complex flat horns with branches at the edge (*Prolibytherium*). The flat ossicones were smooth with vein impressions. The rounded ossicones had regular fine homogeneous grooving. Grooving may be actual hair embedded in the bone and fossilized (as YGSP 31790). The fusion of the ossicone to the frontal leaves no markings or sutures. The apices are pointy.

Ossicone structure: Two specimens were available. In one (of YGSP 31790) the cortex is composed of fossilized coarse hair – and an inner structure composed of disorganized trabeculae). In YGSP 47197 (gen. indet. a cortex composed of isolated hair in a thick matrix and inner trabeculae which are connected by lateral extensions). In UCMP 40461 it is detached from the skull. The base of this ossicone (known from *Pachya*) differs from the giraffe and the okapi. It has flattened small interlocking plates with no openings between them. In the giraffe there are clusters of bony ossifications and large openings for a matrix.

#### Climacocerinae

*Vittoria soriae* nov. gen. nov. sp.

Holotype- NHM M 26690 (identified as a palaeomerycid by Hamilton (1973) and a female *Prolibytherium* by Sánchez et al., 2010). Curated at the NHM (London).

Siwalik specimens- Y GSP 31790, ossicone shaft fragments (loc Y0747), Y GSP 102058, horn fragment (loc. Unknown).

Type locality-Gebel Zelten (Libya). Localities- Y0747. 17.8-17.9 Ma.

Age-Lower Miocene.

Etymology- Vittoria- refers to the V shape of the cranial appendages. V signifies victory - as used by Churchill -*soriae*- Named after. Dolores Soria, the first female vertebrate paleontologist to study giraffids.

Diagnosis- A species with straight unbranching ossicones. There are no keels on the ossicones which curve gently inward. The central canal is large and extremely oval. There is a thin cortex. The Siwalik specimens have a longitudinal groove on one side. The skull roof is very thick. The surface of the ossicones has fine grooving which is due to the excellent preservation. We hypothesize that this is fossilized hair embedded into the surface. The slope angle of the appendage relative to the cranial roof is 40°. Described as a female *Prolibytherium* by Sánchez et al. (2010). It differs from *Prolibytherium* in numerous ossicone characters (Figures 7 and 8). *Climacoceras* from the Miocene of Kenya has forked ossicones with small tines (MacInnes 1936; Hamilton 1978). The un-forked ones as (NHM M 26690) were published as a female *Prolibytherium* by Sánchez et al. (2010).

**Fig.7.**
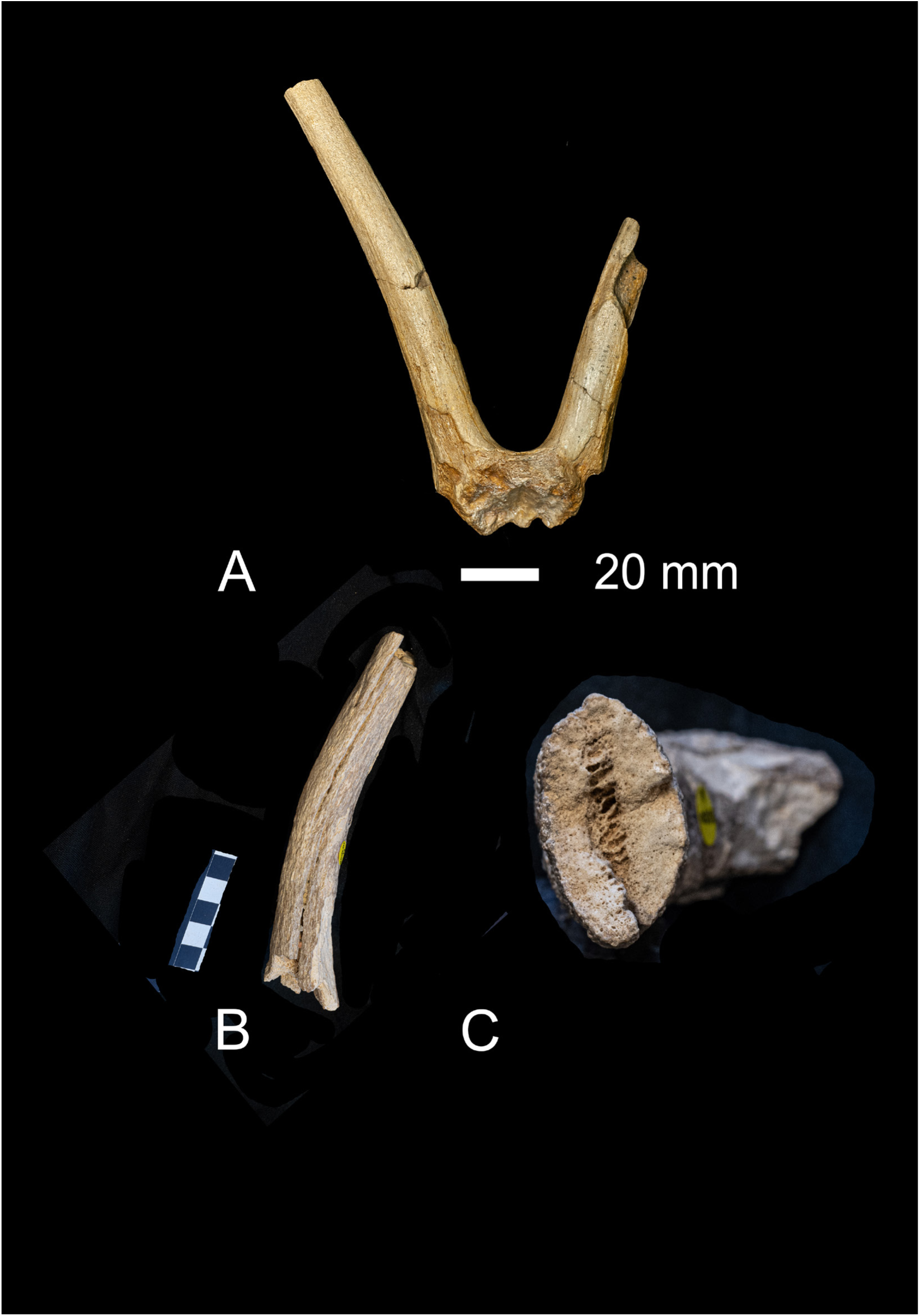
A) *Vittoria soriae* NHM M 26690 (holotype); B) NHM M 102058 both from Gebel Zelten. C) close up of b showing a cross section. Note there is an ossicone sinus.

**Fig. 8.**
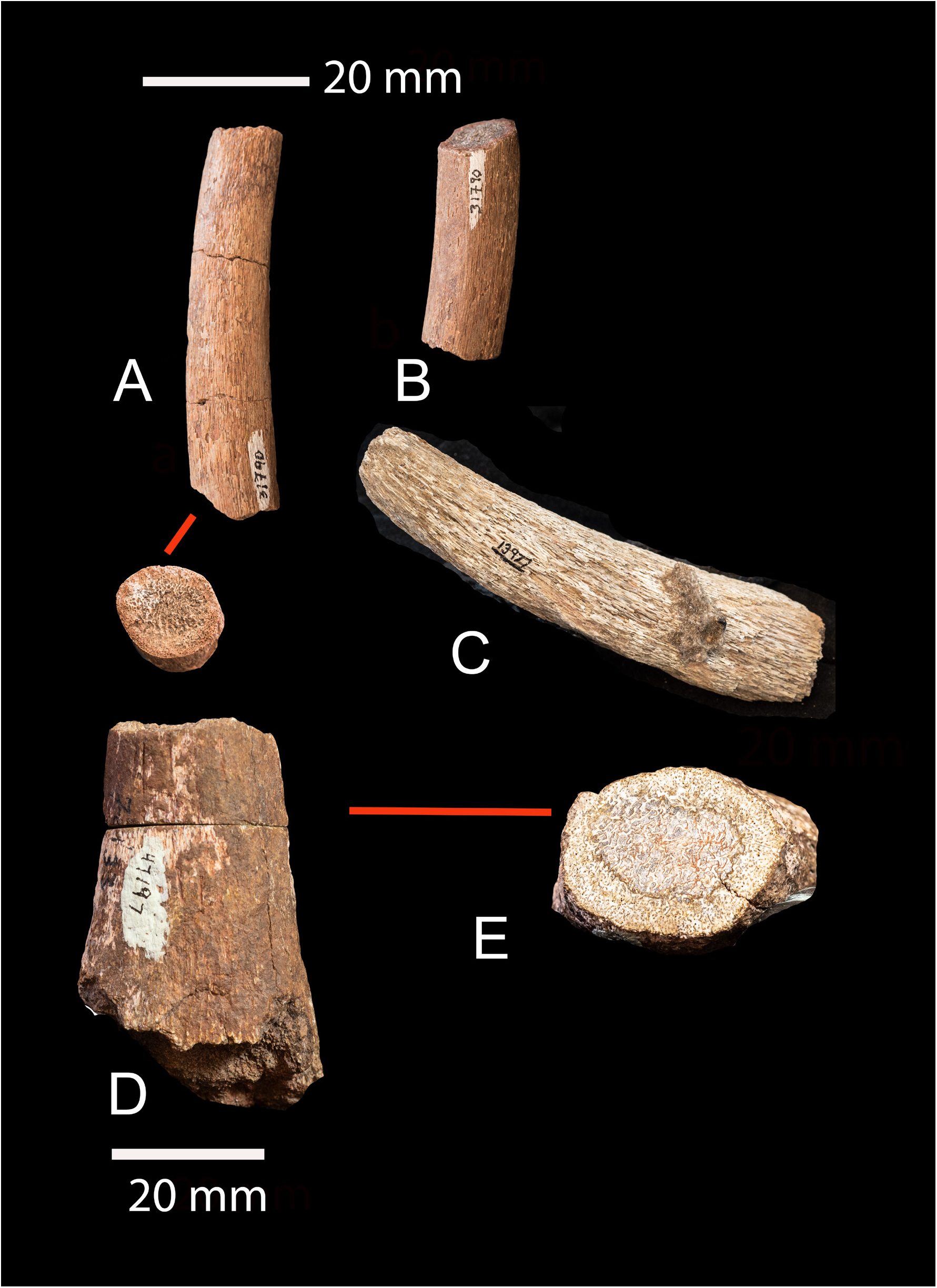
*Vittoria or Climacoceras*. a and b) Y 31790 fragments. No sinus in the ossicone. C) Y 31927 ossicone fragment. The fine striations are hypothesized to be ossified hair; D and E) Y 47197 a large specimen where the double cortex is clearly visible in the cross section. The outer is hair embedded in a white matrix. The inner is elongated laterally haversian structures. Both from the Chinji.

Dimensions: Holotype NHM M 26690 width at base 18 mm. We add specimens that might also be *Vittoria* in the sample: *Climacoceras* YGSP 31927 and YGSP 31790 both from the Chinji (Fig. 7 and 8).

#### Prolibytheriinae

Genus *Prolibytherium* Arambourg, 1961

Type species *Prolibytherium* magnieri Arambourg, 1961

**Locality**—Gebel Zelten, Libya (middle Miocene).

*Prolibytherium fusus* Danowitz et al. 2016.

**Referred Specimen -** *Prolibytherium fusus* PMNH Z 162 a braincase with broken cranial appendages and well-preserved occipital and basioccipital.

**Description—**The Zinda Pir specimen PMNH Z 162 is a braincase. The cranial appendages have been broken off. A description is in Barry et al. (2005).

**Diagnosis**— *Prolibytherium fusus* differs from *Prolibytherium magnieri* in the following characters: the anterior basioccipital tuberosities are less distinct, and their surface contains longitudinal ridges instead of small bumps seen on *P. magnieri*. The elongated fossa between the posterior and anterior basioccipital tuberosities is one unified surface in *P. magnieri* and is separated into two plates by a midline longitudinal groove. The basioccipital bone is boxy-shaped and elongated rectangular-shaped in *P. magnieri*. In *P. fusus*, the posterior basioccipital tuberosities are more approximated with the articular surface of the occipital condyles, and are thicker and shorter. The posterior basioccipital tuberosities are also continuous across the midline in *P. fusus*, separated only by a narrow deep groove, and the caudal surface contains several bony growths concentrated medially. In *P. magnieri*, the posterior basioccipital tuberosities are separated at midline, and the surface contains a distinct elevated transverse ridge, and no bony growths. The U-shaped ventral margin on the occipital condyles is shallower in *P. fusus* and the condyles are oriented laterally, whereas the condyles are oriented dorso-laterally and have a deeper U-shaped ventral margin in *P. magnieri*. The notch between the para-occipital process and the lateral occipital condyles is thicker and lower in *P. fusus*.

**Comments**—Two studies have focused on *Prolibytherium magnieri.* Hamilton (1973) has described in detail the material, and Sanchez et al. (2010) propose that sexual dimorphism exists with females possessing cranial appendages that retain the same pattern but are thin and cylindrical and without the webbed aliform connection. Barry et al. (2005) briefly compares *Prolibytherium* to *Progiraffa* and they explore the large ruminant material from Zinda Pir as only *Progiraffa*.

The discovery of *Prolibytherium* from the lower Miocene of Pakistan, extends the genus to Asia. *Prolibytherium* is currently known from Libya; Gebel Zelten. The new single specimen *Prolibytherium fusus* (Danowitz et al. 2016) is a partial braincase and the ossicones are broken off. This species differs from *P. magnieri* in several basioccipital and atlanto-occipital morphologies. Namely, the posterior basioccipital tuberosities are continuous at midline and lack the elevated transverse ridge seen in *P. magnieri*, and the notch formed between the lateral occipital condyles and para-occipital process is lower. Both species of *Prolibytherium* have a characteristic ventrally fused occipital condyle at midline, with a notably fuller circumferential surface. *P. magnieri* also has thickened dorsal and ventral arches of the atlas. We believe these, plus several other atlanto-occipital morphologies strengthen the cervical support of the head. This is especially important for *Prolibytherium*, as the taxon possesses massive ossicones. The fusion of the occipital condyles is a unique character.

**Locality and Geological Age**—Zinda Pir of Pakistan (early Miocene). It was found in Locality Z 124 which is part of the Vihowa Formation. It is located at the top of a normal polarity zone that has been correlated with the 6n polarity interval. It would be slightly older than 18.7 Ma (Lindsay et al. 2005).

Barry et al. (2005) identified the braincase PMNH Z 162 from Zida Pir as belonging to *Progiraffa exigua*. Numerous postcranial specimens, a giraffid maxilla and an isolated horn were also identified as *Progiraffa* exigua. That horn is a *Hypsodontus*.

The newer finds from Zinda Pir have been connected to the genus *Progiraffa* (Barry et al. 2005). The basis of this connection was size, the geographic proximity and the very old age of these ruminants. However, is common in the Miocene to find three or more giraffid species per locality. At this time, we do not exclude the possibility of the presence of *Progiraffa* in the Zinda Pir collection. We simply exclude the braincase from the remainder of the *Progiraffa* sample, because it better matches that of *Prolibytherium*. The Z 162 braincase is clearly assignable to *Prolibytherium*. The new species *fusus* extends the range of *Prolibytherium* into Asia at 18.7 Ma.

*Orangemeryx badgleyi* nov. sp.

Holotype-Y GSP 23698.

Type locality- Y0666

Age- 13.7-13.8 Ma.

Etymology- *badgleyi* - Named after Catherine Badgley who has worked in the Siwaliks.

Diagnosis- A base of a right ossicone and part of frontal. The surface is smooth There is a medial keel. Laterally the edge makes a flat surface which can be also considered as a broad keel. No sutures with the frontal are visible. The frontal sinus under the horn is well-developed. The sinus has several large bony cells in it and is filled with mud. The current species is very similar to *Orangemeryx* (Morales et al. 1999) (Fig. 9).

**Fig. 9.**
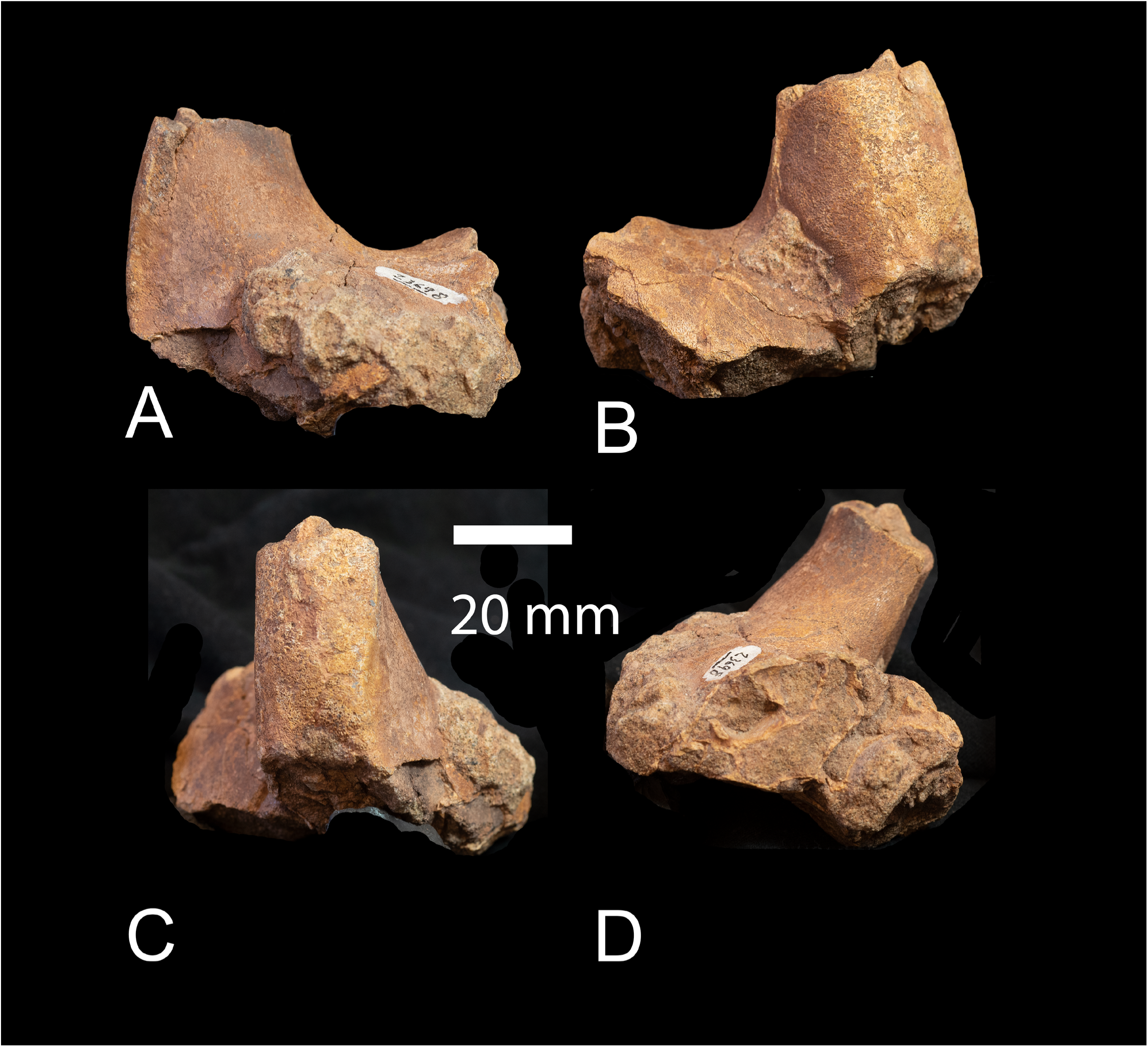
*Orangemeryx badgley* 23698 loc 666 13.7 Ma. A) posterior aspect; B) anterior aspect; C) lateral; medial view – note the large frontal sinus.

Dimensions: YGSP 23698: 4.15 x 3.65 (cm).

*Pachya moruoroti* nov. gen. nov. sp.

Pachya means fat-puffy in Greek as in pachyderm. The trivial name is for the locality. A single ossicone from Moruorot Hill Kenya (UCMP 40461). It is from the Lothidok Formation of West Turkana. It is detached from the skull (unfused). The base of this ossicone differs from the giraffe and the okapi. It has flattened small interlocking plates with no openings between them. It is the only evidence of an ossicone base in all Climacoceridae. The structural difference from *Giraffa* and *Okapia* may explain that is Climacoceridae there are no lineations of fusion at the junction of the ossicone with the skull. One explanation is a strong reworking of the bone after fusion to the frontal. This is best visualized in *Prolibytherium* from Gebel Zelten at the NHM where there at least 5 skulls with ossicones fused. The ossicone is thick in *Pachya* and it has two branches which are also thick. The surface is smooth. The shaft is relatively straight (Grossman and Solounias 2013).

Comment: We include this species only because it reveals the unfused ossicone. This is critical in determining that Climacoceridae had ossicones. Form all Climacoceridae the only specimen known it us of Climacoceridae which shows the ossicone detached is that of *Pachya moruoroti.* This species has not been found with certainty in the Siwaliks. Fig 10 d and e may be Pachya.

**Fig. 10.**
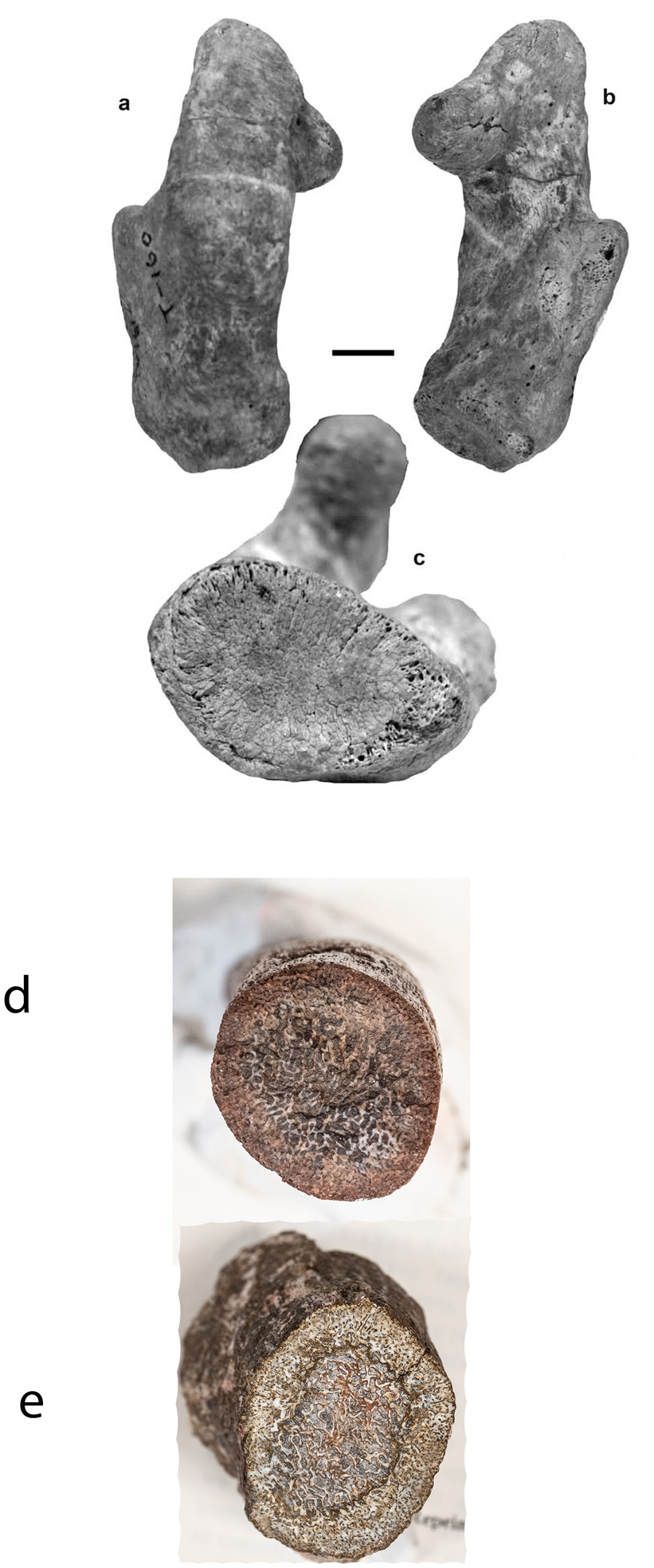
*Pachya moruoroti* nov. gen. nov. sp. from Moruorot Hill Kenya (UCMP 40461). It is from the Lothidok Formation of West Turkana. Figure from Grossman and Solounias (1013). a) posterior aspect; b) anterior aspect; c) attachment side. d) cross section of YGSP 31790 *Vittoria soriae* (the ossicone section reveals a cortex composed of fossilized thick hair – and an inner structure composed of disorganized trabeculae) and e) YGSP47197 (gen. indet. A cortex composed of isolated hair in a matrix and inner trabeculae which are connected by lateral extensions).

## Family

### Giraffidae Gray 1821

#### Progiraffinae new rank

*Progiraffa exigua* Pilgrim, 1908

Holotype- Mandible with m2 and m3 (Pilgrim, 1908, Plate 1: Fig. 1). Geol. Surv. Ind.- XXXVII. p. 165.

Type locality- Upper Nari beds in the Bugti Hills.

Age- Lower Siwaliks, Early Miocene.

Etymology- *Progiraffa* means early *Giraffa*, and *exigua* refers to the small size.

Diagnosis- Small giraffid. Ossicones supraorbital and projecting laterally slightly. Brachydont dentition with rugose enamel, and bilobed canines with pointy larger medial lobe. Upper molars with distinct bifurcation of postmetaconule crista, postprotocrista and premetaconule crista. Anterolingual cingulum prominent. Lower molars with large metastylid well separated from metaconid, m3 with robust entoconulid, *Palaeomeryx* fold absent (modified after Pilgrim, 1911, and Barry et al., 2005). Occipital simple and calvaria without sinuses. Astragalus very robust. Metapodials long (Figures 10, 11, 12).

Material: MT Y 24305 MT and Y 42342 MT are assigned to *Progiraffa*. MC 26709 and MT 4549 are also part of this taxon. Two astragali 41182 and DGK 12 and may belong to *Progiraffa*. Also, maxilla PSAP 312 and mandible PASP 41662 are *Progiraffa*. *Georgiomeryx georgalasi* skull (THB 30 MN5) form Thymiana Chios Island (de Bonis et al. 1997) (Figures 11, 12, 13, 14).

**Fig. 11.**
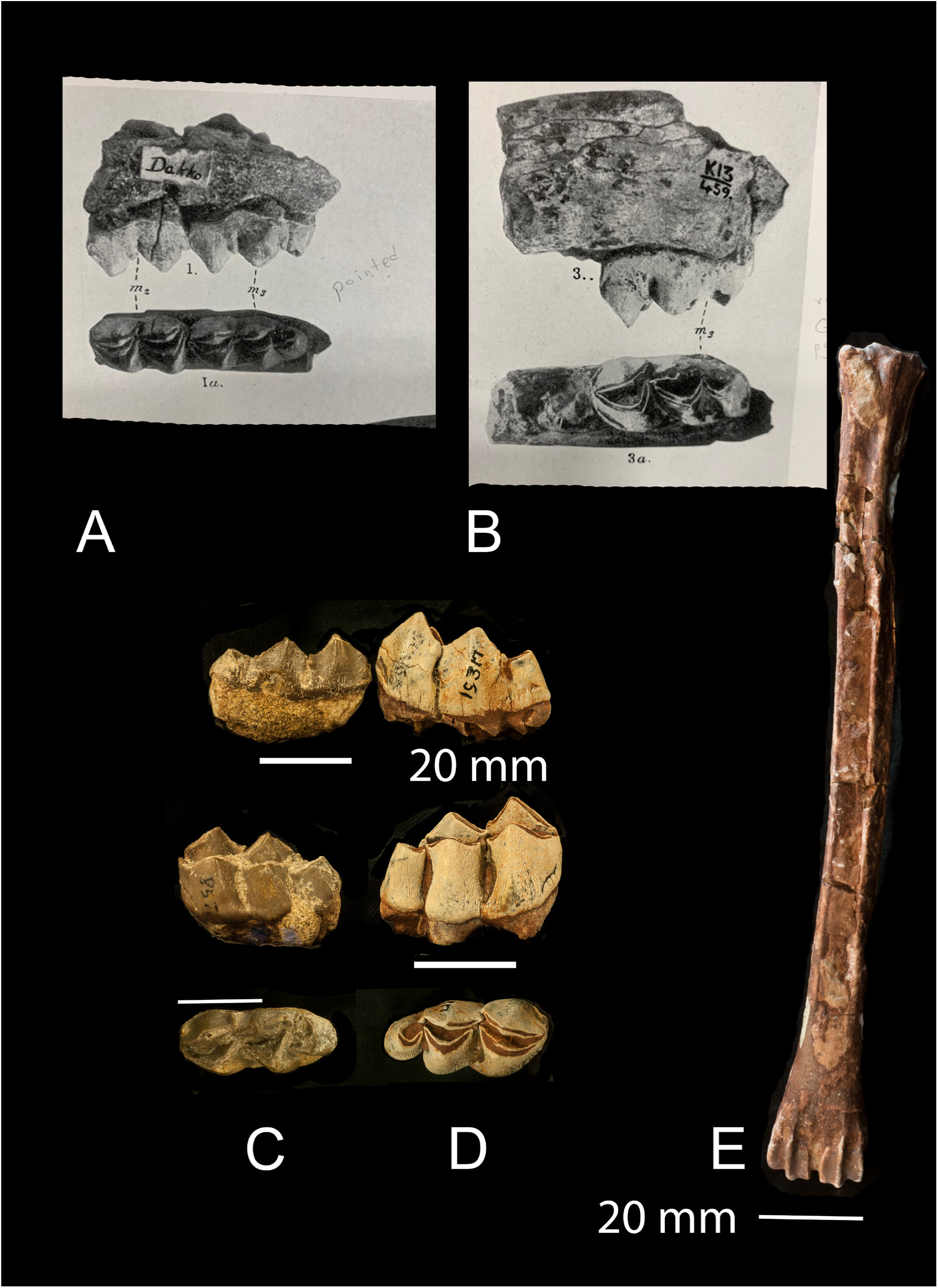
*Progiraffa exigua* A) lower m2 and m3 holotype from Pilgrim 1911 plate one; B) Progiraffa sivalensis from Pilgrim 1911 (probably also exigua) plate one; C) a similar m3 from AMNH 857 from the Chinji; D) m3 of Giraffokeryx AMNH 19317 for a hypsodonty comparison; E) Y 42342 MT.

**Fig. 12.**
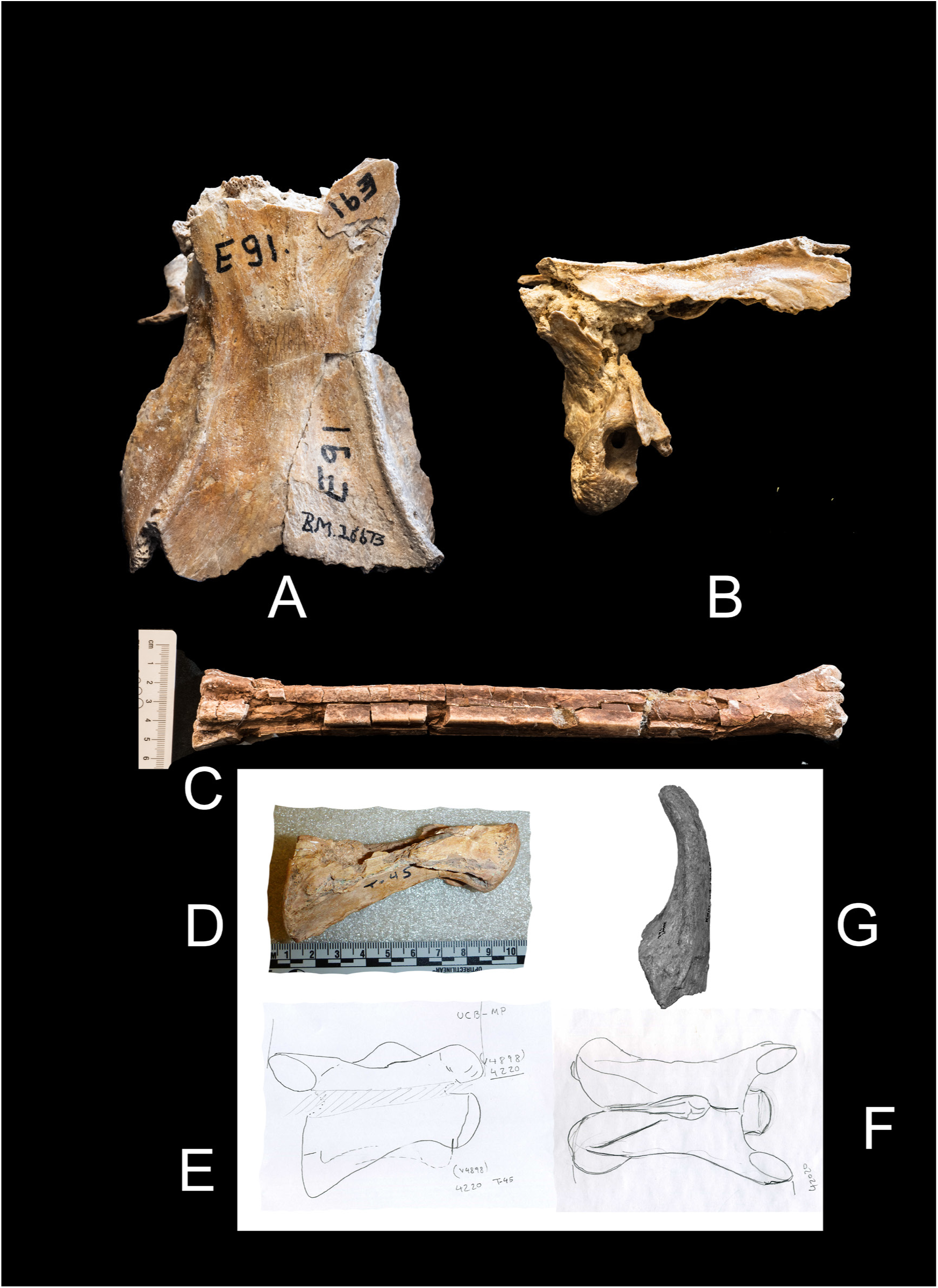
*Progiraffa exigua* A and B) dorsal and lateral view of occipital/parietal NHM M E91 26673 from Gebel Zelten Libya; C) MT YGSP 24305 from the Chinji; D, E and F) C3 vertebra restored UCM 4220 from Moruorot Hill Kenya; G) NMK K 722- 17089 ossicone from Kalodirr Kenya.

**Fig. 13.**
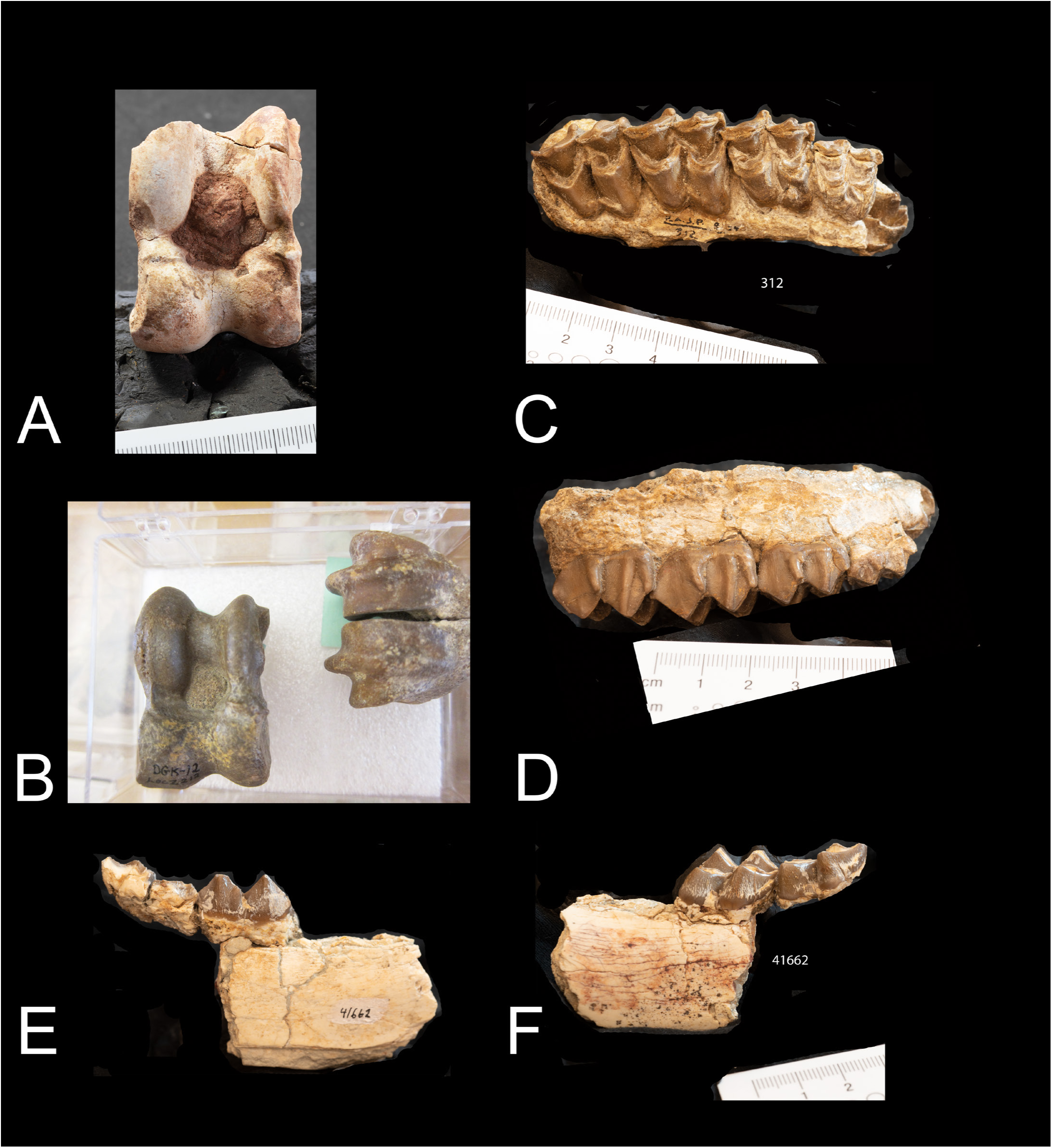
*Progiraffa exigua.* Two astragali A and B) 41182 and DGK 12 and may belong to *Progiraffa*. C and D) maxilla PSAP 312; E and F) mandible PASP 41662 of *Progiraffa*.

**Fig. 14.**
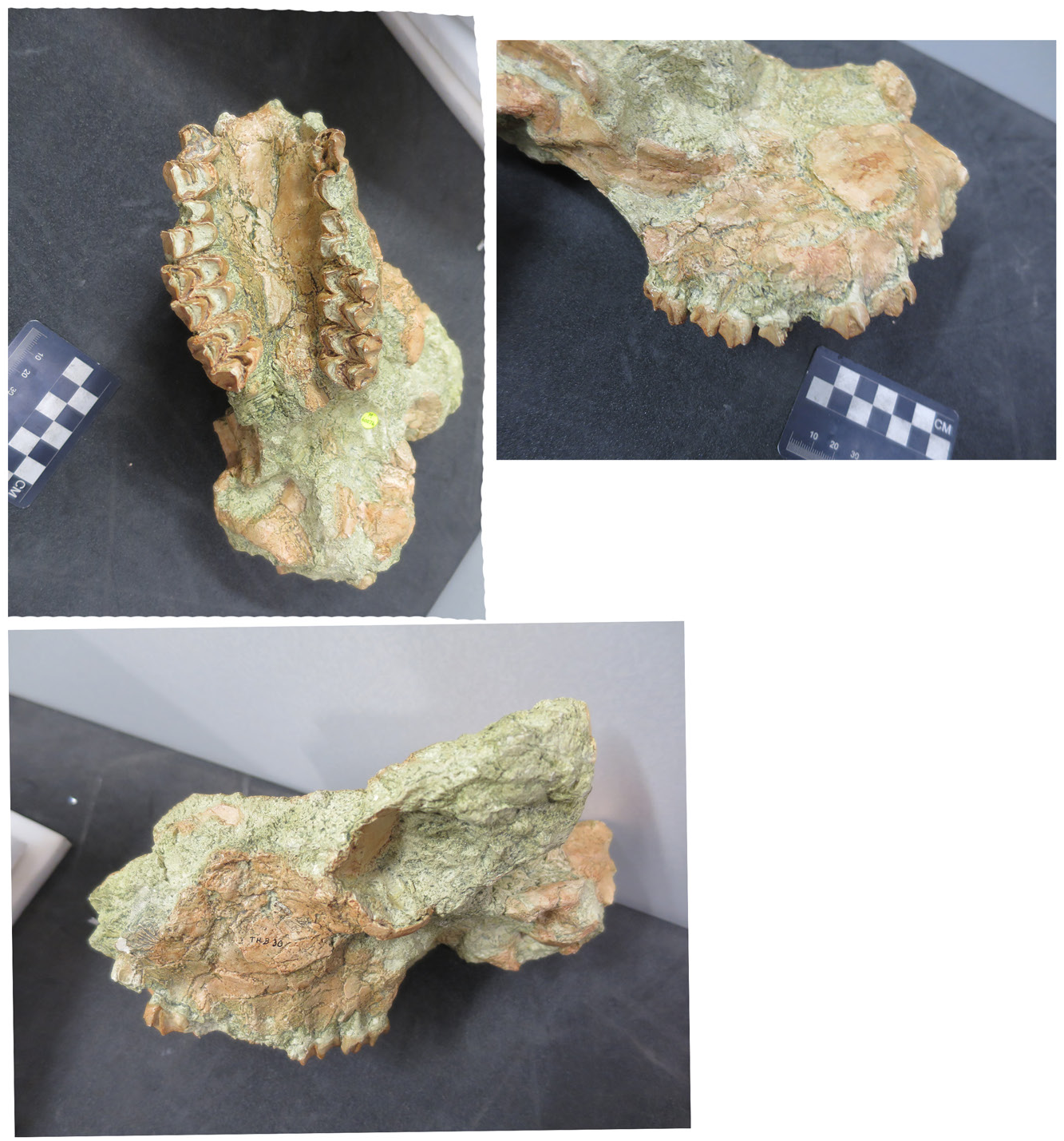
*Progiraffa exigua* skull form Chios – originally named *Georgiomeryx georgalasi*.

Giraffokerycinae Solounias 2007

*Giraffokeryx punjabiensis* Pilgrim 1910

Holotype- a selected neotype AM 19475, a complete skull.

Paratypes-AMNH 19611, teeth; AMNH 19472, left maxilla with P3-M3; AMNH 19587, mandible right ramus with p2-m3. YGSP 19842 maxilla, 42433 astragalus, 29119 complete anterior pair of ossicones, 24306 complete metatarsal, 47198 ossicone juvenile, 1194, 51470, 42426, 27761, ossicones, 5361 m3, 8101 m1-m3.

A sample of smaller individuals: MCs YGSP 26790, 4505; MC 24342; 20416 MC.

Type locality- 1,000 feet below the Bhandar bone bed, and one mile south of Nathot in the Punjab. Middle Siwaliks.

Age- Lower-Middle Siwaliks. Middle Miocene.

Etymology- *Giraffokeryx-* giraffe horn - *punjabiensis*- from the Punjab.

Diagnosis- Four ossicones. Anterior pair more anterior than the orbit and left and right side fused at the base. Posterior ossicones are more posterior than the orbit grooved with simple apex. At the base of the posterior ossicones there is a cluster of disorganized bony mass. The ossicones are deeply grooved. The shafts of the four ossicones are very similar. No sinuses in the calvaria. Upper premolars shortened. Palatine indentation posterior to M3. Masseteric fossa small. Less elongated upper premolars than *G. primaveus*. Metapodials long and slender (Figures 15, 16, 17).

The original sample is mandibular fragments in the IM in Calcutta and a few are figured in Pilgrim 1911. He also did not select a type from the IM. A p4 stands out and is characteristic of *Giraffokeryx* (Pilgrim 1911 – plate II Fig. 1) (our Fig. 15D). Later Colbert and Brown described a nearly perfect skull AMNH 19311. Colbert (1935) selected an upper M3 as a type from the Pilgrim sample. This is not diagnostic. We take the skull AMNH 19311 as a lectotype/neotype.

Most researchers consider that skull as *Giraffokeryx*. Localities are the Chinji Formation for all material.

Dimensions: Metatarsal YGSP 24306 (456X32.6 mm), metatarsal YGSP 47187 (430X30), tibia YGSP 52472 (425X35 mm), radius YGSP 46575B (450X40 mm). Also see Colbert and Brown (1933) and Colbert (1935) for measurements (Fig. 15, 16, 17).

**Fig. 15.**
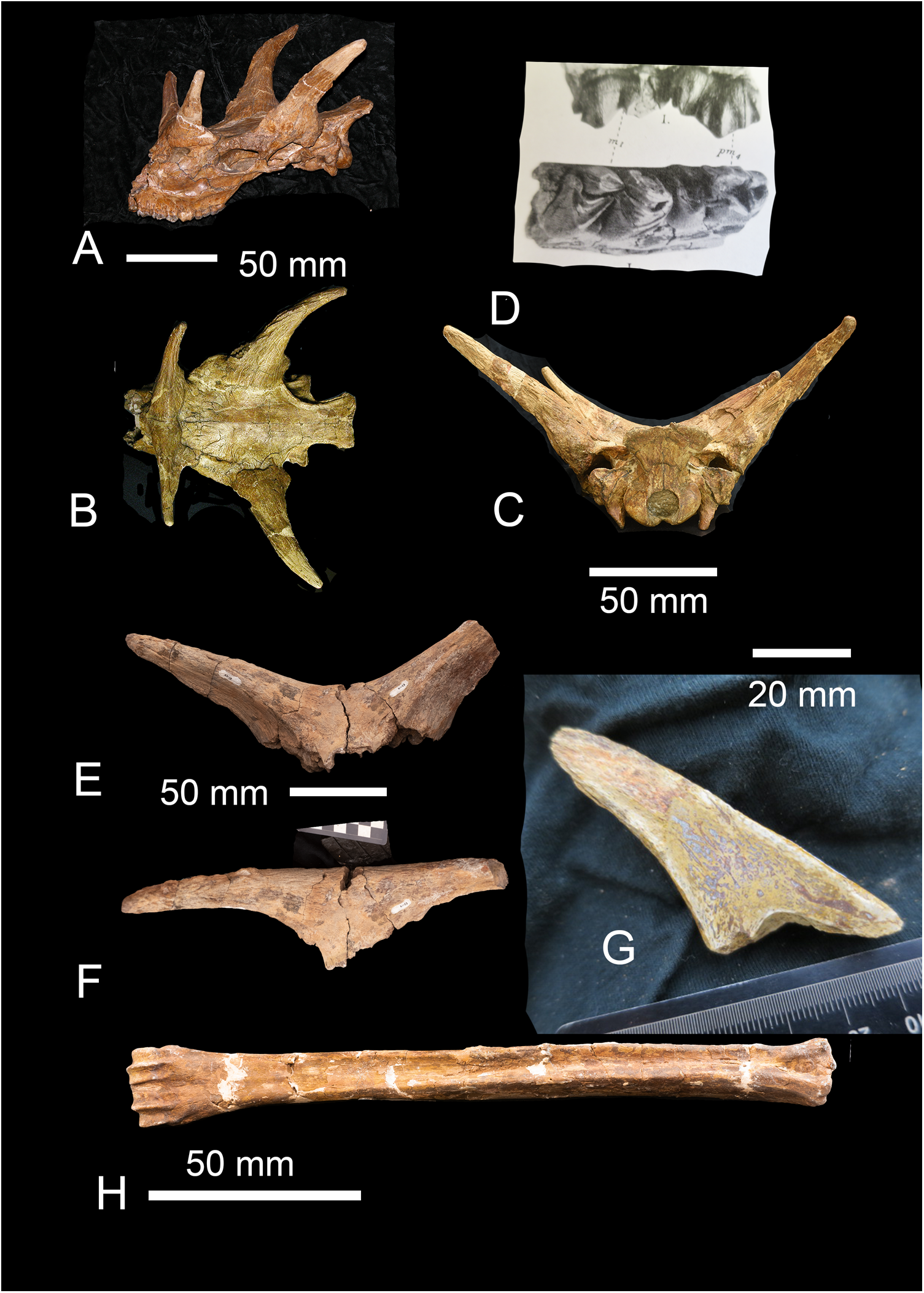
*Giraffokeryx punjabiensis* A, B, C) AMNH 19475 skull of the lectotype of *Giraffokeryx* punjabiensis; D) a p4 figured by Pilgrim 1911; E and F) anterior ossicones Y GSP 23119; G) Y GSP 47198 longitudinal section of a small ossicone assigned to this species; F) metatarsal Y GSP 24306.

**Fig. 16.**
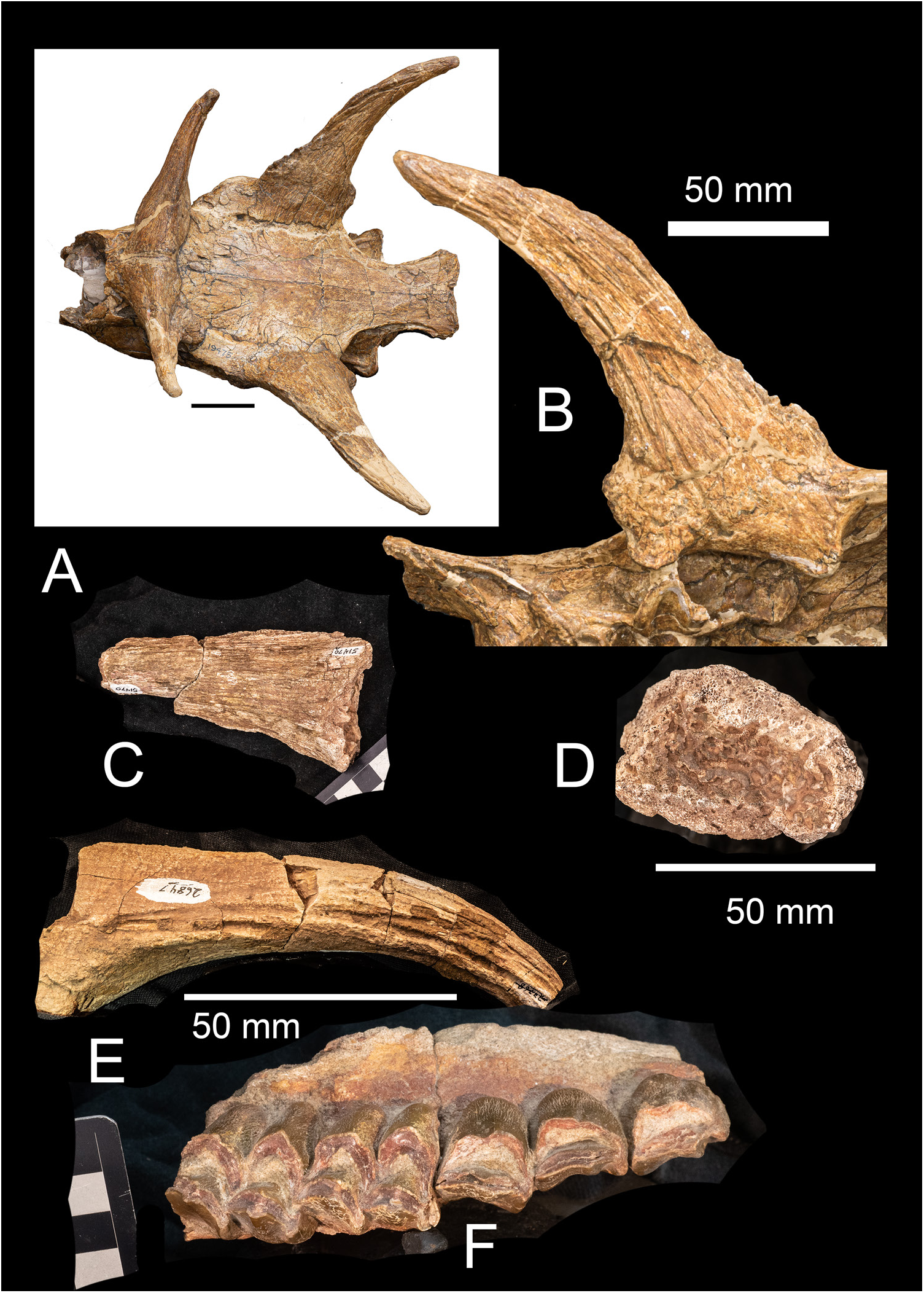
*Giraffokeryx punjabiensis*. A and B) the lectotype AMNH 19475. Note the bony mass at the base of the ossicone; C and D) Y GSP 15470 the base morphology connecting to the frontal is different from that of Giraffa; D) ossicone 26847; E) Y GSP 19842 upper dentition.

**Fig. 17.**
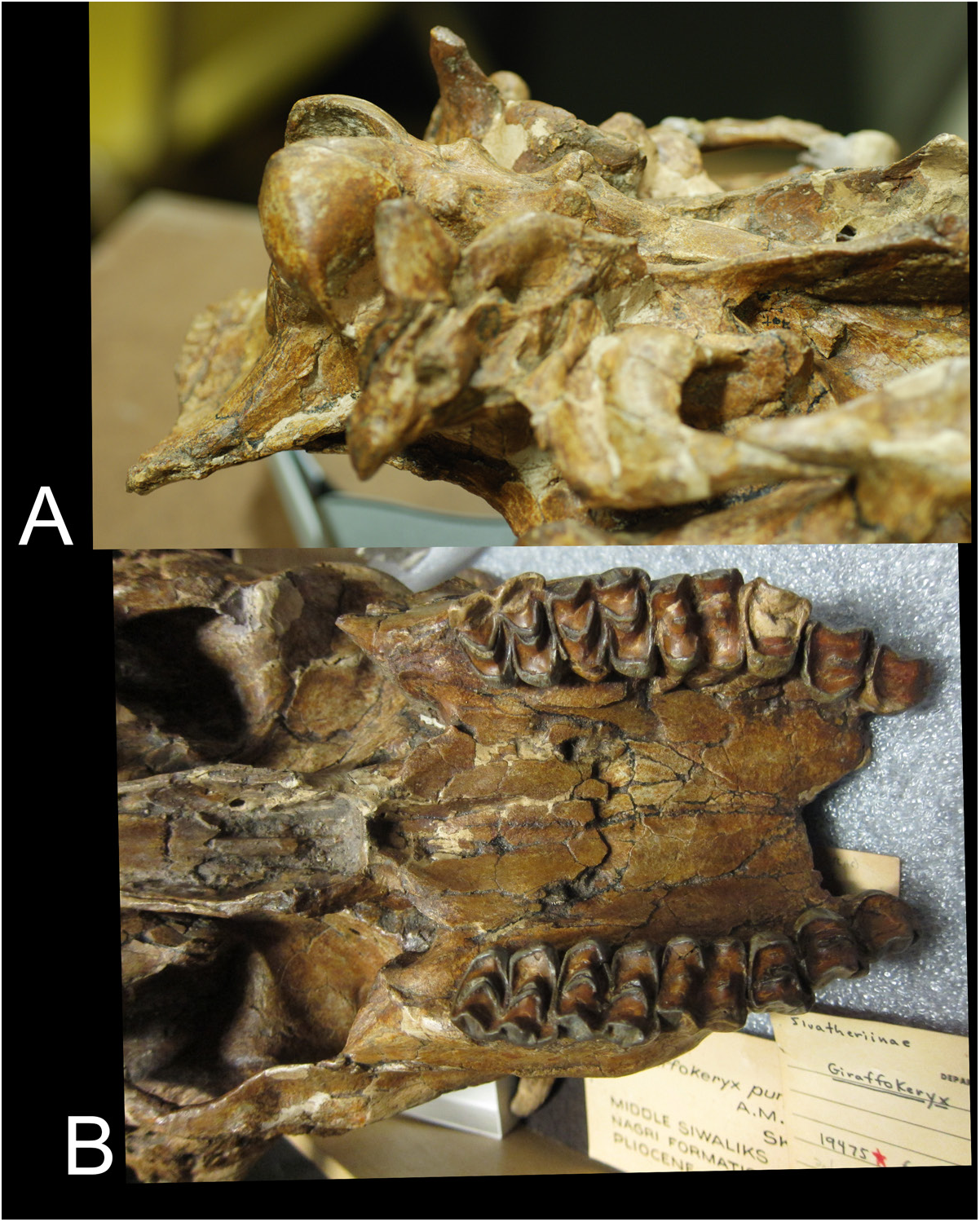
*Giraffokeryx punjabiensis* of lectotype AMNH 19475. A) the basicranium of lectotype; B) the dentition.

*Giraffokeryx primaveus* from Fort Ternan. Metatarsal NHM M30210 (3108). Radius NHM M30215. C3 NHM M3078. The ossicones are lost or misplaced in the KNM Nairobi. We figure them here (based on the originals prior to their loss) (Figure 18). The ossicones are similar to those of *Giraffokeryx punjabiensis*. The only difference so far, the longer premolars in the species from Fort Ternan (Harris et al. 2010). Note that the ossicones are missing from KNM. AMNH 19849 mandible from the Chinji may be primaveus.

**Fig. 18.**
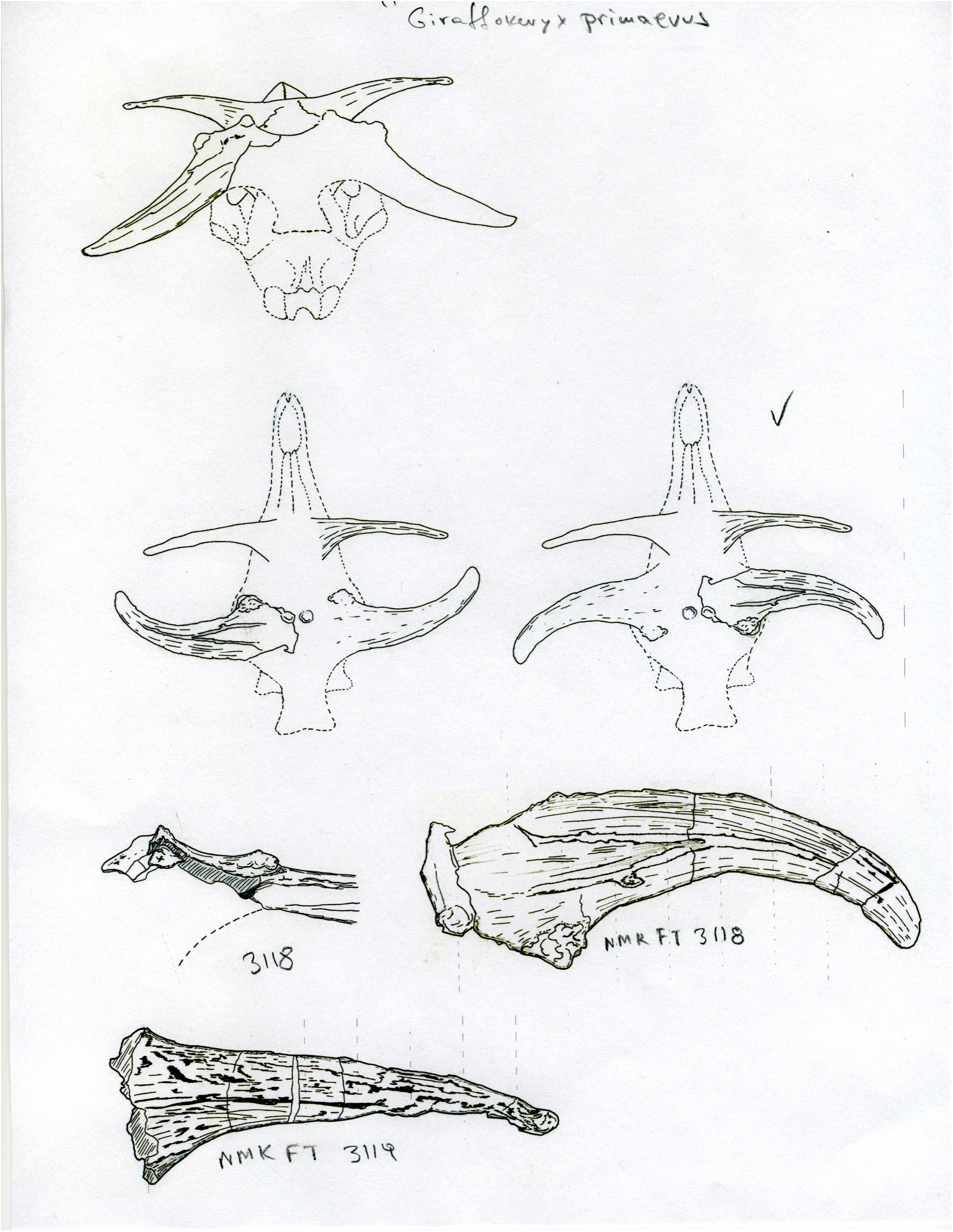
*Giraffokeryx primaveus* from Fort Ternan. The ossicones have been lost in Nairobi – NMK 318 and 319. Restorations of a hypothetical skull. Churcher (1970) thought that the lager ossicone was *Samotherium* and the smaller ossicone was *Palaeotragus*. Both ossicones belong to the same species that had four ossicones.

Okapiinae Bohlin 1926

*Ua pilbeami* Solounias, Smith and Ríos, 2022

Holotype- Y GSP 28483, a partial skull with both ossicones, frontal, ethmoid, and left orbit preserved.

Other specimens- Y GSP 26847 ossicone fragment from locality Y 0691. Type locality- Y0758 (13.940-14.161 Ma), Y 0691 (13.032-13.137 Ma).

Age- 13.032-14.161 Ma.

Diagnosis- *Ua pilbeami* shares with *Okapia* ossicones with an exposed apex and an apical constriction, a large anterior frontal sinus, and a para-style on the upper pre-molars that is deeply separate from the meso-styles. The lower premolar anterior stylids and postero-lingual conids are also deeply separated. It differs from *Okapia* in having long and grooved ossicones. The ossicones are slightly more anterior. In *Okapia* the metapodials are shorter (Figure 19) and (Solounias et al. 2022). Metapodials can be examined in Fig. 20 a and b.

**Fig. 19.**
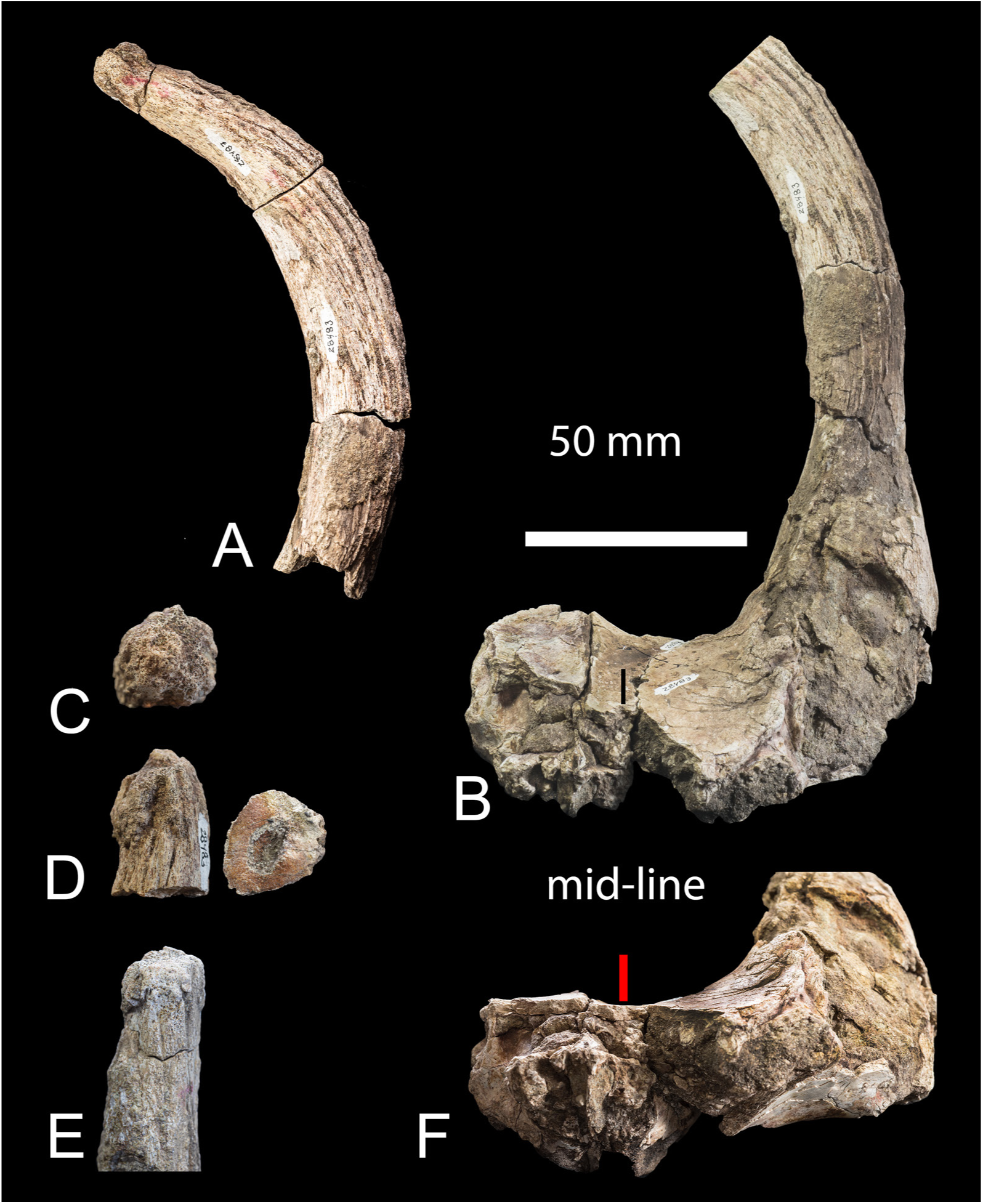
*Ua pilbeami* holotype Y GSP 28483. A) The complete left ossicone. B) View of the skull fragment form an anterior-dorsal aspect to show the frontals and the boss. C) The apex of the ossicone. D) Lateral aspect of the apex with the constriction and a cross section showing the internal canal. E) Medial aspect of the apex. F) A direct anterior aspect of the cranial fragment showing the enlarged frontal sinus. Scale bar: 50 mm for A, B, F; 25 mm for C, D, E.

**Fig. 20.**
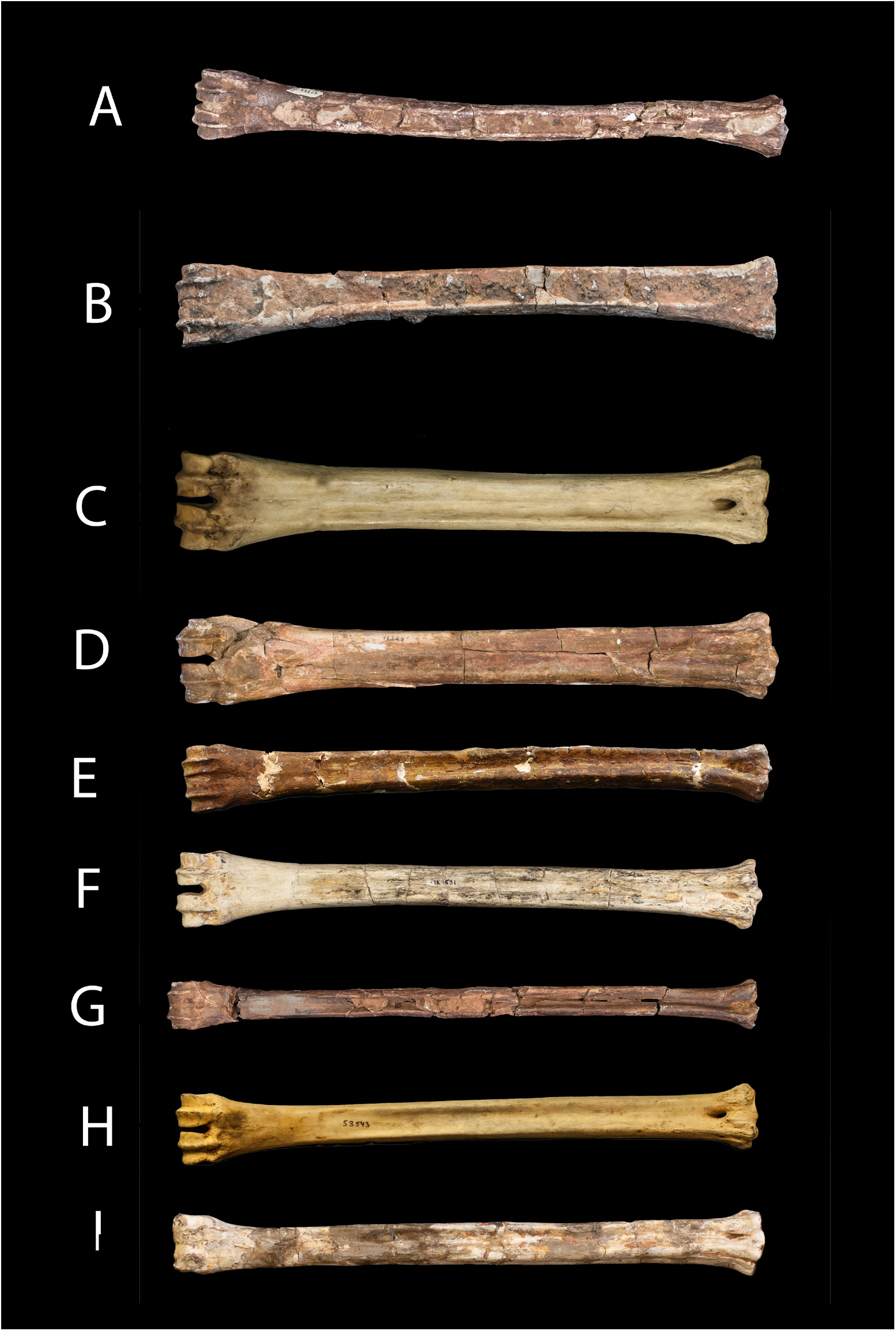
Ventral views of *Ua pilbeami* n. gen. n. sp. metatarsal and metacarpal and other selected metatarsals for comparison. A) *Ua pilbeami* n. gen. n. sp. metatarsal Y GSP 24305. B) Ua pilbeami n. gen. n. sp. metacarpal Y GSP 4505. C) *Okapia johnstoni* AMNH 1196. D) *Decennatherium asiaticum* Y GSP 15184. E) *Giraffokeryx punjabiensis* Y GSP 24306. F) *Palaeotragus rouenii* MNHNP from Pikermi 1691. G) *Orea leptia* Y GSP 20415. H) *Giraffa camelopardalis* AMNH 53543. I) *Bohlinia attica* MNHNP Pikermi 27561. All to the same scale to facilitate comparisons.

Samotheriinae Hamilton 1978

*Injanatherium* Henitz and Sen 1981

Holotype: *Inajantherium hazimi* a skull in Pointers France from Gebel Hamrin Iraq. It was published without a number.

*Injanatherium hazimi* from the Siwaliks YGSP 8869; YGSP 24295; frontal with base of ossicone; YGSP 17891.

Diagnosis – Ossicones rounded smooth surfaced. Apex flat with small apical bumps. Ossicones directed back from a posterior supraorbital position. Braincase similar to *Samotherium boissieri*. Occipital condyles large and approximated. Posterior basioccipital tuberosities flat and transverse. Anterior basioccipital tuberosities rounded and robust. Bulla rounded and small. Glenoid fossa concave. We place this taxon in Samotheriinae.

The new restoration of the type ossicone is a right and is positioned posterior-vertically behind the orbit. The only difference from *Samotherium* is the flat apex of the ossicone (Figure 21 and 22). Heintz and Sen (1981) reconstructed the ossicone as a left and placed it directly lateral to the orbit. In their figure, they do not show the broken edge of the left orbit; a critical oversight. In the same specimen the right orbit is devoid of an ossicone which does not make sense (there should be a right ossicone as well). There is a break in the right posterior orbit where the ossicone was actually attached. This new reconstruction makes *Injanatherium* similar to *Samotherium*. The Siwalik specimens are figured in Fig 22. They are two ossicones and a base of an ossicone on the frontal.

**Fig. 21.**
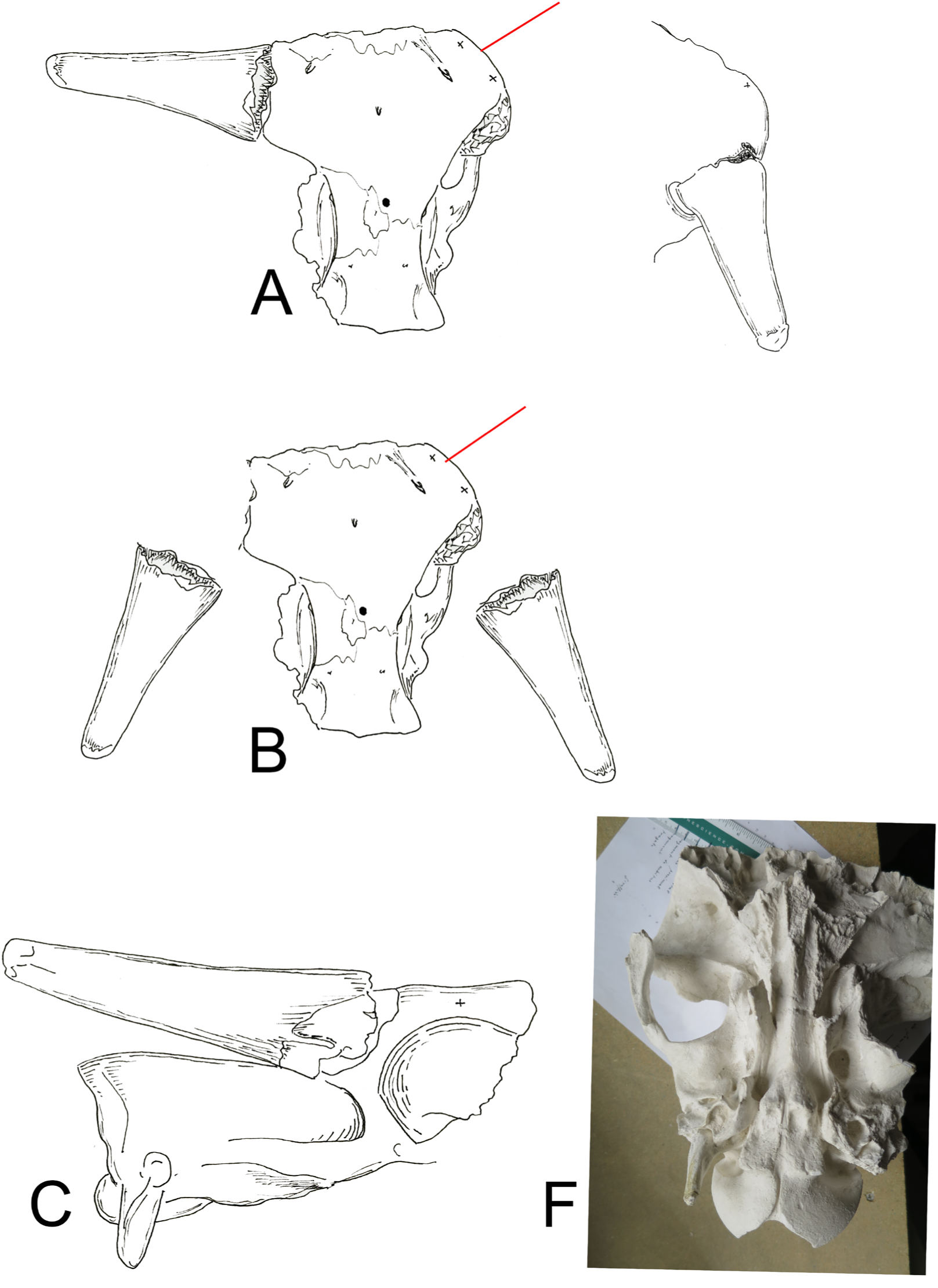
*Injanatherium hazimi* holotype. New restoration. The original *Injanatherium hazimi* specimen. A) the incorrect restoration in the publication. To the right is the correct placement of the ossicone as a right. b) restored ossicones. C) lateral view. Bottom right the basicranium form a cast in MNHNP.

**Fig. 22.**
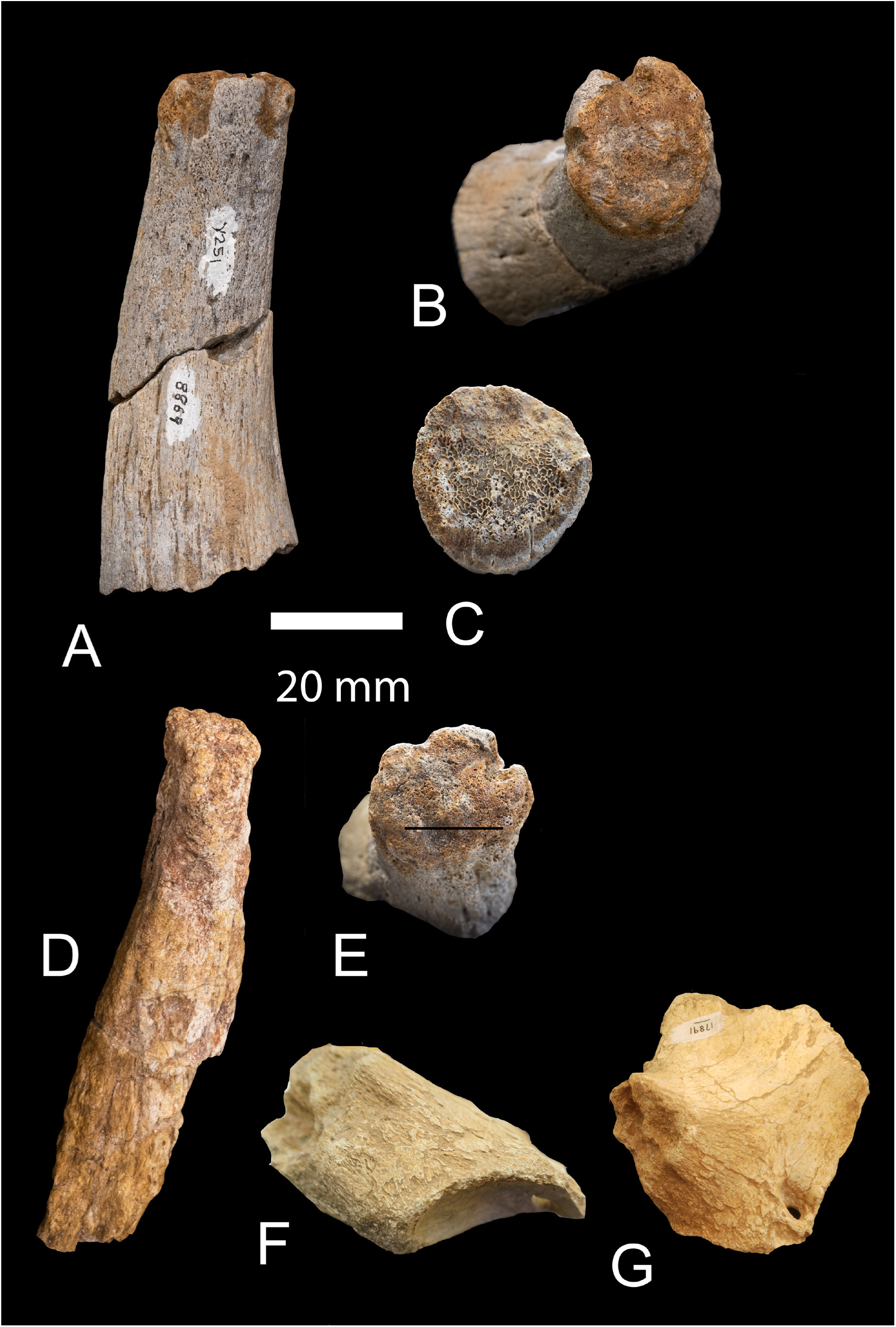
*Injanatherium hazimi* from the Chinji -- a, b (apex) and c sectioned YGSP 8869; d and e) YGSP 24295; f and g) frontal with base of ossicone YGSP 17891.

Palaeotraginae Pilgrim 1911

*Palaeotragus germanii* as in the type Bou Hanifia of Algeria. The Siwalik specimens assigned to this species are: YGSP 16818c, V 5639, radius M0159, tibia YGSP 0985, MT NS 105; MC X 0098. These specimens are placed in *germanii* (Arambourg 1959).

YGSP 16818c, V 5639, Radius M0159, Tibia YGSP 0985, MT NS 105; MC X 0098 are Siwalik specimens referred to *germanii*.

Bohlininae Solounias 2007

*Bohlinia tungurensis* from Tung Gur. New identification of that species as *Bohlinia*. Skull and teeth figured in Colbert (1936). AMNH 26582 is a skull. AMNH 92306 is a metatarsal from Tung Gur. Diagnosis - The parastyle and metastyle of P3 curve inward. They curve in strongly towards the mesostyle. The same character is in *Bohlinia attica* and in *Helladotherium duvernoyi*. Colbert, (1936) described the Tung Gur species as a *Palaeotragus*. The designation of the Tung Gur as *Bohlinia* changes the age of the genus into the middle Miocene (Astaracian). *Bohlinia attica* from Pikermi and Samos is late Miocene (Turolian).

Siwalik specimens: upper premolars YGSP 41413 and 41414.

*Palaeogiraffa macedoniae*

Siwalik specimens: mandible at AMNH from the Chinji AMNH 19323.

*Honanotherium bernori*.

It is known from fragmentary material. Best specimen is a jaw in the AMNH from Dhok Pathan AMNH 19318. Colbert figured this specimen in 1935 fig. 193. And dental specimens: YGSP 27662, 19852, 39136, 20438.

*Visnutherium* sp. and *Vishnutherium psicillum*. Lydekker 1876 pl. VII Fig. 1 and 2 from the Siwaliks of Burma. And Pilgrim 1911 plate II Fig. 17 from the Chinji.

Giraffinae Gray 1821

*Orea leptia* new genus and species

Holotype- Y GSP 20415a – a left complete left metatarsal. Radius Y GSP 20415b distal tibia; Y GSP 20415c malleolar, Y GSP 20415d calcaneus, Y GSP 20415e cubonavicular.

Type locality- Y0640.

Age- 13,649-13,655 Ma.

Etymology- *Orea* is Greek for beautiful – *leptia* to denote its exceptional slenderness and length of the metapodials.

Diagnosis- Large giraffid with extremely slender metapodials and long neck. The ventral metapodial troughs are well-developed. The ventral metapodial ridges are rounded. The distal metapodial end is compact. The malleolar, calcaneus, cubonavicular and astragalus are small and very slender. The astragalus has a head with two facets.

A skull fragment from Al Jadidah in Iraq may also belong to this species. It was erroneously described as *Injanatherium*. There are two metatarsals in that locality which are virtually identical to the holotype metatarsal of *Orea* from the Chinji. Cast: Metatarsal AMNH 127351 B has a deep trough and it is long. Proximal width 38.07 mm; depth 38.07; min width 21.22 mm estimated total length 366 mm. Cast: Metatarsal AMNH 127351 A has a deep trough and it is long. min width 22.97 mm; distal width 40.03. Cast: skull fragment AMNH 127348. The dorsal side of the ossicone base is fattened. There is a posterior ridge- bulge at the posterior edge of the ossicone. There is a similar ossicone in the Chinji. Ossicone is grooved and flat. It has a posterior ridge which is continuous with the temporal ridge. Occipital condylar width 66.08 mm. Minimal distance of temporal lines across calvaria 36.02 (Morales et al. 1987). Holotype: metatarsal YGSP 20416 from Y 640 – Fig. 23 (510X24.1 mm).

**Fig. 23.**
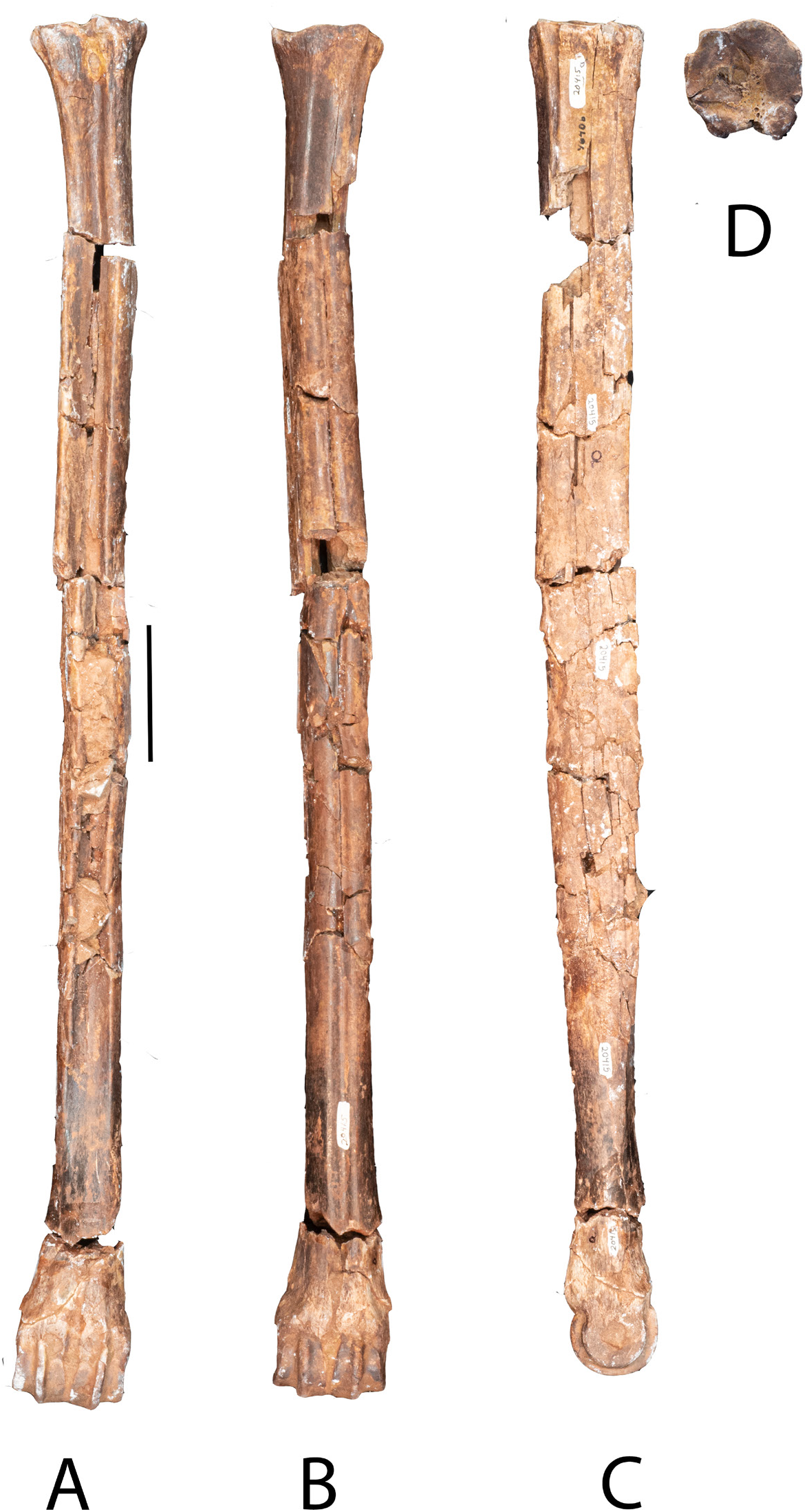
*Orea leptia* YGSP 20415 metatarsal. a) ventral view; b) dorsal view; c) left lateral view; d) proximal end. The holotype is the longest metapodial in any ruminant. From the Siwaliks.

The L: W ratio of the metacarpal is smaller than that of the metatarsal, at 14.9. This is still larger than the L:W ratio of *Giraffa* and *Bohlinia* metacarpals. This L:W highlights the unique slenderness of the metapodials in *Orea*.

Additional Specimens – 37187 left MT fragment, Y 494 12.377 Ma; 4021 left MC fragment Y 076, 11.410 Ma; Y 23694 mid-shaft MC Y 0666, 13.797 Ma; Y 1907 proximal left MC Y076, 11.410 Ma; Y 1168 distal MC Y076, 11.410 Ma; Y 23385 complete radius Y059, 13.62 Ma (520.7×30 mm); Y 23384 complete tibia Y059 13.652 Ma (500X34 mm). Y 20415 cubonavicular, calcaneus and fibula, fragment of astragalus and metacarpal from Y 0640 13.65 Ma. The cubonavicular, calcaneus and fibula are not studied in detail, as we do not have similar known materials from other species to compare in this size range.

The atlas is long as in *Giraffa*. This implies a long neck. The atlas’ dorsal arch and ventral surface are elongated cranio-caudally. The wings are flushed to the body. The L:W ratio of the atlas is 1.39, and its length is 50.61 mm. The distal part of the wing is broken. The foramen for the vertebral artery is oval in this species, similar to the vertebral artery foramen found in *Giraffa*. Fig. 23 shows the metatarsal. Fig. 24 shows additional long limb bones. Fig. 25 shows the atlas of *Orea* and a comparison to *Okapia* and *Giraffa*. The specimens from Saudi Arabia are metatarsals, a skull and ossicone (Fig. 25).

**Fig. 24.**
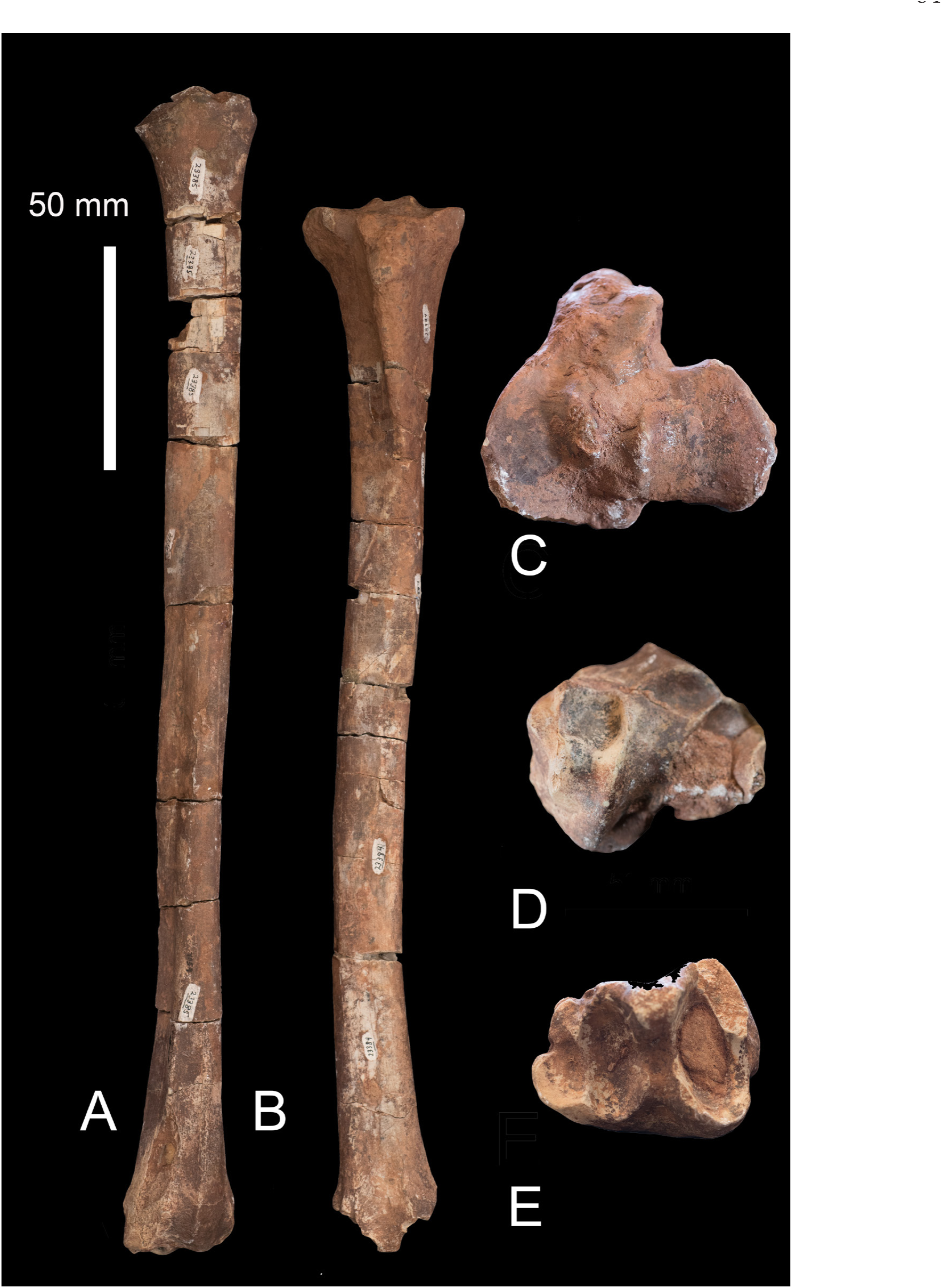
*Orea leptia* a) YGSP 23385 radius; b) YGSP 23384 tibia; c) same tibia proximal end; d) radius proximal end; e) tibia distal end. The radius and tibia. From the Siwaliks.

**Fig. 25.**
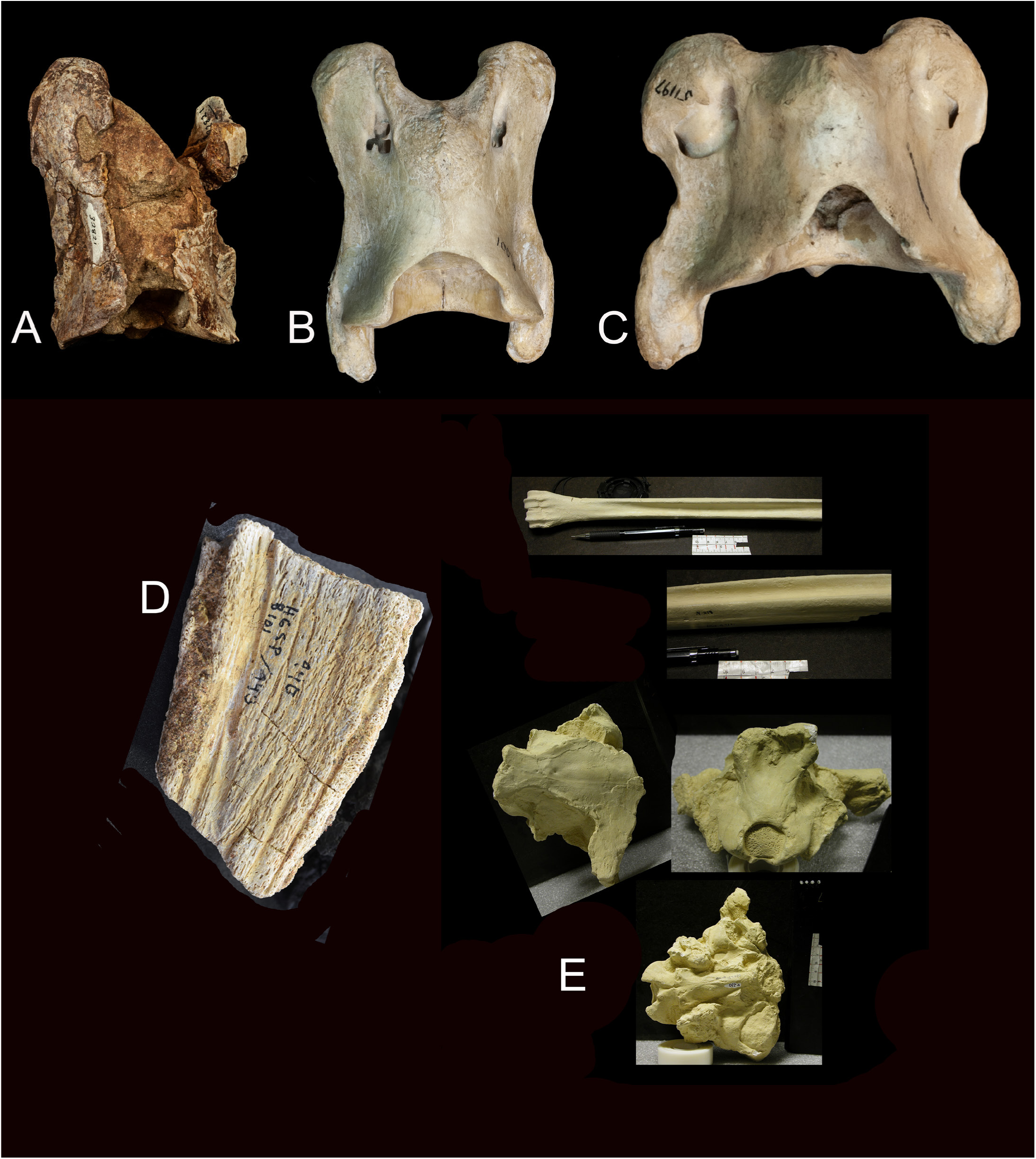
*Orea leptia* A) atlas cervical YGSP 32821; B) *Giraffa* AMNH 82001 C) *Okapia* AMNH 51197. The same scale to facilitate comparisons. The fossil atlas is rather long. D) YGSP 8101/443 an ossicone fragment similar to that of skull AMNH 127340 – from Al Jadidah of Iraq. This skull was named *Injanatherium arabicum* (Morales et al. 1987). D) Metapodials and skull attributed to *Orea leptia*.

Comment: We place *Orea* in the Giraffinae. This makes it the oldest Giraffinae. This taxon could be ancestral to *Giraffa punjabiensis* and to *Giraffa camelopardalis*. *Orea* evolved in the Indian Subcontinent.

*Giraffa punjabiensis* Pilgrim, 1910, 1911.

Holotype- MM no. 20 dentition in IM of Calcutta (never figured).

Type locality-Dhok Pathan

Age-Middle Siwaliks 8.1 Ma

Etymology- *punjabiensis*- from the Punjab.

Diagnosis- Neotype is the ossicone: Ossicone YGSP 16274 was positioned slightly medially and posterior to orbit. No apical knobs and no epikouron on this specimen. The base of the ossicone has small pits on the surface. These differ from those of *Giraffa camelopardalis*. Metapodials are long with ventral trough reduced. Smaller size than *camelopardalis*. P4 is narrower.

Early Late Miocene of the Potwar Plateau, including an 8.1-million-year-old ossicone, 9.4- million-year-old astragalus, and 8.9-million-year-old metatarsal and refer them to *Giraffa*. Left Ossicone: YGSP 16274. We select this to be the lectotype/neotype for the species. The reason is that Pilgrim (1911) did not figure the teeth he refers to this species which are in the IM Indian Museum in Calcutta. Our effort to locate these teeth have been unsuccessful.

Y 20415 cubonavicular, calcaneus and fibula, fragment of astragalus and metacarpal from Y 0640 13.65 Ma. The cubonavicular, calcaneus and fibula are not studied in detail, as we do not have similar known materials from other species to compare in this size range.

*Giraffa punjabiensis* ossicone (Fig. 26 YGSP 16274) in lateral view. (B) *Giraffa sp*. ossicone in medial view. (C) Undersurface of the ossicone. It is likely that this ossicone belonged to a young adult individual, as the undersurface exhibits the characteristic pitting seen in ossicones not yet fused to the skull (Danowitz et al. 2017).

**Fig. 26.**
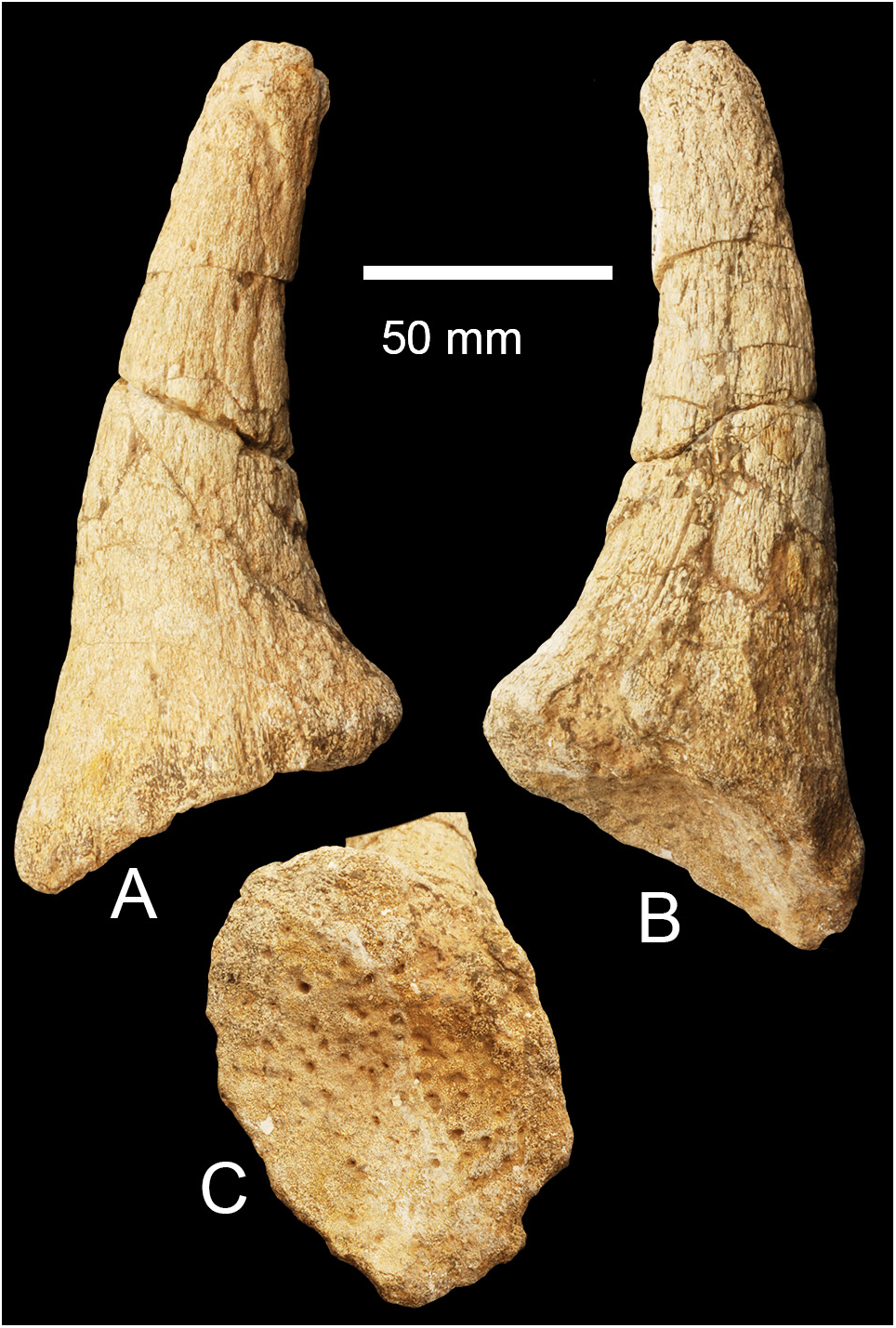
*Giraffa punjabiensis* YGSP 16274. Neotype a) lateral; b) medial; c) base. The attachment differs from that of the okapi and the giraffe.

Measurements: Ossicone 30 mm above base: 50 mm x 35 mm; Length x width of ossicone at apex: 21 mm x 22 mm. Axial length of ossicone: 130 mm. See: Danowitz et al. (2017) (Figure 26).

*Giraffa sivalensis* Falconer & Cautley, 1843

Holotype- cervical specimen figured in Figure 13.

Type locality- Markanda Valley.

Age-Upper Siwaliks – Pliocene.

Etymology- *sivalensis*- from the Siwaliks.

Diagnosis-Molars and premolars very similar to *Giraffa camelopardalis*. Neck long. Metapodials with long ventral deep trough. When compared to *G. punjabiensis G. sivalensis* is larger, has proportionally lower length of upper and lower molars, and a wider p4. Intermediate between *G. punjabiensis* and the more recent *Giraffa*. Figure 13. Van Sittert, and Mitchell (2015). Studied the measurements and proportions of that species. Danowitz et al. (2015) studied the morphometric changes from this species to *Giraffa camelopardalis*. It was found similar but not identical to *G. camelopardalis*. The NHM collection has also metapodials. The collection includes metapodials and isolated dentitions and the cervical vertebra (Figure 27). The teeth are clearly *Giraffa*. The metapodials differ from those of *Giraffa*. They have a deep trough and appear shorter. They may be a different taxon mixed in the sample. If they belong to the same taxon, a new genus is needed. They may be *Honanotherium*.

**Fig. 27.**
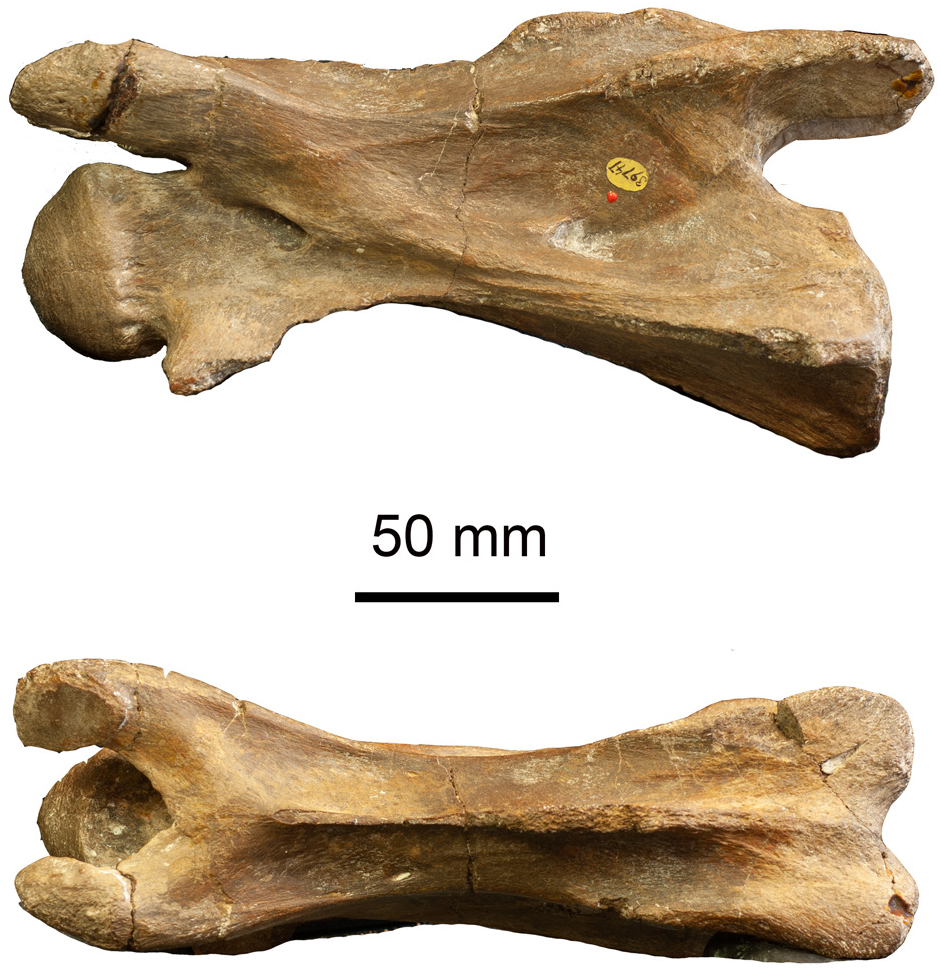
*Giraffa sivalensis* a C3 vertebra NHM M 39747.

#### Bramatheriinae new rank

A type species for the new rank is *Baramtherium perminese* at the AMNH. It includes metacarpals metatarsals a jaw, a skull and many small fragments and teeth, deciduous dentition. We remove *Sivatherium* in its own group.

*Lyra sherkana* Ríos & Solounias, 2023.

Holotype- Y GSP 47357, braincase with both ossicones, basicranium and occipital (face and dentition missing).

Paratypes- Detached ossicone fragments (YGSP 6392- loc. Y0311, YGSP 47192-loc Y0514, YGSP 49597-loc Y0311, YGSP 20651, loc-Y0311).

Type locality- Y0647.

Age-12.352-12.402 Ma.

Etymology- The genus is named after lyra an ancient instrument with similar shape, the specific name it’s in honor of Dr. Tanya Sher Khan daughter of the woman medic assigned in Khaur. She participated in the campaigns of the Harvard Geological Survey of Pakistan Project since she was a little girl and found the holotype during the 1994 expedition.

Diagnosis- Giraffid with ossicones positioned slightly posterior to the orbit, with the anterior ossicone edge reaching the midpoint of the orbit. The ossicones are slightly posteriorly inclined in lateral view. The ossicone surface is composed of two layers, a thinner external layer and thicker internal layer. The posterior ossicone surface is lumpy with streaks of epikouron. The base is smooth and devoid of epikouron. There is a bulge at the base both in the posterior and the anterior side. The shaft is gently curved posteriorly, directed laterally forming a 50- degree angle with the skull roof, and is triangular in cross section. A slight clockwise torsion is present along the axis of the shaft. The apex terminates in a small knob that can be smooth or with small bumps present on its surface. The para-apex is present. The frontal sinus and surface boss penetrate the base of the ossicone. There is a faint posterior keel. There is a small bulge between the two ossicones centrally on the skull. The frontal sinuses are developed anteriorly but are not well developed on the caudal part of the braincase. The braincase is short. The posterior margin of the right supraorbital foramen is situated in the middle of the antero- medial surface of the ossicone. The anterior basioccipital tuberosities are notably large. The mastoids are covered by bone thickenings resembling epikouron. The anterior basioccipital tuberosities are notably large and rough on their surface. The bulla is bulbous and rounded. The occipital condyles are well separated on the ventral surface. Figures 28, 29, 30. Dimensions: See Ríos & Solounias (2023).

**Fig. 28.**
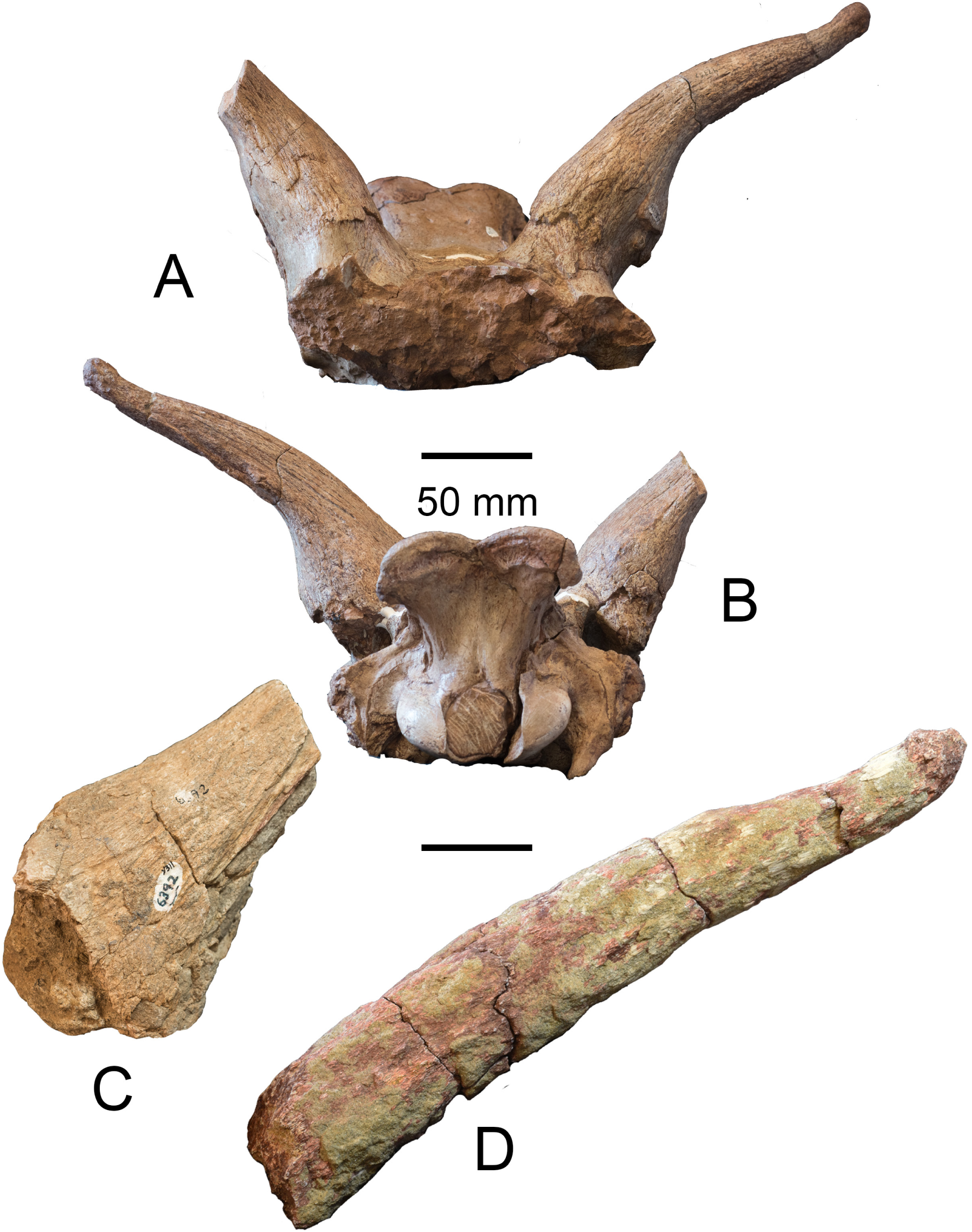
*Lyra sherkana* Three ossicones. a and b) YGSP 43357 c) YGSP 6392; d) 49597.

**Fig. 29.**
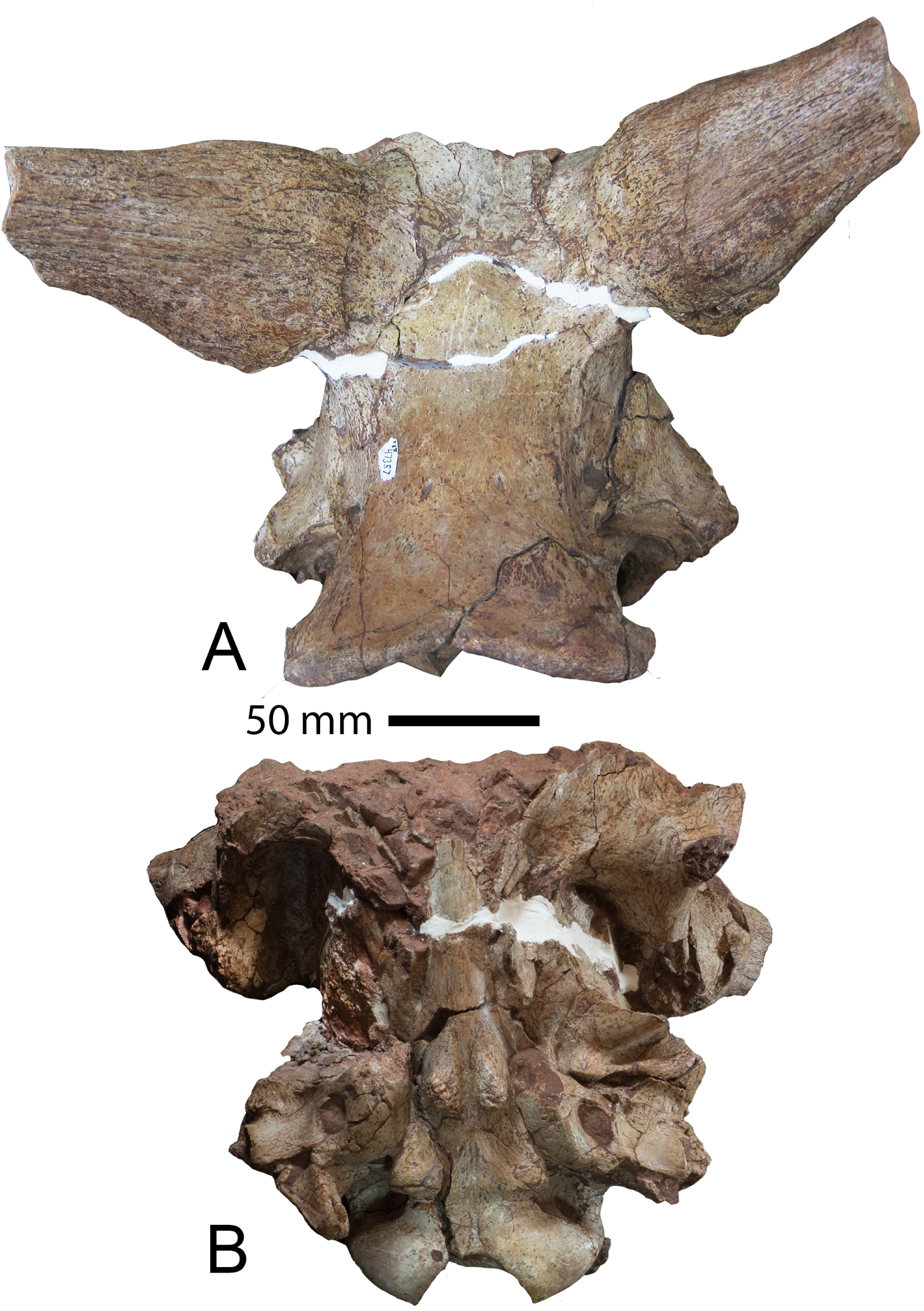
*Lyra sherkana* The holotype. a) dorsal view; b) braincase view of YGSP 43357.

**Fig. 30.**
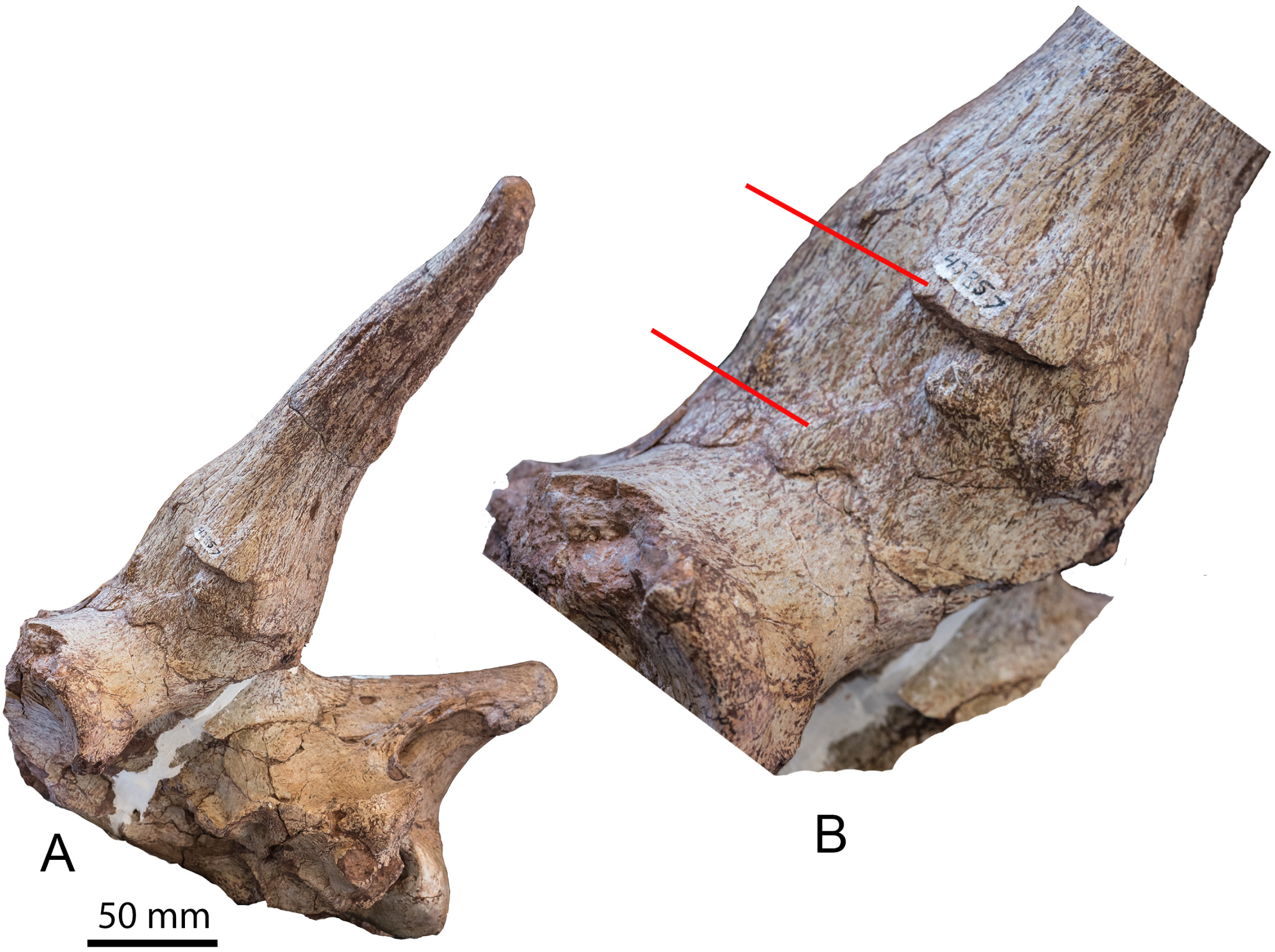
*Lyra sherkana* The holotype. a) lateral view; b) Close up of ossicone of YGSP 43357 showing massive boss and two surface layers of ossicone material.

*Helladotherium grande* a skull figured by Pilgrim 1911. It is an occurrence of a Pikermian Biome taxon also present in the Siwaliks (Pilgrim 1911).

*Decennatherium asiaticum* Ríos, Danowitz & Solounias, 2021

Holotype- metatarsal YGSP 15184.

Paratype- metacarpal YGSP 11455.

Type locality- Y0311.

Age- 10.026-10.100 Ma.

Etymology- *Decennatherium*, beast of the deity Decenna; *asiaticum*- from Asia.

Diagnosis- Large giraffid. Curved ossicones ornamented by a high number of deep ridges on their surface, which run longitudinally to the apex of the ossicone. The apex is blunt, and the section is oval. Smaller than the ones of *Decennatherium rex*. Lower molars longer than wider where the lingual wall shows a developed metastylid. The metacarpal is longer and with a slightly slenderer diaphysis index than the other *Decennatherium* species with a value of 9.18. The lateral and medical epicondyles of the metacarpal are slightly asymmetrical with a more triangular medial epicondyle. The metacarpal palmar shaft shows rounded medial and lateral ridges. Distally, the tibia shows a large, flat, and circular ventral fibular facet and an elongated and thin medial malleolus. The distal APD/distal TD tibia index is lower than in the other Decennatherium taxa with a value of 0. 74. The astragalus intratrochlear notch is broad and shows a distinct medial scala and distal intracephalic fossa. The metatarsal shows no distinct pygmaios and has a slightly slenderer diaphysis index than the other Decennatherium species with a value of 8.44. The proximal phalanx shows a distinct notch in lateral view that separates the lateral palmar eminence from the articular surface of the base. Figure 28. A detailed study by Ríos, Danowitz & Solounias (2021) relates to this species (Figure 31).

**Fig. 31.**
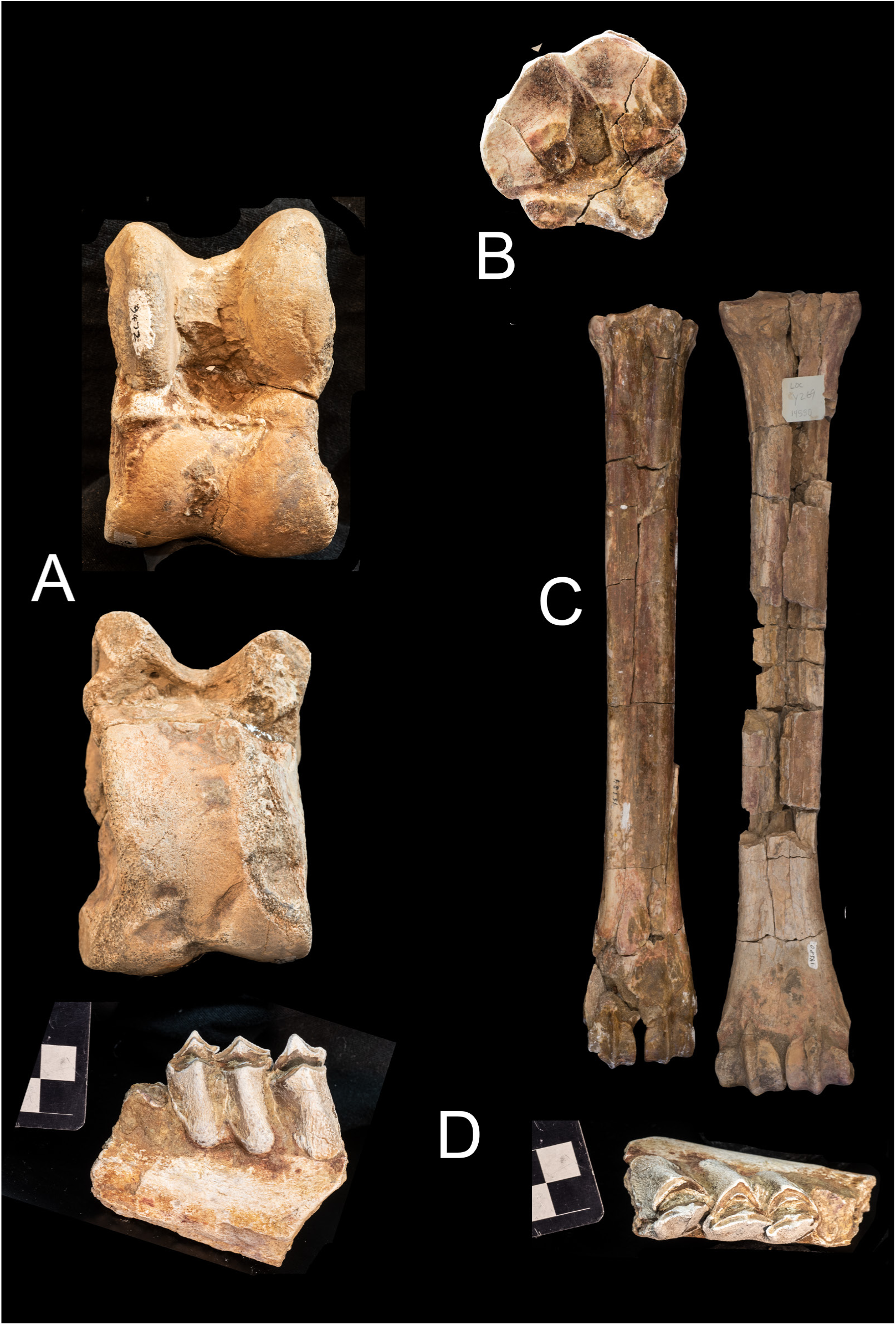
*Decennatherium asiaticum* a) YGSP 9472 astragalus; b) YGSP 15184 MT; c) YGSP 14580 MC; d) m3 tooth YGSP 13201. Scales in cm.

*Libytherium proton* Ríos M., Abbas S. G., Khan M. A, and Solounias N., 2022.

Holotype- Y GSP 46530, complete left ossicone.

Paratype- Y-GSP 49608, right hemimandible.

Type locality- Y0311.

Age- 9.596-10.100 Ma.

Etymology- *Proton* means the first.

Diagnosis- A notably large giraffid. The ossicones are cylindrical, elongated, and curving inward but not twisted. There is lateral prong near the base and no base flange. The cross section is oval near the base. The shaft is rounded and has no bumps. The ossicone extends upwards and curves gently. There is fine grooving on the surfaces (dorsal and ventral). Near the apex the there are two small dorsal bumps followed by a concave excavation. There are also two ventral bumps. The area where the bumps begin is the para-apex. Other ossicones do not have bumps. The apex is a small, flattened disc. There is a large central canal. The jaw is large and robust and with a short symphysis. There is no indentation on the jaw between the angle and ramus. The molars have a small buccal cingulum and reduced styles. The p3 has a mesolingual conid. The posterolingual and posterobuccal conids join posterolingually through the posterior stylid forming a rounded posterior fossette. The p4 is very molarized with a larger anterior lobe that represents two thirds of the tooth length, and the protoconule is labially more extended than the protocone. Enamel is very rugose and thick. The entoconid is separated from the metaconid and the posthypocristid. The protoconid is wider and rounder than the hypoconid, which is U-shaped. There is a thin buccal connection between the wide postprotocristid and the prehypocristid. The radius is large, with a large bulge on the dorsal surface distally above the lunate fossa. The styloid process of the ulna is fused to the radius and is small and round. There is a small depression dorsally for the extensor tendon. The metacarpals are longer than in *Libytherium maurusium* and *Sivatherium giganteum*. The central palmar trough is shallow. The astragalus is not squared. It is a simple rectangular in outline with a small medial crest on the trochlea (Figure 32. Fig. 35 A, B, and C).

**Fig. 32.**
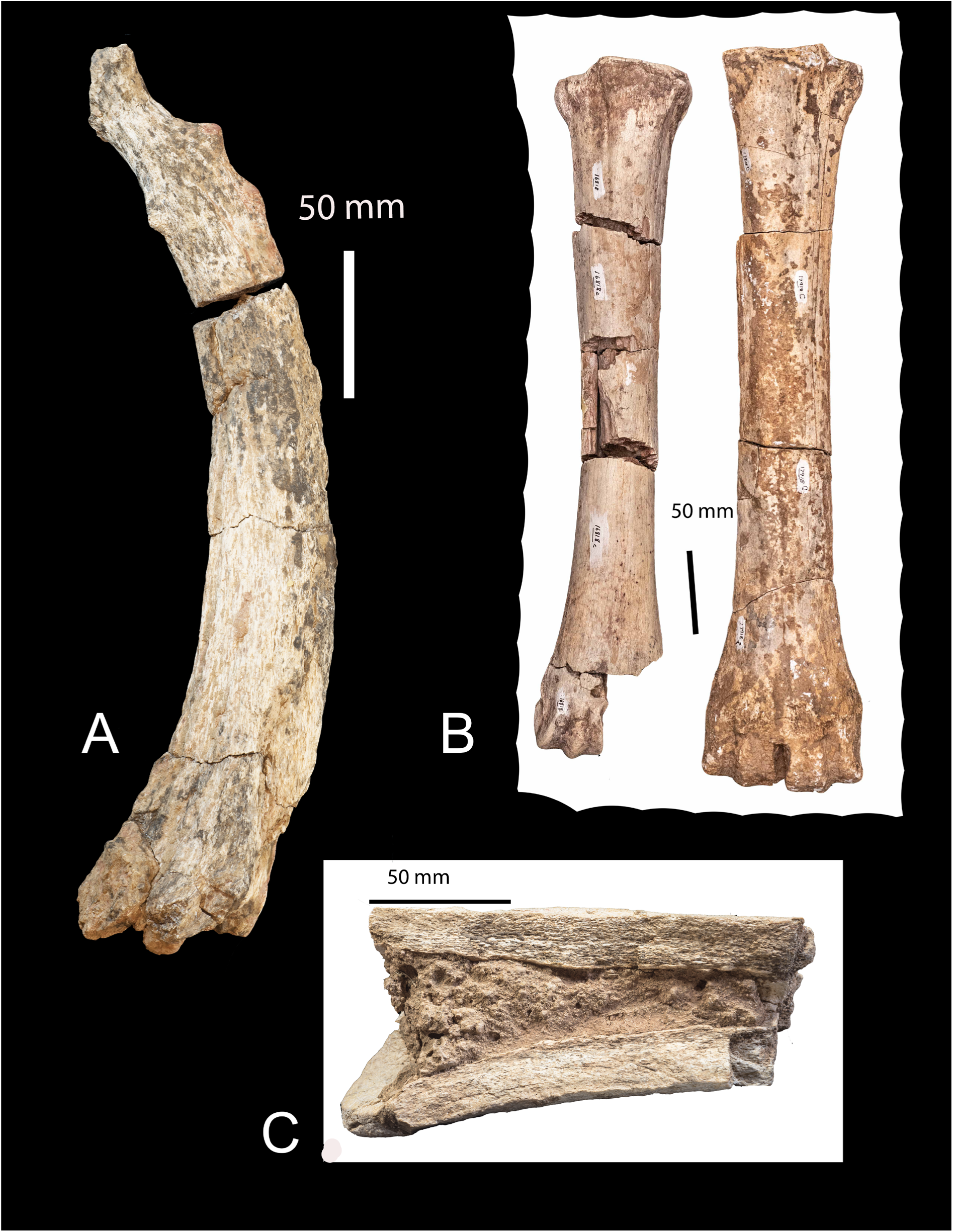
*Libytherium proton* a) ossicone YGSP 46530 holotype; b) Metacarpal 16818; and metatarsal 17918. c) YGSP 14960 ossicone fragment showing inner canal, inner cortex and thick outer cortex. Scales in mm.

**Fig. 33.**
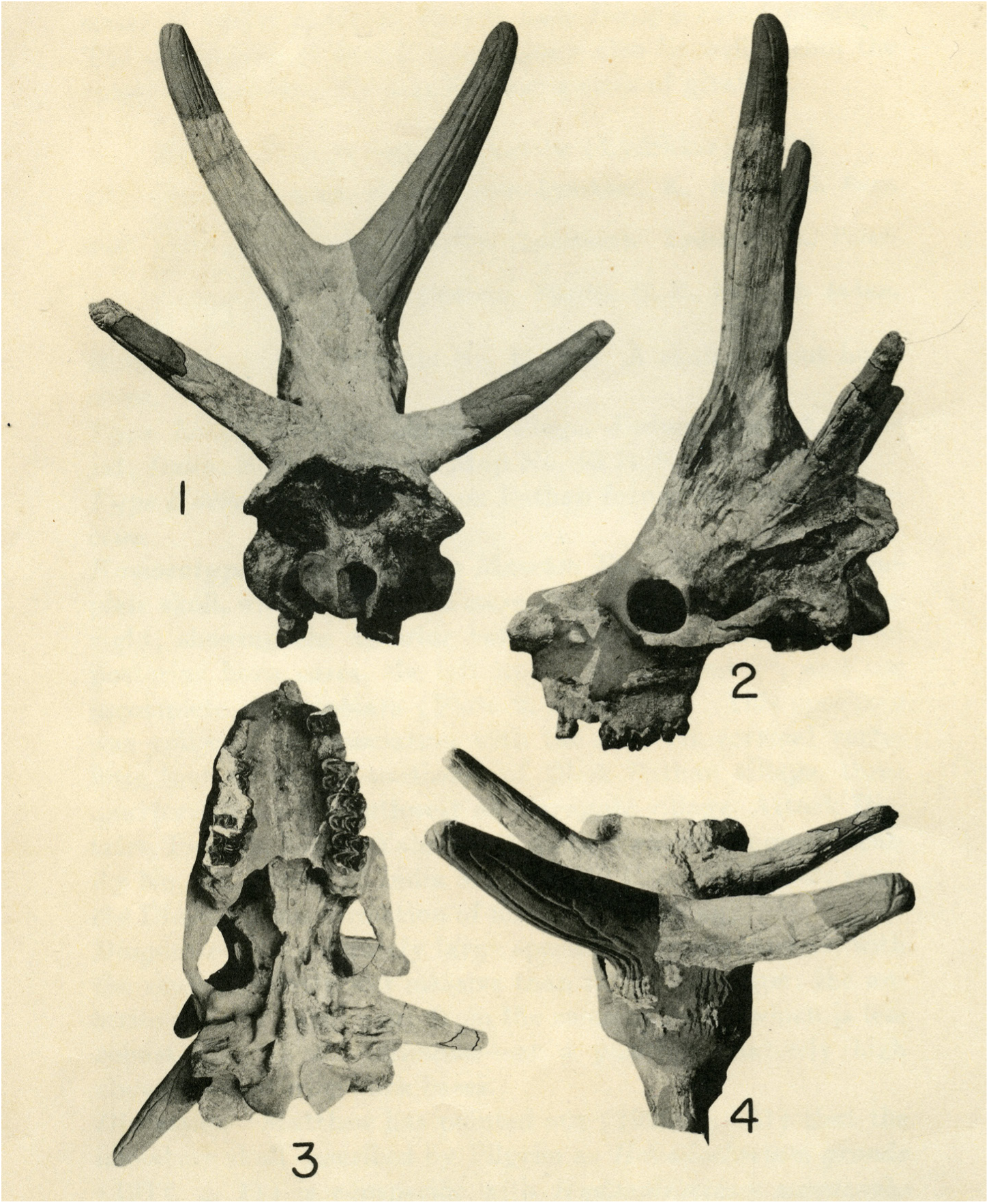
*Bramatherium megacephalum* skull at YPM 13881 - figure from Lewis (1939).

**Fig. 34.**
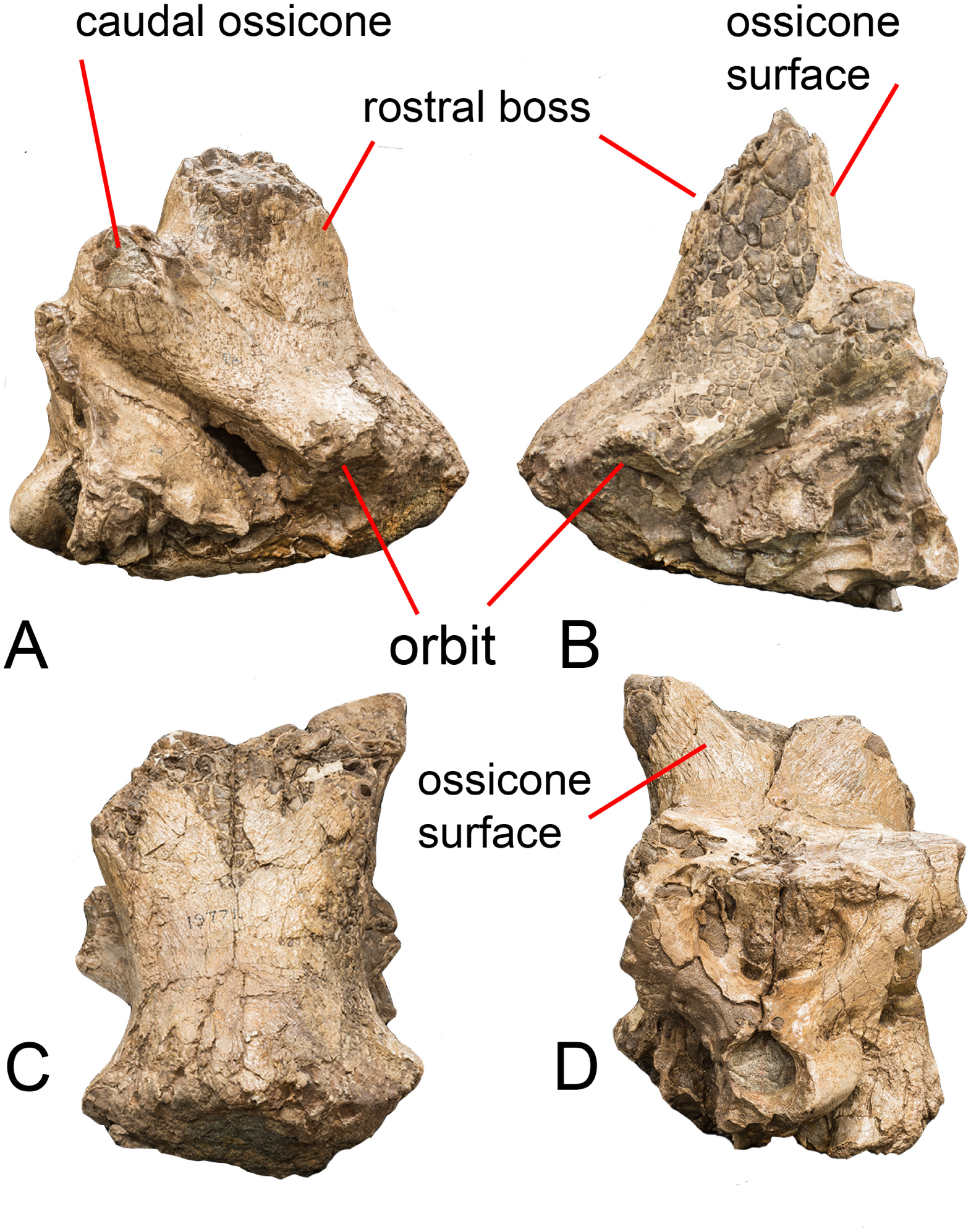
*Baramtherium perimense.* Skull from the Perim Island Pliocene AMNH 19771 (Colbert 1935). A right lateral. B) left lateral. C anterior. D) occipital.

**Fig. 35.**
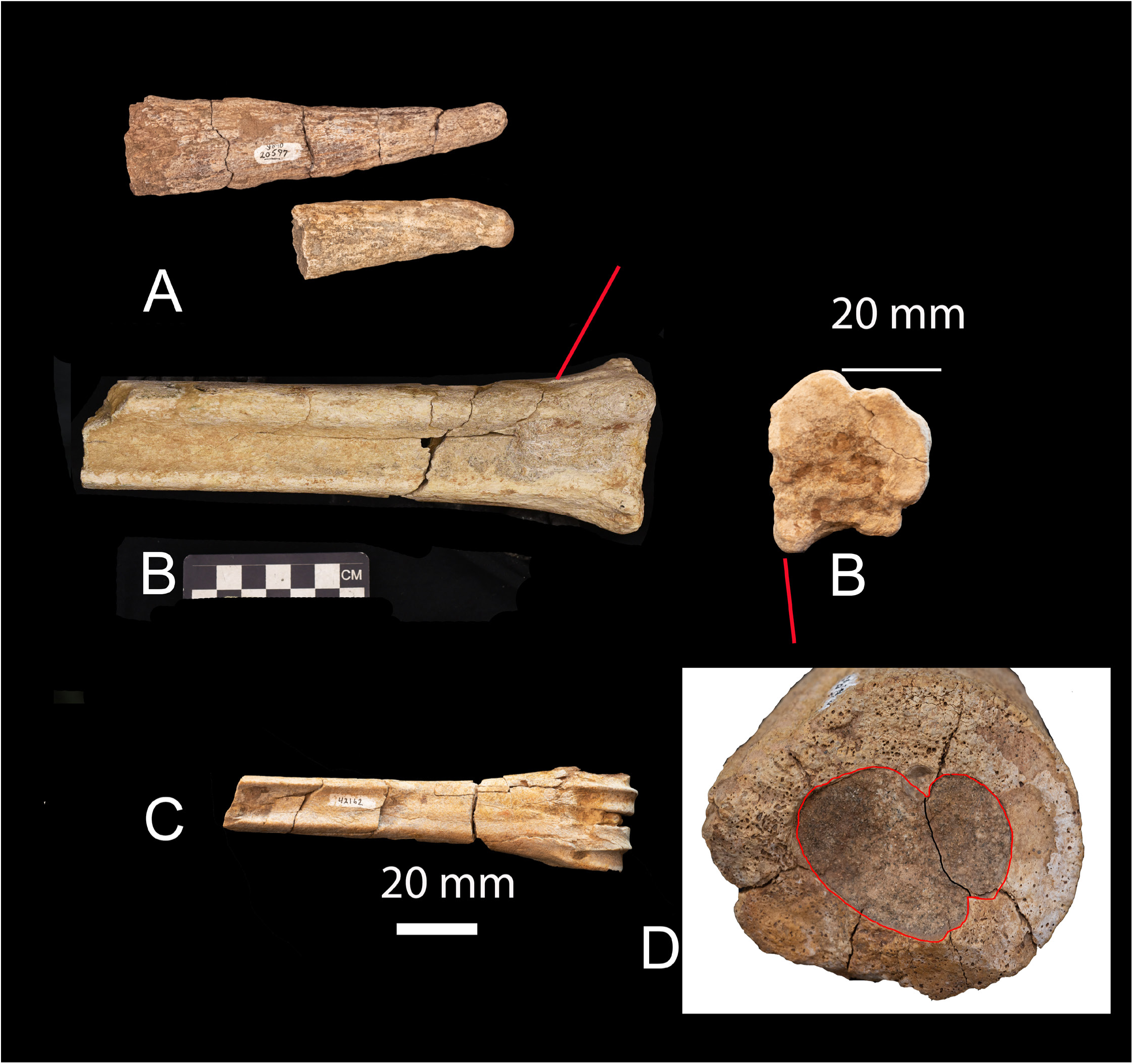
*Bamiscus micros*. A small-sized Bramatherium-like taxon. A) ossicones YGSP 20597; B) Metatarsal YGSP 14998 – red lie shows the characteristic medial large ridge; C) Metacarpal YGSP 42162. D) Bramatherium sp. Ossicone with two internal canals separated by a thin septum (black line). The canals are outlined by a red line. We need the type of Bramiscus

Dimensions: See Ríos, et al. (2022)

*Bramatherium megacephalum* Lydekker, 1876

Holotype- AM-19670, skull.

Type locality- Dhok Pathan.

Age- 7.0-10.2 Ma. Late Miocene.

Holotype IM D 150 skull (Lewis 1939).

Etymology- *Bramatherium*, beast of the Hindu deity Brama; megacephalum, large head, regarding the large ossicones.

Diagnosis- Large-sized giraffid with robust limbs, a short face and a massive and relatively short neck and rugose tooth enamel. *Bramatherium* is further distinguished by two very large, transversely placed ossicones arising from a common, median frontoparietal base and two smaller, more posterior, lateral, parietal ossicones. When compared *to B. perimense,*

*B. megacephalum* has less robust and smaller ossicones, with the anterior ossicones arising from a less developed common base and a less massive skull, plus less prominent anterior styles in the upper teeth. Described and figured by Lewis 1939.

Dimensions of the Yale specimen described by Lewis 1939: width across occipital condyles 137 mm; P3 to M3 179 mm. Neck lengths: atlas 90 mm; axis 192 mm; C3 140 mm; C4 142 mm; C5 160 mm; and C6 163 mm (Figure 33 and Figure 35 D).

*Bramatherium perimense* Falconer 1845

It was collected in the Pliocene sediments of Perim Island. Type skull figured but not named by Bettington (1845). Skull presumably still in the Royal Society of Surgeons collections. Cast of that skull at AMNH is 27016 (according to Colbert 1935; lectotype is NHM maxilla 48933 – not diagnostic). Second skull from Perim Island AMNH 197791 from Dhok Pathan figured in Colbert (1935). Massive anterior ossicone bosses. The actual ossicone was thin over the huge nasofrontal boss. See Colbert (1935) for figures and more information.

Dimensions described by Lewis (1939) and Colbert (1935): width across occipital condyles 131 cm. P4 to M3 163 mm; P3 to M3 168; See Lewis (1939) (Figure 34).

*Bramiscus micros* Ríos M., Abbas S. G., Khan M. A, and Solounias N., (in press)

Holotype- PUPC 22/01, a small fragment frontlet bearing pair ossicones with their fused bases.

Type locality- Chinji Formation, Lower to Middle Siwaliks, Punjab, Pakistan.

Age- 9.3-13.65 Ma.

Etymology- *Bramiscus*: -*iscus* means very small in Greek, means small *Bramatherium*.

*Micros*- means small in size (as in meniscus and *paniscus*).

Diagnosis- Diagnosis. Small size giraffid with two pairs of ossicones. The anterior pair are fused at their base (Bramatherium-like). The posterior pair is straight and possibly directed laterally. All ossicones have an oval cross-section at base and tending to be circular at apex. The ossicone surface is covered with longitudinal ridges of a medium depth. The ridges are deeper and larger ventrally and laterally. The dentition is brachydont with rugose enamel. The upper molars have a postmetaconule fold and a labial cingulum, as well as a developed entostyle. The lower p2 has a crest on the anterior valley. The lower p3 has a well-developed paraconid and parastylid. Metaconid narrower than the entoconid. Lower molars with protoconid and hypoconid slightly rounded and labially has rugose enamel and an anterolabial cingulum. *Palaeomeryx* fold absent. Lower molars with high L/W index. Metatarsal has a characteristic large medial ridge and a low lateral ridge, Figure 35.

Dimensions: See in Ríos et al. in press.

Sivatheriinae Bonaparte 1850

*Sivatherium giganteum* Falconer & Cautley, 1843

Holotype- NHM 15283 skull. (Falconer and Cautley, 1836: plate 2; Harris, 1976a, 1976b); ossicone NHM: 39524 a; ossicone 39525; 39523 female skull; 39533 metacarpal; 39752 metatarsal and a complete neck.

Type locality- Upper Siwaliks Age-Plio-Pleistocene.

Etymology- *Sivatherium* means beast (therium) of Siva, the Hindu god of destruction, and giganteum refers to its large size.

Diagnosis- Skull where the posterior ossicones are composed of hundreds of placations and are covered with ribbons of material – not palmate but with a large bump near the base. Anterior ossicones simple cones. Nasals arched. The frontal sinuses are small but multiple. The molars resemble bovines with additional crests. The metacarpals are shorter than the metatarsals. The neck is short with massive ventral tubercles. Figures 33 and 34. This is a new concept of a family monotypic for *Sivatherium* (Figure 36, 37). In summary, *Sivatherium* is a rather unique taxon. It has thousands of placations in the back “horn.” The placations are horizontal and are covered with broad ribbons of bony material. The metacarpals are shorter than the metatarsals unlike any other ruminant. In that study we show that the “horns” of *Sivatherium* differ strongly from all other Giraffidae (Rios et al. 2022), (Fig. 36 and 37).

**Fig. 36.**
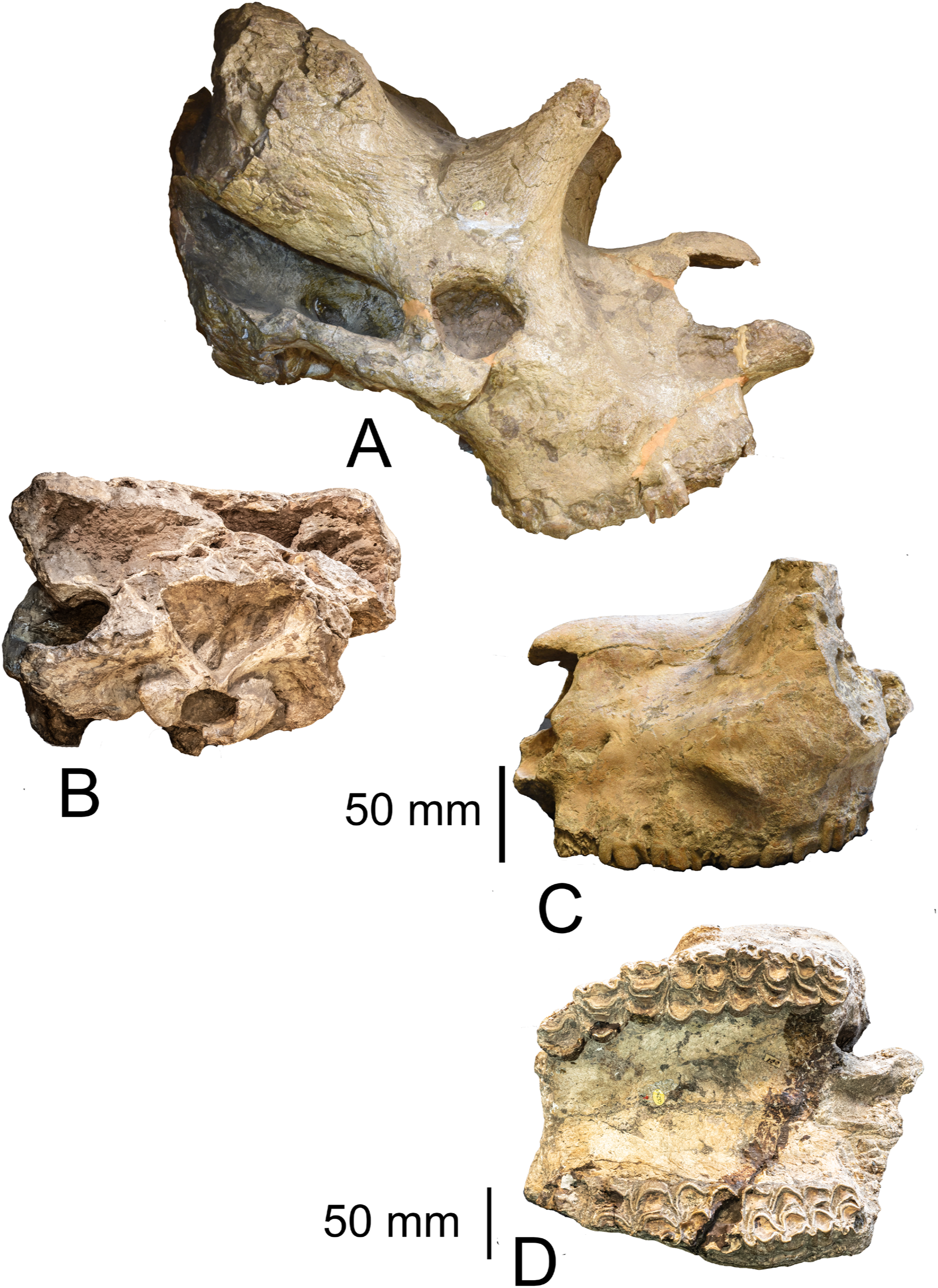
*Sivatherium giganteum*. A) holotype NHM M 39524; B) occipital view; C) NHM M 15284 lateral view; D) same in palatal view. Scales in mm.

Figure 38 shows the proximal metatarsal of six species. Differences in *Giraffokeryx* are evident.

**Fig. 37.**
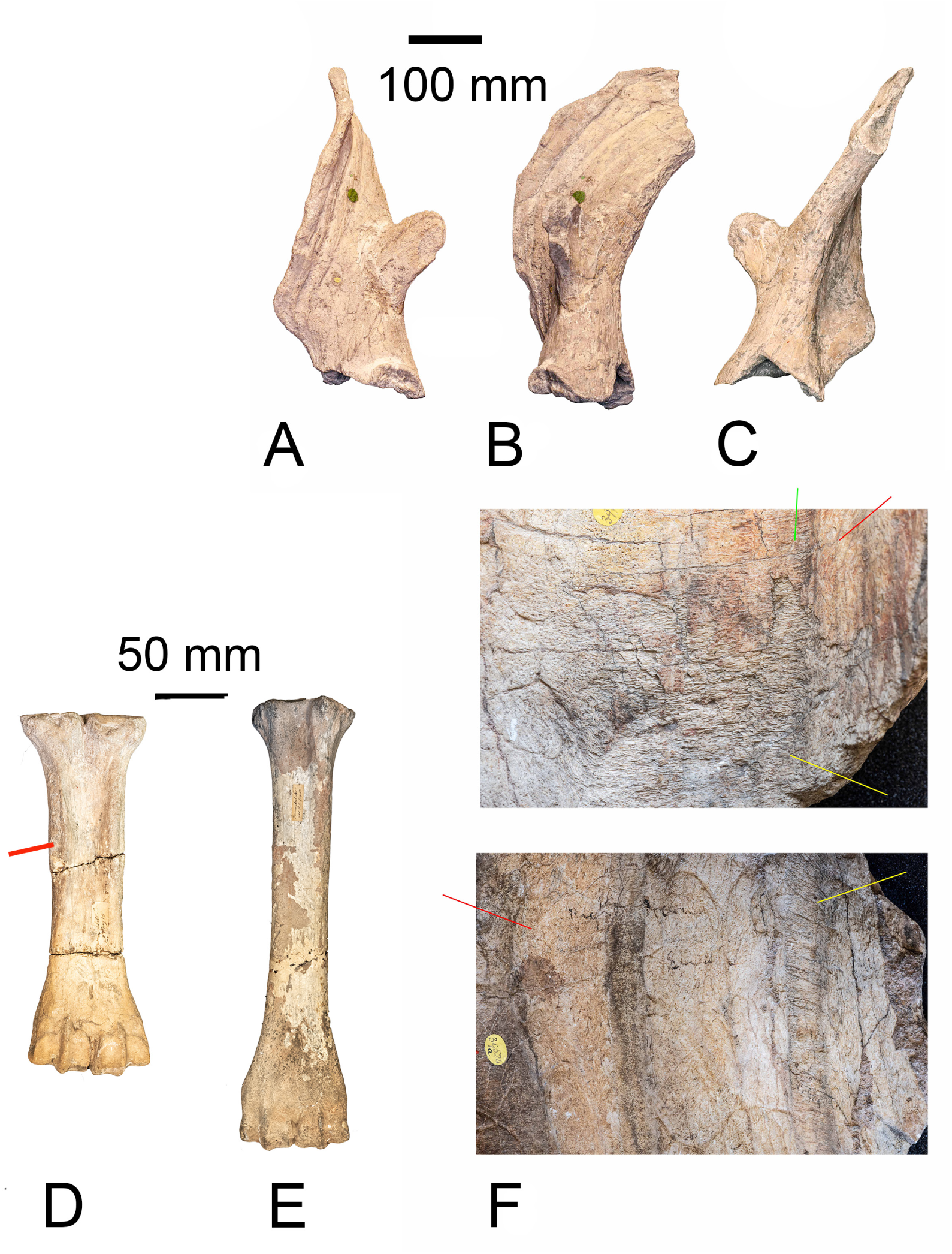
*Sivatherium giganteum.* Best ossicone. A) (anterior), B) (posterior), C) (lateral), of ossicone NHM M39525; D) MC NHM M 39533; E) MT NHM M 39752. F) Close up of an ossicone specimen that shows thousands of internal placations. Ribbons cover the placations.

**Fig. 38.**
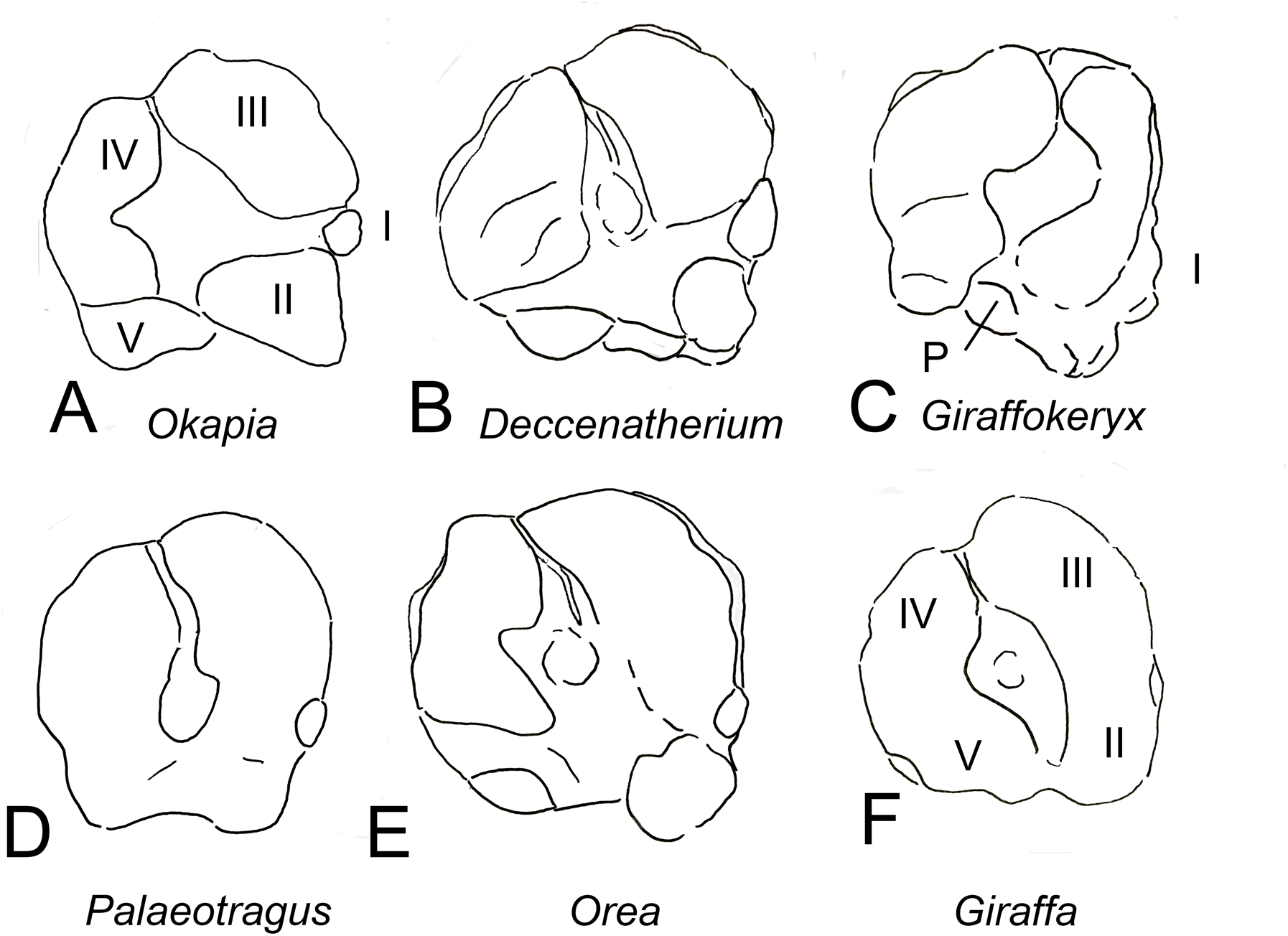
A variety of proximal ends of metatarsals showing the presence of five digits in all. *Giraffokeryx* has an untypical digit III.

## DISCUSSION

The Siwaliks Miocene has the highest diversity in giraffoid taxa of all the fossil record, only similar to the diversity peak found in Greece during the Late Miocene (Solounias 2007; 2023). It has recorded a variety of giraffoids including all three families of the clade: Palaeomerycidae, Climacoceridae and Giraffidae. The earliest recorded are the climacoceratids, which disappear from the Siwalik faunas before the Middle Miocene. Palaeomerycids appear for the first time in the Siwaliks record at the beginning of the Middle Miocene disappearing before the Late Miocene. Giraffids appear first in the Lower Miocene and achieve the highest diversity during the Late Miocene, after, they gradually disappear from the Siwaliks.

### Lower Siwalik Giraffoid faunas (18.3-13.1 Ma)

The earliest giraffoids recorded are the climacocerids *Vittoria soriae* nov. gen., *Goniomeryx* flynni gen. nov. sp nov. and *Orangemeryx badgleyi* nov. sp nov., and the giraffid *Progiraffa exigua*, *Ua pilbeami*, and *Orea leptia*. The earliest palaeomerycid of the Siwaliks record is *Goniomeryx flynni* nov. gen. nov. sp. followed by *Nuchalia gratia* nov. gen. nov. sp.

### Middle Siwaliks Giraffoid faunas (13.1-10.1 Ma)

The majority of plaeomerycids appear during the Middle Siwaliks, including *Lateralia morgani* nov. gen. nov. sp., *Tauromeryx canteroi* nov. sp., and *Fovea fossata* nov. sp. They coexisted with several giraffids including *Giraffokeryx punjabiensis*, *Giraffa punjabiensis*, *Lyra sherkana*, and *Bramiscus micros*.

### Upper Siwaliks Giraffoid faunas (10.1-0 Ma)

The latest giraffoid faunas included the last palaeomerycids which become extinct right at the beginning of the Upper Siwaliks. The main representatives of the clade were the large Sivatheriinae giraffids *Bramatherium megacephalum, Bramatherium perimense*, *Libytherium proton*, and *Sivatherium giganteum* plus the giraffine *Giraffa sivalensis*.

## THE GIRRAFFOIDEA GENERAL CONSIDERATIONS

There are several families of Pecora. They have been generally classified by Janis and Theodor (2014). They are: The Cervidae and primitive cervoids have been most extensively studied (e.g., Azanza et al. 2011; Landette-Castillejos et al. 2019; Rossner et al. 2021; Rossner 2010). There is basically a persisting scientific interest in antlers their histology, rapid growth and antler shedding. Antlers generate above a frontal pedicle and are annually shed. The general result is that cervids have the same type of antler from the beginning of the family. Bovidae have been extensively studied in both systematics as they include cows and sheep which are our food. The Bovidae are very diverse in body size and horn shapes (Simpson 1949; Gentry 1968; Gentry 1992; Hassanin and Douzery 1999). The bovid horns are difficult to interpret. The available studies are old but are informative (Dürst 1926; Dove 1935) and summarized by Janis and Scott (1987) and by Davis et al. (2014). Preliminary data suggest that the horn is a composite of a separate ossification externally (the os cornu), a massive frontal origin horn internally and a keratinous horn sheath cover.

The other Pecora have not been studied as much. The Dromomerycidae are a group of taxa where the horns are constructed by an upward growth of the frontal bone. The dentitions are primitive (Frick 1937). Many confuse the Dromomerycidae with the Palaeomerycidae. Superficially they look similar. The horn of the first family is formed by the frontal whereas the horn of the second family is an ossicone. They both have an occipital horn (a convergence).

The Antilocapridae also have horns formed by the frontal bone but may be distinguished by the covering keratinous horn sheaths and more hypsodont teeth (O’Gara and Matson 1975). The keratinous sheath possesses laminae like in hooves. In Bovidae it does not. Antilocapridae shed one sheath and grow a second. Antilocapridae have been minimally studied.

The Giraffoidea (Climacoceridae, Palaeomerycidae and Giraffidae). The Giraffoidea of the Pecora is a group of taxa united by the presence of supraorbital ossicones. Ossicones are separate ossifications which later in life fuse with the frontal bone (Ganey et al. 1990; Solounias 2007; Solounias 2023).

There are three major studies of cladistic analyses for Giraffoidea (Hamilton 1978; Geraads 1986; and Ríos-Ibàñez et al. 2017).

### Climacoceridae and Palaeomerycidae

Climacoceridae is a difficult to study group from east Africa and now the Siwaliks of Pakistan (this paper) (Hamilton 1978). Presently Climacoceridae are assumed to have ossicones. This is based on two specimens. One form Moruorot Hill where the base is detached (Grossman and Solounias 2014 Fig. 2 – UCMP 40461). *Pachya* described and named here is one with a detached ossicone. The other specimen is from the Siwaliks (Y GSP 30608) and shows at the base a flat surface of a detached ossicone type as in the Moruorot one. Climacoceridae do not have occipital horns.

Palaeomerycidae have ossicones and an enigmatic occipital horn (Qiu et al. 1985; Astibia and Morales 2006; Sánchez et al. 2015). They have often been confused with Dromomerycidae. This is due to the occipital horn which is convergence. Some domesticated cows also have a occipital horn (a small one with no keratin but in the same median position).

Giraffidae possess ossicones only and are only large-sized taxa (Bohlin 1926; Hamilton 1978; Solounias 2007; 2023). No small-sized species exist unlike in Cervidae and Bovidae. They possess crenulated enamel, a canine with a second lobe, and a reduction of the opening of the nasolacrimal duct. The metapodials are large. A metapodial of the okapi is two times longer than that of *Taurotragus oryx* but the two species are of similar size.

We name several new genera and species. The material is very small and one may wonder why name new taxa with such a poor record. This is because we need to know the diversity of life and there are very few changes in the near future to obtain more material. The Climacoceridae and Palaeomerycidae are particularly rare. They may have inhabited highlands as thus they lived outside of the main flood plains where deposition of bones is common. On the other hand, Giraffidae are more abundant and were inhabiting lowland depositional environments. These three groups differentiate near the 18 Ma. There is no record then to document their original appearance. When they first appear, they are full-fledged in their three respective groups. The sediments that record the oldest taxa are Moruorot Hill, Kalodirr, Gebel Zelten, Zinda Pir and Kalmlial.

### Description of the eight subfamilies of Giraffidae

The Giraffidae form a number of subfamilies which are not directly connected. They share the bilobed canine, closed orbital opening for the nasolacrimal duct, crenulated enamel, ossicones and large body size large metapodials and browsing diet (except for the *Samotherium* group).

In Giraffidae the proximal end of the metatarsal can be used as a diagnostic tool as it differs in most species (Figure 38). The ossicones are also useful in separating the subfamilies.

The Giraffidae can be subdivided into subfamilies. These are:

### Progiraffinae new rank

Based on *Progiraffa exiqua* Pilgrim 1911. This includes *Progiraffa* (or *Canthumeryx sirtensis* and *Georgiomeryx georgalasi* and possibly other unnamed early Miocene species). There are undescribed ossicones of this taxon form Kalodirr which were similar to the okapi. It was found in Gebel Zelten (known as *Canthumeryx –* it differs from Zarafa which is a palaeomerycid) and Moruorot Hill (undescribed). In the Zinda Pir and lower Siwaliks, there are dental specimens. This earliest giraffid is *Progiraffa exigua* has been described by Barry at al. (2005). The sample in that description consists of mixed taxa but all of which are found in the same deposits. One horn is a hypsodontine bovid and a skull is a *Prolibytherium*. The material has been covered by Hamilton (1973 and 1978 as *Canthumeryx*). The cervicals of *Canthumeryx* are elongated. We know from these specimens that *Progiraffa* had a short conical ossicone and a longish cervical vertebrae. The premolars are clearly primitive with long crests and lingual valleys open (Hamilton 1978). I thank Dr. Ari Grossman for his help with the Kalodirr material. What makes these specimens giraffid is the presence of ossicones. Note that *Zarafa zelteni* from Gebel Zelten is excluded from these species as it is a palaeomerycid. In *Zarafa* the enamel is not crenulated; thus, it cannot be a giraffid. In addition, the morphology of the occipital is highly suggestive that it had an occipital horn which is broken off. Giraffidae do not have an occipital horn. There is an undescribed orbital ossicone of *Zarafa* in the NHM which clearly point towards this taxon being a palaeomerycid.

The *Giraffa* is close to *Progiraffa*. The true ossicone of the giraffe is short with a broad base. That is the ossicone after removing the external epikouron. The dentition is plesiomorphic. In these two features the *Giraffa* is similar to *Progiraffa*. Thus, in the evolution of the giraffe directly from *Progiraffa* is plausible. The giraffe added a frontal sinus and elongated the metapodials. The cervicals of course became longer. In conclusion, as far as the fossil record can help we have *Progiraffa* as the most likely direct ancestor.

### Giraffokerycinae Solounias 2007

Based on *Giraffokeryx punjabiensis*. *Giraffokeryx* punjabiensis from the Chinji of the Siwaliks and *G. primaevus* from Fort Ternan in Kenya (and possibly from the Chinji) are the two-known species. The second is the more primitive (*Palaeotragus* of Churcher 1970). They had four ossicones and no frontal sinuses. The posterior ossicones were long and deeply grooved and perhaps were covered with keratin. At the base, there is a bulge of disorganized bone. The anterior ossicones were fused at the base. They had long metapodials. *Libytherium* also has basal bulges on the ossicones. They have long metapodials. The metatarsal of *Giraffokeryx* has an untypical proximal digit III.

The short neck of the okapi is similar to *Bramatherium*. They also share the four ossicones and large anterior frontal sinuses (Solounias 2023). The second pair of ossicones of the okapi are reduced plates in pits above the orbits. The metapodials are also similar. Thus, the okapi does have affinities with *Bramatherium*. In *Giraffokeryx* and the okapi they share the four ossicones but the neck of *Giraffokeryx* and the metapodials are longer. Thus, *Giraffokeryx* is less similar.

### Okapiinae Bohlin 1926

Based on *Okapia johnstoni*. There is no fossil record of *Okapia*. The living species is a mystery with many unsolved issues. It is possible that there are fossil okapi-like animals in the Chinji Formation of the Siwaliks. We are currently researching these. *Okapia* possesses two posteriorly placed ossicones that are smooth-surfaced, conical and the bone is commonly exposed at the apices. There may be a second rudimentary pair of ossicones over the orbital margin. All the remains of these ossicones is a pit depression and sometimes a bony plate in the pit. The females are hornless but they may rarely have small ossicones. The females are larger-sized than the males. The ossicone area may produce periodically keratinous horn sheaths.

*Okapia* has well developed nasal and anterior frontal sinuses and inflated bullae. They have shortened necks. A compressed proximal metatarsal and cuneiforms fused to the cubonavicular. They have medium-sized metapodials.

A Miocene species from the Baringo Basin of Kenya; namely *Afrikanokeryx leakey*. It has bulbous bullae and canals in the ossicones. It is a rare and unique taxon. It may be a distant relative to *Okapia* (Harris et al. 2010).

*Ua pilbeami* from the Chinji is close to the okapi. It has distal constriction in the ossicone. It also has an anterior large frontal sinus (Solounias et al. 2022).

### Bramatheriinae new rank

Based on *Bramatherium perimense*. The following group is commonly termed Sivatheriinae but ironically, this name is not correct as we remove *Sivatherium* from the subfamily. The subfamily can be named Bramatheriinae as *Bramatherium perimense* is the more complete specimen and the oldest named genus in the extinct Giraffidae (Falconer and Cautley 1843). In *Bramatherium perimense* from Perim Island there is a skull, limbs and a complete mandible. *Bramatherium permimense* is a large giraffid found in the Perim Island and in Dhok Pathan. It has anterior bosses that were huge and commonly mistaken for ossicones. Although these are the anterior, they were situated posterior to the orbits. The bosses may had a thin veneer-like ossicone on them. The posterior ossicones were massive straight and directed laterally. Both ossicones share a common base. Within the ossicones, there are two canals. Thus, both the back and front ossicones were formed my merging of separate ossifications. The canals are separated by a think plate of bone. The neck, known from *Bramatherium megacephalum* at Yale Peabody Museum, was short and massive. The metatarsals (of *B. perimense*) display accessory digits and two well-developed ventral ridges. The medial right (digit II) is taller than the lateral. The median ridge of the metatarsals in very strong while the lateral ridge is very weak.

*Bramatherium* is also found in the Pliocene of Turkey and the Miocene of Samos (the oldest at 7.2 Ma). They have short to medium in length metapodials.

*Helladotherium duvernoyi* is very similar. It probably had small posterior disc shaped ossicones that have never been identified. The main ossicones were probably disc-shaped. The boss is a long oval. There is a possibility that the anterior ossicones have been found. NHM M37843 is one of these specimens. It is a flattened disc-like ossicone with a rich sinus at the base. The boss of the holotype skull is long and oval as was the adhering ossicone. The surface is rounded and full of vessel impressions and irregular islands of epikouron. It is similar to the anterior ossicones of *Libytherium maurusium* form Djibouti (Geraads 1985). There may have not been posterior ossicones as there are no bosses on the type skull. We place *Vishnutherium iravadicum* from Burma in *Helladotherium*. It is a rare large taxon with extremely large parastyles metastyles and cingula and it possesses deep troughed metapodials. The premolars resemble those of *Helladotherium*. This taxon has been found in Pikermi, Samos and Maragheh.

*Helladotherium grande* a skull figured by Pilgrim 1911. It is an occurrence of a Pikermian Biome taxon also present in the Siwaliks (Pilgrim 1911).

*Libytherium maurusium* was widespread in the Pliocene of Africa. *Libytherium proton* in the Siwaliks. Many consider it to be congeneric with *Sivatherium* on the overall superficial similarity of the ossicones and the large body size. The ossicones gently curve and are positioned posteriorly on the skull in both genera. The ossicones are twisted and deeply grooved. The anterior ossicones are flattened discs. We believe most of the African *Sivatherium* are probably *Libytherium* instead. *Birgerbohlinia* and *Decennatherium* are very similar to *Libytherium*.

*Bramiscus* is a small-sized taxon similar to *Bramatherium*. The median ridge of the metatarsals in very strong while the lateral ridge is very weak. The anterior ossicones are fused at the base (*Bramiscus micros* Ríos, Abbas, Khan & Solounias, 2023).

### Palaeotraginae Pilgrim 1911

Based on *Palaeotragus rouenii* from Pikermi. Palaeotraginae are represented by two species: *Palaeotragus rouenii* (smaller) and *Palaeotragus coelophrys* (larger); both are very similar species. A new genus (unnamed) *germanii* from Bou Hanifia is a little older and can be included. The stratigraphy and morphology suggest a progressive reduction in size from a larger species to smaller ones. Bou Hanifia is 12 Ma and Pikermi is 7.5 Ma. The ossicones of *germanii* are simple medium-sized and grooved-surfaced spikes positioned more internally than the orbital rim (Arambourg 1959). The apices are often bare bone. The females were hornless. The premolars were long. In *Palaeotragus* there is a gap between the external auditory meatus and the glenoid fossa that gap is filled with a bony shelf. They may specializations in their premaxillae suggesting a different type of upper lip. We know this from specimens from Gansu. The metapodials are long and slender.

### Samotheriinae Hamilton 1978

Based on *Samotherium boissieri* from Samos. It includes *Injanatherium hazimi*, *Samotherium*, *Schansitherium*, and *Alcicephalus*. They have long spike like ossicones that are supraorbital (posterior orbital margin). The ossicones are smooth surfaced and are often exposed at the apex. The fontal sinuses are small to absent. The premolars are reduced in size. There is a morphocline of increasing masseteric fossae and the displacement of the orbit posteriorly. The smallest masseter is in *Alcicephalus*, followed by *Samotherium boissieri* and *Schansitherium*. *Samotherium major* has an extremely large masseter and a posterior orbit reminiscent of Alcelaphini. *Samotherium boissieri* possessed a squared premaxilla for mixed feeding. *Alcicephalus* had a wide occiput the others had a narrow one. The females can be horned but not always. The necks are longish and the metapodials are short. *Injanatherium hazimi* Henitz et al. 1981 from Injana province Gebel Hamrin Iraq (Henitz et al. 1981). This may be congeneric with *Samotherium boissieri*. Samotheriinae are the only Giraffidae that possessed mixed and grazing diet. All other taxa of Giraffidae were browsers.

### Bohlininae Solounias 2007

Based on *Bohlinia attica*. This is an example of a taxon described by Gaudry on postcranial material only in 1862. The skull was described in 1926 by Bohlin. *Bohlinia* has massive ossicones without canals. Skull roof flat without sinuses. P3 and P4 with enlarged curved inward parastyles. Cervicals long. Metapodials long. Metapodial trough deep. Medial femoral epicondyle reduced.

*Bohlinia* possesses P2 and P3 which are very specialized. They have styles turned inwards The lingual surface is circular in occlusal view. *Bohlinia* has a long neck (we have two axis vertebrae and they are long like in *Giraffa*) but the C7 is not specialized (unpublished material in the Athens Paleontology collection). It is a long C7 but without ventral tubercles (where the giraffe is specialized with a C7 that is exceptional in having ventral tubercles). In other words, the C7 of *Bohlinia* is rather normal. What is untypical is the lack of a large and medial epicondyle on the femur. In this respect, *Bohlinia* is unlike other giraffids. The femur resembles more that of a dromedary. Thus, *Bohlinia* resembles a camel in limb posture and is rather different from the majority of the Giraffidae.

*Honanotherium* is recently known by complete skulls from Gansu (unpublished data). They are very similar to *Bohlinia* except of the upper premolars that are normal (less specialized). The metapodials were thicker and shorter than in *Bohlinia*. *Bohlinia* evolved in the Pikermian Biome. *Honanotherium* in the Beotian Biome. The metapodials are either medium or long in this subfamily. In summary, they differ from Giraffinae in the distal femur and the deep troughs of the metapodials, the C7 vertebra is a primitive and unspecialized. The massive ossicones and the absence of head sinuses are characteristic.

### Giraffinae Gray 1821

Based on *Giraffa camelopardalis*. *Giraffa* evolved in the Siwaliks of Pakistan. It migrated into Africa rather recently. The ossicones are covered in males with epikouron which extends down the face up to the face muscles. The parietals are not covered by epikouron. There is usually a third central ossicone in males. The frontal sinuses are strongly developed (Solounias 2023). The teeth are robust and the premolars are large. The cervicals are long. C5, C6 and C7 are close to homogeneous in shape. T1 resembles a C7. The metapodials are long and slender and the ventral troughs are minimal. *Giraffa jumae* from Olduvai has smaller frontal sinuses and not nasal ossicones. Thus, the modern giraffe is a rather recent species. *Giraffa punjabiensis* is as large as modern *Giraffa*. It is the oldest of the genus and apparently *Giraffa* is closely related to *Bohlinia*, it evolved in the Siwaliks of the Indian Subcontinent (Danowitz et al. 2017).

The name camelopardalis is ancient. The name means camel with spots – pardalos is with spots or silly. Like leopardalis which is a lion with spots. The Greeks would probably have a different name if they referred the species to the ones with square patches. These are what we have now. I think that they named it after *Giraffa sahara* which was living in Northern Africa and the Sahara. Sahara was 10,000 years ago a woodland with numerous lakes. There are many petroglyphs of this species which is now extinct (Solounias 2023). They show a giraffe with spots which is appropriate for the name *camelopardalis*. You can see some of these in Google images. The *Giraffa* with squares (four to eight species) is not ideal for the name *camelopardalis* and ancients may not have seen these giraffes.

### Sivatheriinae Bonaparte 1850

Based on *Sivatherium giganteum* which is the only species in the subfamily. It is difficult to interpret this taxon from the Pleistocene of India. It is large, the size of *Bubalus*. The posterior ossicones are composed of thousands of horizontal laminations. The laminations are millimeters thick and are covered by flat broad ribbons of bony material. The frontal sinuses are eight small chambers but are small in volume. Females without ossicones. *Sivatherium giganteum* Pliocene of the Siwaliks of India. It is unique because of the structure of the ossicones. They actually may not be ossicones but a unique type of horn. The internal anatomy of the “horn.” Lateral bumps are present on the edges of the twisted horn shaft structure. There are no internal canals. The sinuses of the skull are small. The anterior ossicones seem normal giraffid ossicones and are conical. The females were hornless (female skull NHM M 39523). The metacarpals are notably shorter than the metatarsals unlike other Giraffidae (Fig. 37 D and E). This morphology can be seen in some petroglyphs of *Sivatherium*. There are petroglyphs of the untypical species in Africa (Solounias 2023). The neck was short with massive cervical processes. The head may have been positioned down as in *Bison*. It was a mixed feeder and probably a browser (incisors NHM M 15288). They have short metapodials and the anterior are notably shorter like a *Hyemoschus*. *Sivatherium* may be a palaeomerycid. It is also possible that the back horns are occipital horns and, in that case, *Sivatherium* only has a single pair of ossicones.

### Relations of Giraffidae

Gradual changes are found in many examples of evolution. For example, changes in the mammalian ear in the origin of birds from species like *Archaeopteryx* and in numerous fish lineages where skulls and fins and body shapes can be organized in evolutionary sequences. We have good sequences in human and Cetacean evolution. You can read about sequences in many books (such as: E. Mayr 1988 p. 406 “Toward a new philosophy of biology;” G. G. Simpson 1944 “Tempo and mode of evolution).”

Alternatively, evolutionary relationships in various taxa can be with small gaps or large gaps. In addition, horizontal or vertical sequences are desired but not always available. At the phylum level, most phyla are distinct with very few intermediates. This is well-documented. In this case the gaps are large. The baupläne is key and recognizable by morphology this is true in the classification of phyla. At a smaller level, the Giraffidae are in a similar situation. There are subfamilies but no clear way of connecting these in a coherent evolutionary fashion. Gaps do exist.

## CONCLUSIONS

Lower Siwaliks Giraffoid faunas (18.3-13.1 Ma) are: The earliest giraffoids recorded are the climacoceratids *Vittoria*, *Goniomeryx flynni* and *Orangemeryx badgleyi,* and the giraffids *Progiraffa exigua, Ua pilbeami*, and *Orea leptia.* The earliest palaeomerycid of the Siwaliks record is *Goniomeryx flynni* followed by *Nuchalia gratia.* The earliest giraffid is *Progiraffa exigua* (it includes *Canthumeryx* and *Georgiomeryx*).

Middle Siwaliks Giraffoid faunas (13.1-10.1 Ma) are: The majority of plaeomerycids appear during the Middle Siwaliks, including *Lateralia morgani, Tauromeryx canteroi* and *Fovea fossata nov.* They coexisted with several giraffids including *Giraffokeryx punjabiensis, Giraffa punjabiensis, Lyra sherkana,* and *Bramiscus micros*.

Upper Siwalik Giraffoid faunas (10.1-0 Ma) are: The latest giraffoid faunas included the last palaeomerycids which disappeared right at the beginning of the Upper Siwaliks. The main representatives of the clade were the large Bramatheriinae *Bramatherium megacephalum, Libytherium proton. Sivatherium giganteum* end *Giraffa sivalensis* were also found at these levels.

### New taxa

(1) The Palaeomerycidae: *Tauromeryx canteroi* nov. sp., *Nuchalia gratia* nov. gen. sp. nov., *Fovea fossata* nov. gen. sp. nov, *Goniomeryx flynni* gen. nov. sp. nov., and *Lateralia morgani* nov. gen. nov. sp. nov.
(2) new Climacocerinae: *Vittoria soriae* gen nov. sp nov., *Orangemeryx badgleyi* nov. sp., *Pachya moruoroti* gen. nov. sp. nov.
(3) Preliminary systematics suggest that Giraffidae can be subdivided into two broad clades: the long and the short-neck groups. Short-neck are: *Giraffokeryx punjabiensis*, *Ua pilbeami*. *Decennatherium asiaticum*, *Bramatherium megacephalum*, *Bramatherium perminese*, *Bamiscus micros*, and *Sivatherium giganteum*. The okapi fits here. Longer neck are: *Progiraffa exigua*, *Orea leptia*, *Injanatherium hazimi*, *Giraffa punjabiensis, Giraffa sivalensis*, *Palaeotragus germanii* and *Bohlinia tungurensis. Giraffa camelopardalis* fits here.

### Neotypes

*Giraffokeryx punjabiensis* and *Giraffa punjabiensis*.

## ABBRIVIATIONS

AMNH: American Museum of Natural History, New York
GSI (IM): Indian Museum in Calcutta
MCZ: Museum of Comparative Zoology at Harvard, Cambridge
MNHNP: Museum national d’Histoire naturelle de Paris
NHM: Natural History Museum London
NHMBa: Natural History Museum of Basel
MNCN: Museo Nacional de ciencias naturales Madrid
UCMP: Berkeley Museum, YGSP, Yale-Geological Survey of Pakistan collections Islamabad
YPM: Yale Peabody Museum New Haven.

## Acknowledgments

We thank John Barry. Without his meticulous collecting and curation these specimens would be unknown to science. We also thank David Pilbeam for making the expeditions to the Siwaliks possible. We thank Catherine Badgley, Larry Flynn, and Michèle Morgan. We thank Mahmood Raza. Thanks to Ari Grossman. We thank the museums and curators of AMNH, MCZ, YPM, NHM, MNHNP, NHMBe, for specimens. We thank Mahnaz Tehrani, Melody Young, Julia Molnar, Meredith Taylor, Kelsi Hurdle, and Dan Gibbons.

## Studies related to the current publication

- Ríos Ibáñez, M. (2017) - Evolution and systematics of the late Miocene Spanish Giraffidae (Mammalia, Ruminantia, Pecora). Doctoral thesis. Universitat de Valencia. URI: http://hdl.handle.net/10550/57778
- Ríos, M., Danowitz M. and Solounias N., (2016) - First comprehensive morphological analysis on the metapodials of Giraffidae. *Palaeontologia electronica*, 19(3), pp.1-39.
- Rios M., Danowitz M., Solounias N. 2019. First identification of *Decennatherium* Crusafont, 1952 (Mammalia, Ruminantia, Pecora) in the Siwaliks of Pakistan. *Geobios* 2019.10.007
- Ríos, M., Sánchez, I. M. & Morales, J. 2017. A new giraffid (Mammalia, Ruminantia, Pecora) from the late Miocene of Spain, and the evolution of the sivathere-samothere lineage. *PLoS ONE*, **<u>12</u>**(<u>11</u>), e0185378. doi:10.1371/journal.pone.0185378
- Ríos, M. & Morales, J. 2019. A new skull of Decennatherium rex Ríos, Sánchez and Morales, 2017 from Batallones-4 (upper Vallesian, MN10, Madrid, Spain). Palaeontologia Electronica, 22(2), PVC_1. doi:10.26879/965
- Ríos, M., Abbas, S. G., Khan, M. A., & Solounias, N. (2022). Distinction of *Sivatherium* from *Libytherium* and a new species of Libytherium (Giraffidae, Ruminantia, Mammalia) from the Siwaliks of Pakistan (Miocene). *Geobios*, *74*, 67-76.
- Ríos, M, Solounias N. (2023) - *Lyra sherkhana* gen. nov. sp. nov., a new genus and species of giraffid from the Miocene of the Siwaliks (Pakistan), Journal of Vertebrate Paleontology Ms number: JVP-2023-0014 (under revision final: just figs and appendices)
- Ríos, M, Solounias N., Smith S. Orea leptia: a new genus and species with the most slender metatarsal of any ruminant (Giraffidae; Mammalia). Need reference: we need to figure where this one is submitted or resubmit it, we have everything ready
- Ríos-Ibàñez M. Sayyed Ghyour Abbas2, Muhammad Akbar Khan2, and Nikos Solounias3 A new giraffid Bramiscus micros nov. gen. nov. sp. (Ruminantia, Giraffidae) from the Miocene of northern Pakistan. Paleontologia electronica.) PE MS No: 1243 (under revision almost final (R2).

### Also

Ríos, M., Sánchez, I.M. and Morales, J., 2017. A new giraffid (Mammalia, Ruminantia, Pecora) from the late Miocene of Spain, and the evolution of the sivathere-samothere lineage. *PLoS One*, *12*(11), p.e0185378.

Ríos-Ibàñez M. Sayyed Ghyour Abbas^2^, Muhammad Akbar Khan^2^, and Nikos Solounias^3^ – 2022. Distinction of *Sivatherium* from *Libytherium* and a new species of *Libytherium* (Giraffidae, Ruminantia, Mammalia) from the Siwaliks of Pakistan Need reference

### How to cite this study

Solounias, N. and Ríos Ibáñez M. 2024. Giraffoids from the Siwaliks of Pakistan. bioRxiv 40 pp. and 38 plates.

## Notes

### Competing Interest Statement

The authors have declared no competing interest.

### Summary of Updates

Correction of a genus and addition of a genus

